# RiboGRAM learns multi-scale non-coding RNA grammar for structural and regulatory inference

**DOI:** 10.64898/2026.10.12.758266

**Authors:** Hui Wang, Zhaoming Chen, He Huang, Shengdi Li, Andres Jäschke, Qiwei Ye, Yaqing Zhang

## Abstract

Protein-coding sequences are read through the triplet genetic code, whereas the organization of molecular information within non-coding RNA sequences remains elusive. Here, we identify a non-coding RNA grammar defined by multi-scale motif usage and 5’-to-3’ positional gradients, and develop RiboGRAM to learn this organization. Across nine organisms, non-coding RNAs showed scale-dependent divergence from coding sequences, with directional U-rich and AU-rich gradients. Integrating multi-scale sequence encoding with contextual modeling, RiboGRAM organized RNA families, revealing motif-associated and pair-like patterns across layers. Without structural supervision, non-coding RNA pretraining improved base-pair recovery over coding or messenger RNA pretraining, with representations supporting alignment-free diffusion-based 3D modeling of short RNAs. In human RNA–protein binding profiles, RiboGRAM identified position-resolved motif organization across protein targets and cell types, with sequence-preference shifts associated with reduced cross-cell transferability. Together, these results establish a learnable grammar of non-coding RNA sequence organization for structural and regulatory inference.

## Introduction

Understanding how sequence shapes the molecular properties of RNA is a fundamental goal of RNA biology. Protein-coding sequences are interpreted through the triplet genetic code within a defined reading frame^1^, whereas non-coding RNAs (ncRNAs) function as RNA molecules without a known universal sequence unit^2^. Although conserved motifs^3, 4^, compositional biases^5, 6^ and structural elements^7^ have largely been characterized within specific RNA families or biological contexts, how these features are organized across sequence scales and positions is still poorly understood.

Nucleotide language models have recently emerged as powerful approaches for learning sequence patterns that can be applied across diverse biological tasks^8–17^. RNA-FM^13^ and RiNALMo^10^ exemplify this paradigm by leveraging millions of ncRNA sequences to learn representations that support RNA structural and functional prediction. Recent models have further incorporated biological priors from motif-and RNA-type-aware objectives^12^, evolutionary relationships and transcript isoforms^18^. However, despite advances in predictive performance, the biological information encoded within the RNA representation remains less well characterized. Establishing the transferability of this biological information across RNA structure and molecular interaction contexts remains a major challenge.

Here, we first identify systematic organization of ncRNA sequence across multiple scales and along the 5′-to-3′ axis in nine species, which we term ncRNA grammar. This model-independent characterization informed the design of RiboGRAM, which combines multi-scale local encoding with contextual sequence modeling. Layer-resolved analyses showed a hierarchy of biological signals, with RNA-family information becoming more apparent, motif-associated signals emerging in intermediate layers and pair-like patterns appearing later. Base-pairing information was markedly more recoverable with ncRNA-rich than with coding sequences (CDS) or messenger RNA (mRNA) pretraining in architecture-matched models. The learned representations further supported pairing-state inference across unseen RNA families and multiple sequence alignment (MSA)-free diffusion-based 3D modeling of short RNAs. Post-training on human RNA-binding protein (RBP) profiles extended the model to endogenous interaction contexts, where RBP-associated motif organization was mapped across cell types. These analyses support a learnable ncRNA grammar of primary sequence organization, with RiboGRAM capturing RNA family organization and structural information while extending to diverse molecular interaction contexts.

## Results

### Multi-scale ncRNA grammar guides RiboGRAM design

To examine how ncRNA sequence organization changes with sequence scale across species, we profiled ncRNAs across nine model organisms, from *E. coli* to human, at nucleotide, triplet and longer k-mer scales. Species-specific ncRNA profiles became increasingly distinct at longer sequence scales, with the mean pairwise profile distance increasing from 0.375 for 1-mers to 0.643 for 9-mers (**Fig. 1a** and **Supplementary Fig. 1a**). This scale dependence differed from that of coding sequences. Relative to CDS, ncRNA profiles showed greater cross-species similarity at the nucleotide scale, comparable distances near the triplet scale and greater divergence at longer scales (ΔD =-0.35 at 1-mer and 0.19 at 9-mer; **Fig. 1b**). A similar scale-dependent pattern was observed relative to mRNA, whereas mRNA and CDS remained closely matched across sequence scales (**Supplementary Fig. 1b)**. These comparisons indicate that ncRNA primary sequence organization differs from that of CDS and mRNA in a scale-dependent manner.

**Figure 1.**
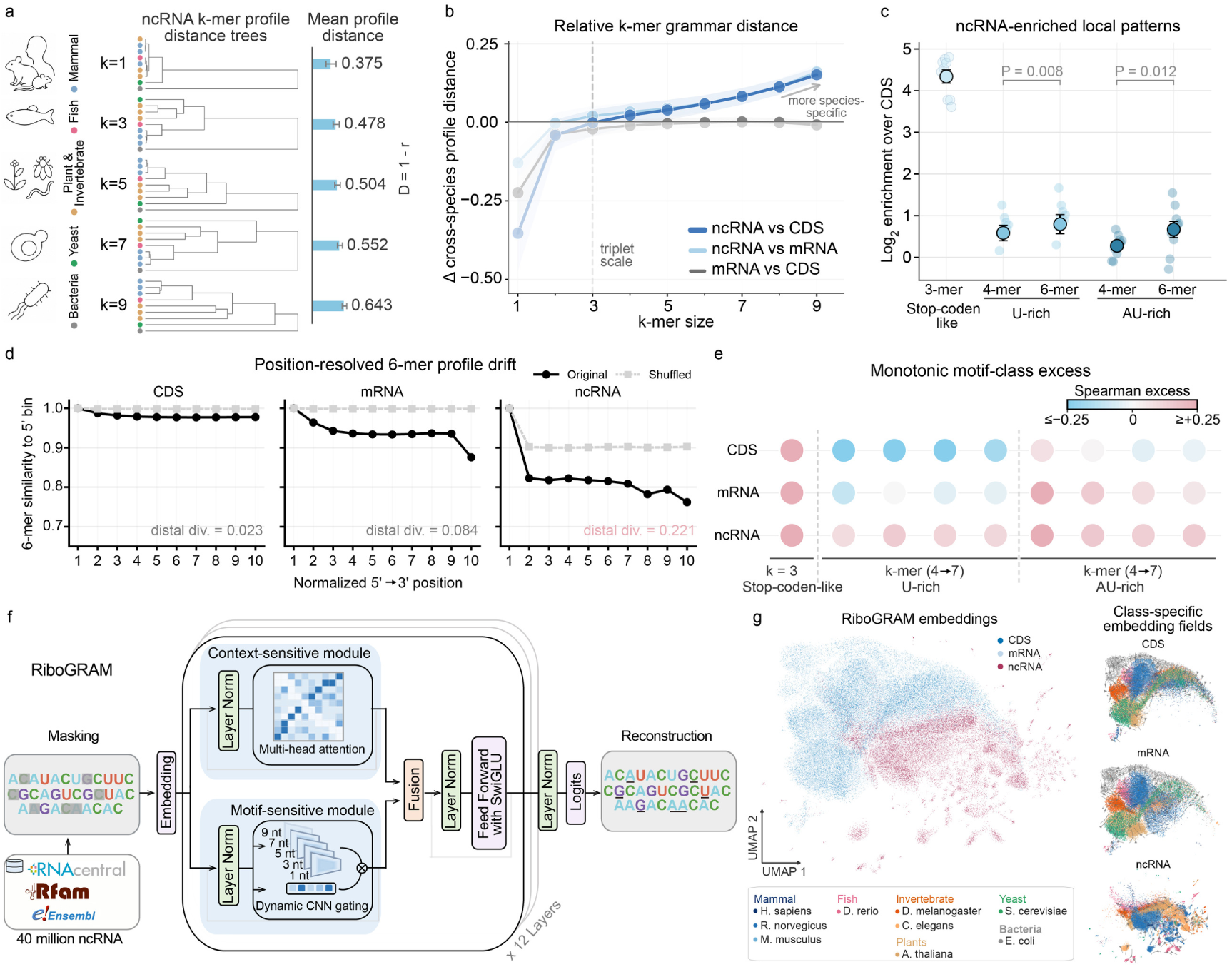
Multi-scale ncRNA grammar guides RiboGRAM design. a, Scale-resolved ncRNA k-mer profile distance across nine model organisms. For each species and k-mer size, ncRNA k-mer counts were normalized as log2(CPM + 1), and pairwise profile distance was defined as 1 − Pearson r. Trees show hierarchical clustering at increasing k-mer scales. Right, mean pairwise profile distance; error bars show standard error of the mean (s.e.m.) across species pairs. **b,** Relative k-mer grammar distance between RNA types. Lines show paired differences in species-pair profile distance between ncRNA, mRNA and CDS, using the distance metric in panel a. Positive values indicate larger cross-species profile distance for the first RNA class. Shaded bands show s.e.m. across species pairs. **c,** ncRNA-enriched local sequence patterns. Selected motif classes are shown as log2 enrichment in ncRNA relative to same-species CDS. Dots show species-level values and black circles show cross-species means. n = 9. P values are from paired Wilcoxon signed-rank tests across species. **d,** Position-resolved 6-mer profile drift. Profiles show similarity to the 5′-proximal bin across ten normalized 5’-to-3′ positions, with original sequences in black and sequence-shuffled controls in grey. Distal divergence summarizes the original–shuffled difference in 5′-to-distal profile drift, with distal similarity averaged over bins 8–10. **e,** Monotonic motif-class excess from nine model organisms. Circle color shows species-averaged motif-class Spearman excess for each RNA type, computed as the motif-class positional Spearman ρ minus the median ρ of composition-matched non-class k-mers. Positive values indicate stronger 5’-to-3’ monotonic enrichment than the matched background. **f,** RiboGRAM pre-training architecture. RiboGRAM was pretrained on approximately 40 million ncRNA sequences using masked sequence reconstruction and integrates a context-sensitive attention module with motif-sensitive convolutional modules including 1-, 3-, 5-, 7-and 9-nt kernels. **g,** UMAP projection of sequence-level RiboGRAM embeddings for CDS (n = 44,239), mRNA (n = 44,357) and ncRNA (n = 35,145) sequences from nine model organisms. Points are colored by RNA type. Right, RNA-type-specific local embedding fields summarize neighborhood structure in the same representation space.

We next identified the local sequence features that distinguished ncRNAs from CDS. At the triplet scale, UAG, UGA and UAA showed the highest enrichment in ncRNAs relative to in-frame CDS codons, with stop codons restricted to CDS 3’ ends (mean log2 enrichment = 4.65; **Fig. 1c**). Alternative triplet-counting schemes yielded different enrichment profiles, with UAG and UAA, but not UGA remaining among the most enriched when CDS and mRNA triplets were counted at all positions (**Supplementary Fig. 1c-f**). At longer scales, enrichment of U-rich and AU-rich local patterns relative to CDS increased from 4-mers to 6-mers (U-rich, 0.59 to 0.80; AU-rich, 0.28 to 0.67; **Fig. 1c**). These patterns were also enriched relative to mRNA, although with smaller effect sizes (**Supplementary Fig. 1g**). Across 4-to 9-mer scales, U-rich, AU-rich and dinucleotide-repeat-like pattern classes showed recurrent enrichment, with generally stronger enrichment relative to CDS than to mRNA (**Supplementary Fig. 1h,i**).

These enrichment analyses identified sequence features that distinguish ncRNAs from CDS, but did not resolve their positional organization. To assess this organization, we divided sequences of at least 60 nucleotides (nt) into ten length-normalized bins from 5′ to 3′ and compared each bin’s 6-mer profile with the 5’-proximal bin. CDS and mRNA profiles remained comparatively stable, whereas ncRNA profiles showed greater positional drift, with distal divergence of 0.221 compared with 0.023 for CDS and 0.084 for mRNA (**Fig. 1d**). Sequence-shuffled controls reduced this drift, supporting its dependence on sequence order. This positional drift persisted across k-mer sizes, with ncRNA profiles showing steeper declines and stronger monotonic trends at longer sequence scales (**Supplementary Fig. 2a,b**). Position-resolved motif-excess analyses further showed stronger 5’-to-3’ monotonic increase for U-rich and AU-rich motif classes than for composition-matched background k-mers across 4-to 7-mer scales, with the strongest ncRNA–CDS contrasts for U-rich 4–6-mers and AU-rich 4-mers (Δ = 0.380–0.482; **Fig. 1e** and **Supplementary Fig. 2c**). Stop-codon-like triplets also showed positive 5’-to-3’ monotonic excess, but this trend was shared across RNA types rather than specific to ncRNAs. The greater positional divergence of ncRNA profiles remained evident when same-species CDS profiles were used as an alternative reference (**Supplementary Fig. 2d**). These findings reveal multi-scale motif usage and directional 5’-to-3’ sequence organization in ncRNAs.

We next asked whether this multi-scale sequence organization could distinguish RNA types across species. When evaluated on held-out species, integrating sequence features across 1-, 3-, 5-, 7-and 9-mer scales improved balanced accuracy over triplet features alone, both for binary classification of ncRNA versus a pooled CDS/mRNA class (0.728 versus 0.698) and for three-class classification of CDS, mRNA and ncRNA (0.540 versus 0.511; **Supplementary Fig. 3a,b**).

Motivated by this sequence organization, we developed RiboGRAM, a self-supervised RNA representation model designed to integrate local patterns across multiple scales with broader sequence context. The model was pretrained on approximately 40 million ncRNA sequences assembled from RNAcentral^19^, Rfam^20^, and Ensembl^21^. Its architecture combines self-attention for contextual sequence modeling with parallel convolutional branches using 1-, 3-, 5-, 7-and 9-nt kernels to capture local patterns across multiple scales. The resulting representations are fused for masked sequence reconstruction, in which masked nucleotides are predicted from the remaining sequence context (**Fig. 1f**).

Ablation analyses showed that both multi-scale local encoding and contextual modeling contributed to RiboGRAM performance, with the full architecture yielding the lowest reconstruction loss and highest RNA-type classification performance across nine species (**Supplementary Fig. 3c-e**). UMAP projections of RiboGRAM embeddings further illustrated the separation of ncRNAs from CDS and mRNA across species (**Fig. 1g**).

### RiboGRAM organizes ncRNA families through layer-resolved representations

We next examined how ncRNA class organization emerged across RiboGRAM layers. UMAP projections showed clearer separation of the 13 ncRNA classes in intermediate-to-late layers than in early layers (**Fig. 2a**). Mapping sequence length and nucleotide composition onto the same projections revealed heterogeneous distributions across the embedding space (**Supplementary Fig. 4a**). Average product correction (APC)^22^-adjusted self-attention profiles varied across layers in their positional concentration and nucleotide-specific patterns (**Supplementary Fig. 4b-d**). In representative RNAs, layer-7 attention concentrated around annotated conserved elements across several ncRNA classes, with pair-like patterns in later layers (**Fig. 2a** and **Supplementary Fig. 5a,b**). These layer-wise changes in class organization were quantified by linear probes, with macro-F1 increasing from 0.730 at layer 1 to 0.978 at the final layer and a marked rise near layer-7 (**Fig. 2b** and **Supplementary Fig. 6a**). These analyses show that ncRNA class information becomes increasingly accessible across RiboGRAM layers.

**Figure 2.**
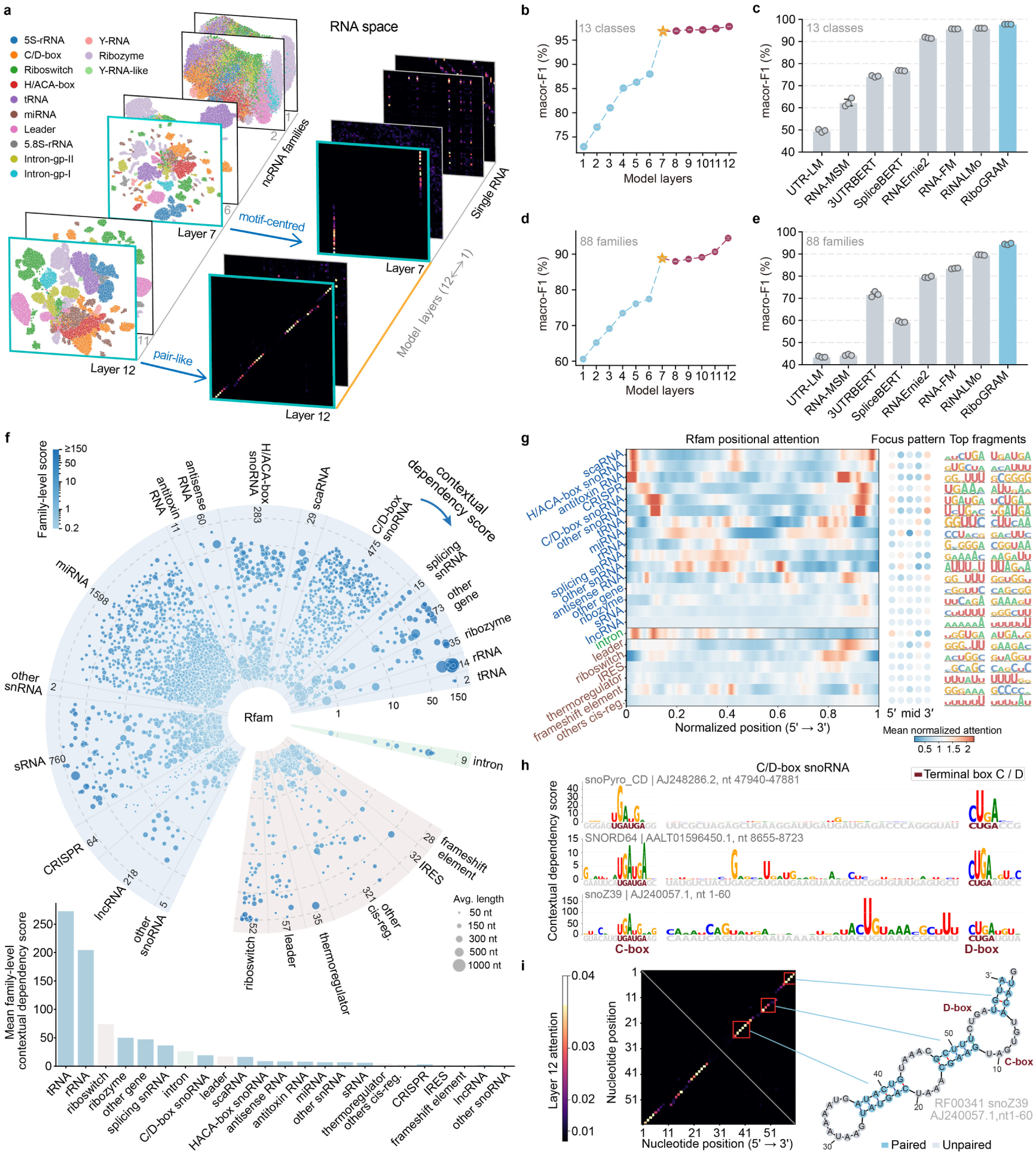
Layer-resolved RiboGRAM representations capture ncRNA class, family and motif organization. a, Layer-resolved embedding and attention organization in RiboGRAM. Left, UMAP projections of sequence-level embeddings for all 45,447 sequences in the 13-class NCYPred dataset23 at representative RiboGRAM layers. Right, head-averaged APC-adjusted self-attention maps for the representative C/D box snoRNA snoZ39 at the same layers, which show motif-centered and pair-like attention patterns. **b,** Layer-wise linear probing classification performance on the held-out NCYPred test set23 comprising 13,646 sequences. Points and error bars indicate mean ± standard deviation (s.d.) macro-F1 across three random seeds (n = 3). **c,** Final-layer performance of RiboGRAM and representative RNA language models on the same test set as in panel b. Error bars indicate mean ± s.d. macro-F1. n = 3. **d,** Layer-wise linear-probing performance on the held-out 88-family Rfam test set comprising 25,342 sequences53. Error bars indicate mean ± s.d. macro-F1. n = 3. **e,** Final-layer performance of RiboGRAM and representative RNA language models on the same test set as in panel d. Error bars indicate mean ± s.d. macro-F1. n = 3. **f,** Distribution of contextual dependency across all 4,178 families in Rfam 15.020. Family-level contextual dependency scores were calculated from the corresponding Rfam seed sequences. Each bubble represents one Rfam family, grouped by ncRNA subtype within the gene, cis-regulator (’cis-reg.’) and intron classes. Radial distance and bubble color denote the family-level contextual dependency score on a logarithmic scale, and bubble size indicates mean sequence length. Bottom, mean family-level contextual dependency score for each ncRNA subtype. **g,** Position-resolved contextual dependency and associated sequence motifs across ncRNA types shown in panel f. Left, mean contextual-dependency profiles along the length-normalized 5’-to-3’ coordinate, with ncRNA subtypes grouped according to the Rfam annotation hierarchy. Middle, dot summaries of regional focus across the same coordinate. Right, the two top-ranked hexamer motifs from contextual dependency enrichment for each ncRNA subtype. **h,** Nucleotide-level contextual dependency profiles for three representative C/D box snoRNAs, snoPyro_CD, SNORD64 and snoZ39. Letter height represents the position-specific contextual dependency score; the conserved C-box (UGAUGA)24 and D-box (CUGA)24 are highlighted. **i,** Comparison of pair-like signals in the layer-12 APC-adjusted self-attention map for the C/D box snoRNA snoZ39 with its Rfam-annotated secondary structure.

Using the same linear-probing framework, we compared RiboGRAM representations with those of representative RNA language models at both class and family levels. On the 13-class NCYPred benchmark^23^, several models achieved high performance, with RiboGRAM reaching macro-F1 of 0.978 compared with 0.957 for RiNALMo^10^, the strongest baseline (**Fig. 2c** and **Supplementary Fig. 6b)**. The difference was greater on the finer-grained 88-family Rfam benchmark, where RiboGRAM reached a macro-F1 of 0.945 compared with 0.896 for RiNALMo (**Fig. 2d,e** and **Supplementary Fig. 6c,d**).

Having established that RiboGRAM representations distinguish ncRNA classes and families, we next asked whether the model also captured sequence constraints at individual nucleotide positions. We quantified this property using contextual dependency scores, with higher scores indicating more confident recovery of a masked nucleotide from its surrounding sequence context. Across all 4,178 RNA families in Rfam 15.0, family-level contextual dependency varied systematically among ncRNA subtypes. Subtype-level mean scores were higher for ribosomal RNA (rRNA), transfer RNA (tRNA), riboswitch and small nucleolar RNA (snoRNA)-related subtypes, whereas several *cis*-regulatory subtypes, including internal ribosome entry site (IRES) and frameshift elements, showed lower scores (**Fig. 2f**). The highly conserved 5S rRNA family provided a focused case of within-family organization, in which RiboGRAM resolved sequence variation associated with taxonomic lineage and cellular compartment within a shared structural scaffold. Candidate plant 5S-like sequences mapped near annotated photosynthetic organellar 5S rRNAs in RiboGRAM space, with greater similarity to organellar 5S rRNAs and plastid-like flanking genomic context. In addition, mutation scores obtained without task-specific training captured part of the variation in 60S subunit incorporation among human 5S rRNA variants (**Supplementary Note 1**).

We next mapped contextual dependency along RNA sequences and examined local sequence features associated with high-scoring positions. Across Rfam annotation classes, gene-associated subtypes showed broadly symmetric profiles, whereas intron-associated and *cis*-regulatory subtypes showed greater contextual dependency toward the 5′ and 3′ ends, respectively (**Fig. 2g**). In C/D-box snoRNAs, high-scoring positions mapped to the conserved C and D boxes (UGAUGA and CUGA, respectively)^24^ (**Fig. 2h**). Elevated scores were also observed at the H and ACA boxes of H/ACA-box snoRNAs and within T-loop regions of tRNAs, whereas selected long non-coding RNA (lncRNA) examples showed high scores within low-complexity A-or U-rich tracts (**Supplementary Fig. 6e-g**). Complementing these position-wise profiles, the layer-12 APC-adjusted self-attention map for the C/D-box snoRNA snoZ39 contained pair-like patterns aligned with base-paired regions in its Rfam-annotated secondary structure^20^ (**Fig. 2i**). Analogous pair-like patterns were observed in representative tRNAs and H/ACA-box snoRNAs (**Supplementary Fig. 6h,i**).

### RiboGRAM representations support transferable inference of RNA structure

We therefore asked whether base-pairing information could be recovered directly from self-attention. For each language model, an attention head and pairing threshold were selected on the bpRNA-1m VL0 validation set^25, 26^ and fixed for held-out evaluation (**Supplementary Fig. 7a-c**). Base-pair recovery from self-attention varied markedly across pretraining corpora. Models pretrained on untranslated region (UTR), CDS or mRNA sequences showed only weak recovery of annotated base pairs (pair-level micro-F1 ≤ 0.9%), whereas models pretrained on ncRNA-rich corpora showed substantially greater recovery (micro-F1 = 31.7–58.2%; **Fig. 3a** and **Supplementary Fig. 7b-d**). RiboGRAM reached a micro-F1 of 58.2%, compared with 55.6% for the structure-supervised RNet-SS^27^ and 51.1% for RiNALMo^10^. Architecture-matched CDS-and mRNA-pretrained controls, trained using the same procedure as RiboGRAM, reached micro-F1 scores of 0.4% and 0.5%, respectively, further supporting the contribution of ncRNA-rich pretraining to the recoverable pairing signal.

**Figure 3.**
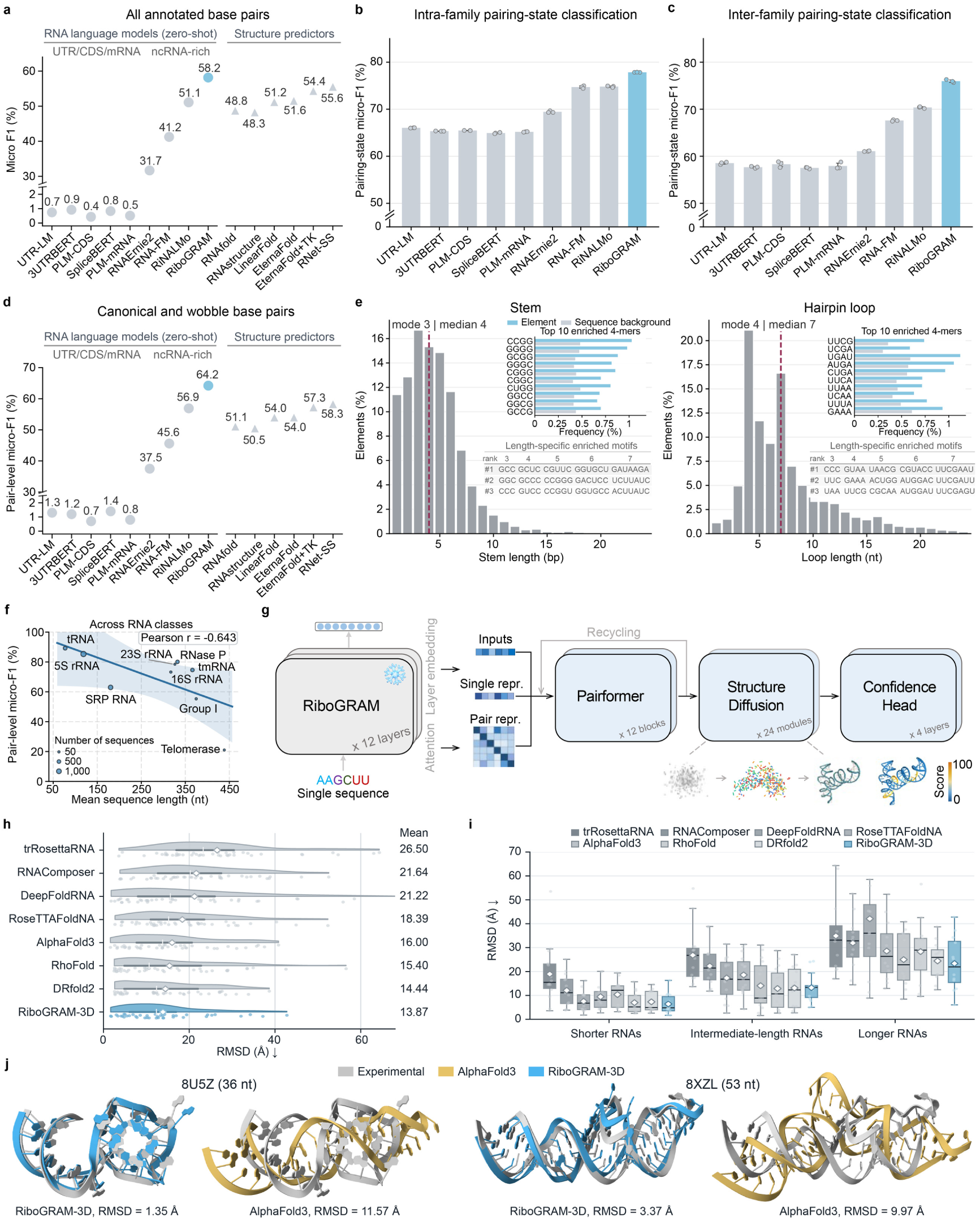
RiboGRAM representations support transferable RNA structure inference. a, Zero-shot recovery of all annotated base pairs on the bpRNA-1m TR0 set25, 26 (n = 10,814 sequences). RNA language models are grouped by pre-training corpus, compared with secondary structure predictors. EternaFold+TK denotes EternaFold with ThreshKnot decoding59, 60 and RNet-SS is structure-supervised27, 61. For each language model, the attention heads and decision thresholds were selected on VL025 and fixed before TR025 evaluation. **b,** Intra-family probing of nucleotide pairing states using frozen nucleotide representations. MLP probes were trained on TR0, selected on VL0 and evaluated on TS0. Bars show mean ± s.d., with n = 3. **c,** Inter-family probing of nucleotide pairing states using frozen nucleotide representations. MLP probes were trained and selected on TrainSetA28 and evaluated on TestSetB28. Bars indicate mean ± s.d., with n = 3. **d,** Recovery of canonical and wobble base pairs (A–U, G–C and G–U) on TR0 using the same VL0-selected attention head and decision threshold in panel a. **e,** Length distribution and local sequence patterns of annotated stems and hairpin loops in TR0. Histograms show the proportions of structural elements at each length, with mode and median marked. Insets show the ten most enriched 4-mers in each element type and compare their frequencies within the element with those across the corresponding full-length sequences. Lower tables show the top three highest-ranked enriched motifs at the indicated element length. **f,** Class-level RiboGRAM zero-shot pair-level micro-F1 across nine ArchiveII RNA classes with different mean sequence length, comprising 3,865 sequences29, 30. Points denote RNA classes, and point area indicates the number of sequences in each class. The line shows an ordinary least-squares fit, with the shaded band indicating the 95% confidence interval. **g,** Architecture of MSA-free RiboGRAM-3D model. Frozen RiboGRAM nucleotide embeddings are used to construct single and pair representations, with APC-adjusted attention maps incorporated as additional pairwise features. The resulting representations are processed through Pairformer31 blocks with recycling prior to structure-diffusion modules31 and a confidence head. **h,** RMSD distributions for eight tertiary-structure prediction methods across 41 held-out RNA targets. Points denote individual targets. Horizontal bars, vertical ticks and diamonds indicate interquartile range, medians and means, respectively. Mean RMSD values are listed at right. **i,** RMSD distributions for the same eight methods across the 41 held-out RNA targets stratified by length: shorter (27–55 nt; n = 14), intermediate-length (57–118 nt; n = 15) and longer (134–363 nt; n = 12) RNAs. Boxes show median and interquartile ranges, whiskers extend to 1.5 times the interquartile range, points denote individual targets and diamonds indicate means. **j,** Structural superpositions for the representative targets 8U5Z (36 nt) and 8XZL (53 nt). Grey denotes the experimental structure, blue for the RiboGRAM-3D prediction and ochre for the AlphaFold 331 prediction. RMSD to the experimental structure is indicated for each prediction.

We also tested whether frozen nucleotide representations carried information about nucleotide pairing state (paired versus unpaired), using identically configured multilayer perceptron (MLP) probes. In the intra-family evaluation, RiboGRAM reached a micro-F1 of approximately 78% (**Fig. 3b** and **Supplementary Fig. 7e**). For cross-family evaluation, the probes were trained on eight RNA families and evaluated on 22 held-out families^28^. RiboGRAM retained a micro-F1 of approximately 76%, compared with approximately 58% for the architecture-matched PLM-mRNA control (**Fig. 3c** and **Supplementary Fig. 7f**). Thus, pairing-state information remained accessible from frozen representations across held-out RNA families.

Base-pair recovery from self-attention was highest for canonical and wobble pairs (A–U, G–C and G–U), with RiboGRAM reaching a pair-level micro-F1 of 64.2% (**Fig. 3d**). Recovery was lower for non-canonical pairs and pseudoknot-associated crossing pairs, but the separation between ncRNA-rich and coding-or mRNA-pretrained models persisted, with RiboGRAM exhibiting micro-F1 scores of 19.6% and 39.5%, respectively (**Supplementary Fig. 7g,h**). Across individual RNAs, recovery also varied with global GC content and local GC heterogeneity (**Supplementary Note 2**).

We next examined local and positional sequence organization across annotated structural elements. Structural elements differed in both length distributions and local motif composition, with stems enriched for G/C-rich motifs, hairpin loops for UUCG, and multiloops and exterior regions for AU-rich motifs (**Fig. 3e** and **Supplementary Fig. 8a**). Position-resolved enrichment profiles also varied across structural-element boundaries and along the 5’-to-3’ axis within elements (**Supplementary Fig. 8b,c**).

To assess whether base-pair recovery was retained across diverse RNA classes, we applied the attention head and pairing threshold selected for each model, without recalibration, to nine ArchiveII classes^29, 30^ spanning a broad range of sequence length and base-pairing architectures. RiboGRAM achieved the highest mean class-level micro-F1 (69%), compared with 68% for the structure-supervised RNet-SS and 67% for RiNALMo (**Supplementary Fig. 9a**). Base-pair recovery varied substantially across classes (**Supplementary Note 3**) and was inversely associated with mean sequence length (Pearson r =-0.645; **Fig. 3f** and **Supplementary Fig. 9b**) with lower recovery among longer RNA classes, including telomerase RNA and group I introns (**Supplementary Fig. 9c**).

Building on the structural information recoverable from RiboGRAM representations, we developed RiboGRAM-3D, an MSA-free tertiary-structure prediction model that integrates nucleotide representations and self-attention features from the frozen encoder (**Fig. 3g**). The resulting single and pair representations were refined by Pairformer blocks^31^ with recycling before diffusion-based generation of all-atom coordinates.

We evaluated RiboGRAM-3D on 41 experimentally determined RNA structures from the PDB^32^, held out by deposition date with sequence-similar RNAs removed from the training set (see Methods). RiboGRAM-3D achieved the lowest mean RMSD (13.87 Å) among eight tertiary structure prediction methods, compared with 14.44 Å for DRfold2^33^ (**Fig. 3h**). Mean TM-scores were similar among RiboGRAM-3D, DRfold2^33^ and AlphaFold 3^31^ (0.350, 0.350 and 0.347, respectively; **Supplementary Fig. 10a**). Across the 41 targets, RiboGRAM-3D had lower RMSD than AlphaFold 3 for 29 targets and DRfold2 for 28, and remained among the lower-RMSD methods across targets stratified by detected homologs (**Supplementary Fig. 10b,c**). We then examined the relationship between attention-derived base-pair recovery and 3D prediction accuracy. Higher attention-derived base-pair F1 was associated with lower RiboGRAM-3D RMSD (Pearson r =-0.539; **Supplementary Fig. 10d**), although substantial dispersion indicated that pair recovery alone did not fully account for variation in tertiary structure accuracy. Across methods, longer RNAs showed higher RMSD (**Fig. 3i**). Representative RNAs further illustrated target-level variation in tertiary structure accuracy and attention-derived pair recovery (**Fig. 3j** and **Supplementary Fig. 10e-h**). Together, these results extend the structural utility of RiboGRAM representations from base-pair recovery to MSA-free 3D modeling, while longer RNAs remained more challenging in both tasks.

### RiboGRAM captures positional sequence organization in human RNA–protein interactions

We next examined whether RiboGRAM representations could resolve sequence organization in endogenous RNA–protein interaction profiles. To this end, RiboGRAM was post-trained on K562 RNA–RBP pairs from the HDRNet dataset using a joint objective that combined masked language modeling with binary interaction classification, yielding RiboGRAM-RPI (Fig. 4a).

**Figure 4.**
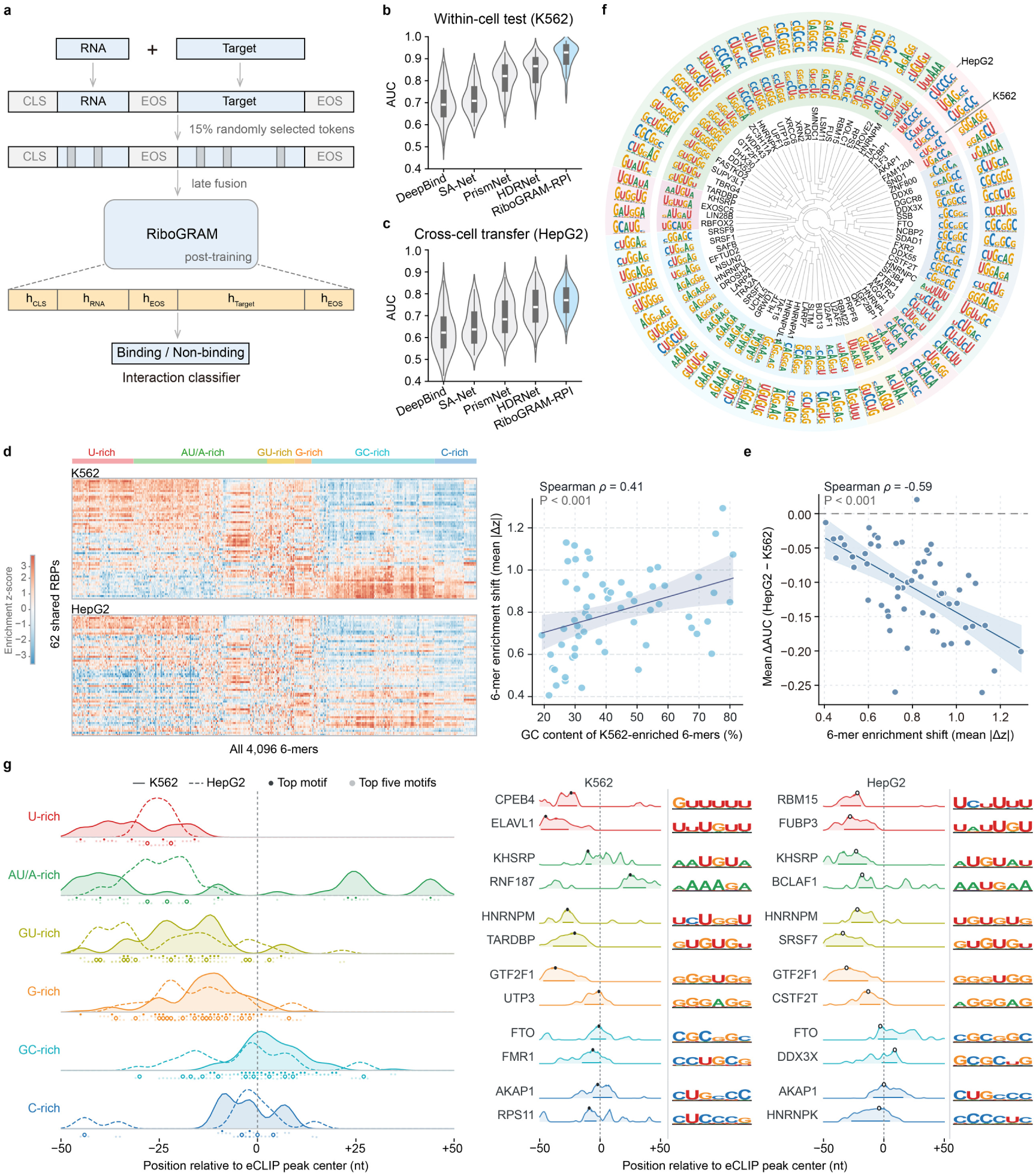
RiboGRAM captures positional sequence organization. a, Schematic of the unified post-training framework for RNA–target interaction prediction. RNA and target sequences are separated by special tokens, with 15% of tokens randomly selected for masking. RiboGRAM jointly optimizes masked-token reconstruction and binary interaction classification. **b,c,** RBP-level AUC distributions for DeepBind76, SA-Net77, PrismNet41, HDRNet34 and RiboGRAM-RPI in the within-cell K562 test across 112 RBPs (**b**) and during cross-cell transfer to HepG2 across 62 shared RBPs (**c**). The K562-trained models were applied to HepG2 without cell-line-specific retraining. Violin widths show kernel-density estimates, and internal box plots indicate medians and interquartile ranges. **d,** Cross-cell 6-mer enrichment profiles of the 62 shared RBPs. Left, enrichment z scores for all 4,096 6-mers in K562 and HepG2 eCLIP peaks relative to matched non-peak regions. The heatmaps use identical RBP and 6-mer ordering and a shared color scale. Right, GC content of K562-enriched 6-mers versus the cross-cell 6-mer enrichment shift, quantified for each RBP as the mean absolute K562–HepG2 difference in enrichment z score across the full 6-mer profile. **e,** Cross-cell 6-mer enrichment shifts versus changes in transfer performance across the same 62 RBPs. Mean ΔAUC across the five models was calculated as HepG2 minus K562, and the horizontal dashed line marks no change in AUC. For both scatter plots, each point represents one shared RBP; solid lines show linear fits and shaded bands indicate 95% confidence intervals (CI). Spearman ρ and associated P values are shown in the panels. **f,** Cross-cell comparison of RiboGRAM-RPIplus-derived 6-mer motifs for 75 RBPs shared between K562 and HepG2. RBPs are ordered by hierarchical clustering based on K562 motif similarity; inner and outer rings show the K562 and HepG2 motifs, respectively. Shaded sectors indicate groups of related K562 motifs. **g,** Peak-centered positional organization of six RBP motif classes. Left, peak-minus-non-peak motif occupancy profiles in 101-nt eCLIP peak-centered windows, aggregated by motif class across 138 K562 and 105 HepG2 RBPs. Solid and dashed curves denote K562 and HepG2, respectively. Dark and light points mark, for individual RBPs, the peak-occupancy positions of the top-ranked motif and the five highest ranked motifs, respectively. Right, two representative RBPs from each motif class in each cell line are shown with their positional profiles and corresponding PWMs. A-rich and AU-rich motifs are grouped as AU/A-rich. Vertical dotted lines mark the eCLIP peak center.

Across 112 RBPs in K562 cells, RiboGRAM-RPI achieved a mean area under the receiver operating characteristic curve (AUC) of 0.912 and a mean accuracy of 0.841, compared with 0.848 and 0.742, respectively, for HDRNet^34^, the strongest baseline model (**Fig. 4b** and **Supplementary Fig. 11a**). For cross-cell evaluation, the K562-trained RiboGRAM-RPI was applied to HepG2 data for the 62 RBPs shared between the two cell lines, without cell-type-specific retraining. The model achieved a mean AUC of 0.771 and a mean accuracy of 0.689, compared with 0.742 and 0.660 for HDRNet^34^ (**Fig. 4c** and **Supplementary Fig. 11b**).

To examine whether variation in cross-cell transfer performance was related to changes in RBP-associated sequence profiles, we quantified the enrichment of all 4,096 6-mers in eCLIP peaks relative to matched non-peak regions for the 62 RBPs shared between K562 and HepG2 (**Fig. 4d** and **Supplementary Fig. 11c**). RBPs whose K562 peaks were enriched for U/AU-rich 6-mers showed higher cross-cell predictive performance and smaller enrichment shifts, whereas those enriched for G/GC-rich 6-mers showed lower performance and larger shifts (**Supplementary Fig. 11c,d**). Across RBPs, higher GC content among K562-enriched 6-mers was associated with larger cross-cell enrichment shifts (Spearman *ρ* = 0.41; **Fig. 4d**) and lower similarity between the complete K562 and HepG2 6-mer enrichment profiles (Spearman *ρ* =-0.41; **Supplementary Fig. 11e**). Larger enrichment shifts, in turn, were associated with greater declines in mean AUC across the five models (*ρ* =-0.59; **Fig. 4e**), with similar associations for accuracy and across individual models (**Supplementary Fig. 11f,g**). These shifts were also associated with greater redistribution of eCLIP peaks across the 5’ UTR, CDS, 3’ UTR and ncRNA regions (*ρ* = 0.36; **Supplementary Fig. 11h,i**).

To assess the effect of eCLIP data scale, RiboGRAM-RPI was further post-trained using increasing numbers of K562 eCLIP peaks per RBP (**Supplementary Fig. 12a**). RiboGRAM-RPI^plus^, obtained with up to 100,000 K562 peaks per RBP, showed the highest overall performance, with mean AUC increasing from 0.912 to 0.927 in K562 and from 0.771 to 0.788 in HepG2 transfer, with corresponding gains in accuracy (**Supplementary Fig. 12b,c**). Using the expanded K562 and HepG2 eCLIP collections, we then derived representative RBP-associated motifs from nucleotide-level attention signals. Among RBPs with available references, the derived motifs matched reference motifs in mCrossBase^35^ or CisBP-RNA^36^ for 70 of 89 K562 RBPs and 58 of 74 HepG2 RBPs (TOMTOM^37^ *P* ≤ 0.05; **Supplementary Fig. 12d,e**). Across the 75 RBPs shared between K562 and HepG2, motifs grouped into multiple classes, with variation both among RBPs and between cell types (**Fig. 4f**).

We next asked whether these motif classes showed distinct positional distributions relative to eCLIP peaks. Mapping representative motifs across 101-nt peak-centered sequences and matched non-peak sequences revealed RBP-specific positional motif-occupancy profiles in both cell types (**Supplementary Fig. 12f,g**). The motif classes showed distinct peak-relative patterns. U-rich motifs were enriched upstream of the peak center, AU/A-rich motifs toward both flanks, and GU-rich motifs showed broader enrichment extending from upstream regions to the peak center, whereas G-rich, GC-rich and C-rich motifs were concentrated near the peak center (**Fig. 4g**). Independent nucleotide-and k-mer-level analyses showed similar peak-relative positional patterns (**Supplementary Fig. 13a-d**). Peak-relative RNA accessibility profiles measured by icSHAPE^38^ showed less RBP-specific variation, with increased reactivity at the shoulders flanking the peak center in both cell types (**Supplementary Fig. 14a,b**). Thus, RBP-associated sequence organization varied in both motif identity and peak-relative positioning.

To assess whether this framework was applicable to other molecular interaction settings, we evaluated it on human RNA–RNA and assay-defined RNA–chromatin interaction datasets. RNA–RNA interaction pairs showed greater local complementarity potential and GC-content similarity than matched controls (**Supplementary Note 4**). For RNA–chromatin interactions, post-training on assay-defined contacts yielded representations associated with complementarity potential, RNA class and transcript identity (**Supplementary Note 5**). In both settings, RiboGRAM showed higher predictive performance than the corresponding comparison models. Together, these results indicate that RiboGRAM captures RNA sequence features associated with different interaction types and cellular contexts, including the positional organization of RBP-associated motifs.

## Discussion

In the diverse ncRNA repertoire, sequence information is reflected not only in individual features but also in their organization across scales and along the 5’-to-3’ direction. Across species, ncRNAs differed from CDS and mRNA in both motif composition and directional organization at multiple sequence scales. These patterns informed the design of RiboGRAM, which integrates multi-scale local encoding with contextual sequence modeling to support analyses of RNA sequence organization, structure and molecular interactions.

The learned representations resolved organization in RNA sequence space, from broad RNA types through ncRNA classes and families to sequence-and position-specific contextual patterns. Several of these signals corresponded to established ncRNA elements, including the conserved C-box (UGAUGA) and D-box (CUGA) regions in C/D-box snoRNAs^24^. Beyond these conserved elements, contextual dependency scores were elevated within low-complexity A-or U-rich tracts of selected lncRNAs, consistent with the U-rich or AU-rich patterns identified independently of the model. Such organization extends earlier observations relating lncRNA k-mer composition to protein association and subcellular localization even when primary sequence similarity is limited^6^. In our analysis, integrating multiple k-mer scales improved ncRNA discrimination across held-out species relative to triplets alone, and ablations supported the combination of the multi-scale local encoding and contextual sequence modeling.

The learned RNA space is shaped not only by how sequences are modeled, but also by the biological composition of the pretraining corpus. This was evident in architecture-matched controls, in which CDS-and mRNA-pretrained models showed little recoverable base-pairing information compared with ncRNA-rich pretraining. Large-scale ncRNA pretraining in RNA-FM^13^ and RiNALMo^10^ has also yielded representations useful for RNA structure prediction. Biological priors can likewise shape RNA representations by making motif information, base-pairing compatibility or biological relatedness explicit during training. RNAErnie^12^ uses motif-and subsequence-aware masking, ERNIE-RNA^39^ incorporates base-pairing informed attention bias, and Orthrus^18^ treats splice isoforms and orthologous transcripts as related sequence pairs for contrastive learning. RiboGRAM self-attention supported recovery of canonical and wobble pairs as well as pseudoknot-associated crossing pairs. Across ArchiveII classes, RiboGRAM achieved class-averaged pairing recovery similar to that of the structure-supervised RNet-SS^27^. Frozen RiboGRAM representations further supported 3D modeling of short RNAs without target-specific MSA input, whereas both pairing recovery and 3D accuracy declined for longer RNAs. Pretrained representations may therefore complement evolutionary, thermodynamic or experimental constraints for larger and more complex RNAs.

Endogenous RBP binding introduces an additional layer of context, with sequence preference expressed within a cellular binding landscape^4, 40^. Across RBPs shared between the human K562 and HepG2 cell lines, larger shifts in RBP-associated sequence preferences were accompanied by greater redistribution of eCLIP peaks across transcript regions and larger declines in prediction performance when the K562-trained model was evaluated on HepG2 data. These cross-cell effects are consistent with previous work showing that RBP-binding prediction is sensitive to cellular context, with PrismNet^41^ incorporating *in vivo* RNA structure and HDRNet^34^ explicitly addressing prediction across cell types. RiboGRAM further resolved a positional organization of RBP-associated sequence preference, with motif classes showing distinct distributions relative to eCLIP peak centers. Binding-site position, motif spacing and local sequence context are known to modulate regulatory outcomes in specific RBP systems, including Nova^42^ and Pumilio^43^ proteins. Peak-relative icSHAPE profiles were comparatively similar across RBPs, in contrast to the more RBP-specific sequence preferences. Distinct sequence association in RNA–RNA and assay-defined RNA–chromatin interactions reinforced this view, indicating that informative RNA sequence features vary with both the molecular partner and biological context.

A central challenge is to determine how organization in ncRNA sequence space contributes to function through molecular interactions. This relationship is illustrated by the lncRNA NEAT12, where UG-repeat tracts at distinct positions mediate different modes of TDP-43 recruitment, affecting paraspeckle stability and stress-associated cytoprotection^44^. In our 5S rRNA analysis, RiboGRAM mutation scores showed modest correspondence with experimentally measured effects on 60S incorporation but did not reproduce the enrichment of functional impairment at positions contacting the ribosomal proteins uL5 (RPL11) or uL18 (RPL5) (**Supplementary Note 1**). This discrepancy highlights that functional constraints associated with protein contacts are not fully reflected in sequence-derived scores. Understanding this relationship will require elucidating how specific molecular partners recognize positional and multi-scale sequence features, and how cellular context shapes the functional consequences of such recognition.

RiboGRAM converts multi-scale and positional ncRNA grammar into a reusable sequence representation. The web server (www.ultrarnalab.com/ribogram_usage.php) supports ncRNA family comparison, structural analysis and regulatory interaction modeling through pretrained embeddings and task-specific models. This framework provides an interpretable basis for investigating ncRNA primary sequence organization across structural and regulatory contexts.

## Materials and Methods

### Scale-resolved k-mer profiling across organisms and RNA types

Annotated CDS, mRNA and ncRNA FASTA files were obtained from the Ensembl^45^ FTP resources for nine model organism reference sets (Supplementary Table 1), comprising *Escherichia coli* K-12 MG1655 (Ensembl Bacteria release 57), *Saccharomyces cerevisiae (*Ensembl release 109), *Arabidopsis thaliana (*Ensembl Plants release 57), *Drosophila melanogaster (*Ensembl release 109), *Caenorhabditis elegans (*Ensembl release 109), *Danio rerio (*Ensembl release 109), *Rattus norvegicus* (Ensembl release 109), *Mus musculus (*Ensembl release 109) and *Homo sapiens* (Ensembl release 109). For the downstream analyses, mRNA denotes annotated cDNA transcript sequences. All annotated sequences were converted to uppercase and characters other than canonical bases A/C/G/U were excluded before k-mer counting.

At each k-mer scale, defined by the k-mer length *k*, species-and RNA-type-specific profiles were generated by counting sliding k-mers with the step size of 1 nt. Counts were represented over the full 4^k^ vocabulary, with absent k-mers assigned zero counts. To make profiles comparable across species and RNA types with different sequence totals, raw counts were then normalized to counts per million total k-mer observations (CPM) at the same k-mer scale and transformed by log2 (CPM+1). For a species *s*, RNA type *t*, k-mer scale *k* and k-mer *m*, the normalized profile was computed as:

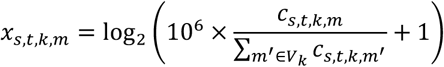

where *c*_s,t,k,m_ is the raw count of k-mer *m*, and *V*_k_ denotes the full 4^k^ k-mer vocabulary.

For each RNA type and k-mer scale from k = 1 to 9, cross-species similarity was calculated as the Pearson correlation between normalized species-level k-mer profiles. At a given k-mer scale *k*, each species-level profile consisted of 4^k^ normalized k-mer features. Profile distance between two species was defined as *D = 1 − r*, where *r* is the Pearson correlation between their normalized k-mer vectors. Distances were calculated for all 36 species pairs among the nine model organisms. For the scale-resolved ncRNA analysis, the profile-distance matrices were computed at k = 1, 3, 5, 7 and 9. Hierarchical clustering was then applied to the species-pair distance matrix at each k-mer scale to visualize ncRNA profile relationships among organisms.

To compare scale-dependent behavior among RNA types, the same species-pair distance calculation was repeated for CDS, mRNA and ncRNA from k = 1 to 9. Relative k-mer grammar distance was defined as a paired species-pair difference between two RNA types for the same species pair and k-mer scale. For RNA type *A* and *B*, the relative distance for a species pair *(i, j)* was calculated as:

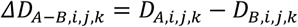

where *D*_A,i,j,k_ and *D*_B,i,j,k_ are the cross-species profile distances for RNA types *A* and *B*, respectively. The comparisons included ncRNA − CDS, ncRNA − mRNA and mRNA − CDS. Positive values indicate that the first RNA type has a larger cross-species profile distance than the second RNA type at the same k-mer scale.

### Frame-aware triplet and k-aware motif-class enrichment

To identify interpretable local sequence features that distinguish ncRNA from other RNA types, motif-class enrichment was quantified from species-level k-mer frequency profiles. For each species and motif class, class frequency was calculated by summing the frequencies of all k-mers assigned to that class. Enrichment of motif class *C* was defined as the log2 ratio of ncRNA motif-class frequency to the corresponding same-species reference frequency:

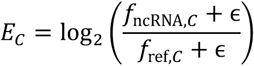

where *f*_ncRNA,C_ and *f*_ref,C_ are the motif-class frequencies in ncRNA and the reference RNA type, respectively. *ε* = 10^-10^ was added to the numerator and denominator to avoid undefined ratios.

Stop-codon-like triplets were defined as UAA, UAG, and UGA during the triplet analysis. Since these triplets are frame-constrained in CDS but frame-independent in ncRNA, the primary stop-codon-like comparison was conducted using sliding ncRNA 3-mer profiles against in-frame non-overlapping CDS codon profiles. In parallel, sliding ncRNA 3-mer profiles with sliding CDS 3-mer profiles and phase-averaged non-overlapping ncRNA triplet profiles with in-frame CDS codon profiles were also compared and averaged across nine model organisms. For the phase-averaged ncRNA control, non-overlapping triplets were counted from all three possible offsets and averaged together, with incomplete terminal fragments excluded. The same enrichment framework was computed for nucleotide-, dinucleotide-and all 64 codon triplets.

U-rich and AU-rich local patterns were assigned using the same k-aware, mutually exclusive motif definition in the enrichment analyses. Stop-codon-like triplet were assigned first, with k = 3. For k ≥ 4, U-rich k-mers were assigned before AU-rich k-mers. U-rich k-mers were defined as having at least three U bases at k = 4, and U at ≥ 60% of positions for k ≥ 5. AU-rich k-mers require at least three A/U bases with both A and U present at k = 4, and at least 0.75k A/U bases with both A/k and U/k ≥ 0.25 for k ≥ 5. This priority scheme was designed to prevent strongly U-rich k-mers from being double-counted in the AU-rich class.

To extend the selected motif-class analysis to a broader set of local sequence patterns, systematic enrichment was quantified for k = 4 to 9 motifs in ncRNA compared with same-species CDS and mRNA references using species-level sliding k-mer frequency profiles with a more stringent composition-rich scan. For each k-mer, composition-rich motif classes were assigned using a fixed 0.8 fraction threshold. Single-base-rich classes were assigned when the corresponding nucleotide occurred at least 80% of the k-mer. Two-base-rich classes, including AU-rich, AG-rich, AC-rich, UG-rich, UC-rich and GC-rich k-mers, were assigned when the two corresponding nucleotides together accounted for in at least 80% of the k-mer positions. Single-base-rich classes were assigned first. Two-base-rich classes were assigned only to k-mers that did not already satisfy a single-base-rich definition. Repeat and complexity-related classes were defined independently from the composition-rich classes and were allowed to overlap with them. Homopolymers were defined as k-mers composed of a single repeated nucleotide. Dinucleotide-repeat-like k-mers were defined as exact two-nucleotide periodic repeats, generated by repeatedly copying the first two nucleotides to length k; for odd k, the terminal nucleotide was required to follow the same periodic pattern. Low-complexity k-mers were defined as k-mers containing no more than two distinct nucleotides. For both CDS and mRNA references, species-level class enrichments were computed using the log2 ratio framework described above and summarized for each pattern class and k-mer scale by the cross-species mean and the fraction of species with positive enrichment.

### Position-resolved k-mer and motif-class gradient analysis

To quantify positional sequence organization, sequences of at least 60 nt were partitioned into ten approximately equal length-normalized 5’-to-3’ bins using integer boundaries from Python Numpy.linspace (0, L, 11), where L is the sequence length. For lengths not divisible by ten, nucleotides were assigned according to these boundaries, so no terminal nucleotides were discarded. For each species, RNA type, and k-mer scale, sliding k-mers were counted separately within each bin. Original sequences were analyzed together with five sequence-shuffled controls generated by randomly permuting the bases within each sequence while preserving single-nucleotide composition.

For position-resolved profile drift, bin-wise k-mer count vectors were converted to frequency profiles. At each k-mer scale, the profile of each positional bin was compared either with the 5’-proximal bin from the same species and RNA type or with the same species global CDS profile at the same k-mer scale. Cosine similarity to the 5’ bin quantified within-transcript profile drift, while cosine similarity to the same-species CDS profile quantified divergence from CDS sequence organization. Species-level similarity profiles were averaged across the nine model organisms. Positional trends were summarized using Spearman correlation, linear slope and end-to-end change across the ten bins. For the 6-mer drift analysis, distal similarity was averaged over bins 8-10, 5’-to-distal drift was defined as the difference between bin-1 similarity and distal similarity, and distal divergence was calculated as the original-shuffled difference:

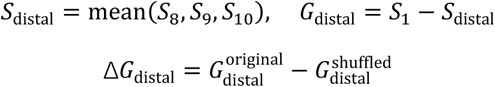

where *S*_b_ denotes the cosine similarity of bin *b* to the corresponding reference profile.

Using the selected motif classes defined above, positional gradients were evaluated separately for stop-codon-like 3-mers and k-aware U-rich and AU-rich 4-to 7-mers. For matched-background construction, each motif class was compared with the same-length k-mers outside the same motif class and with similar nucleotide composition. For U-rich and AU-rich motif classes, matched backgrounds were required to have U counts and total A/U counts within one nucleotide of the motif k-mer. Stop-like triplets were matched to non-stop-like triplets with total A/U counts within one nucleotide. If fewer than ten matched background k-mers were available, all non-class k-mers of the same length were used as a fallback. This correction separated motif-class-specific positional behavior from broader trends associated with similar base composition.

Motif-class excess trajectories were computed for each positional bin as the mean bin-wise log2CPM profile of motif class k-mers minus the corresponding matched-background profiles. Species-level trajectories were summarized as the cross-species mean. Positional motif excess was quantified by Spearman excess, defined for each motif-class k-mer *m* as:

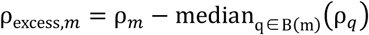

where *ρ*_m_ is the Spearman correlation between bin index and the bin-wise log2CPM profile of k-mer *m*, and *B(m)* denotes its matched non-class background.

Positive values indicate stronger monotonic 5’-to-3’ increase than expected from composition-matched non-class k-mers. Late-minus-early excess and linear slope excess were used as sensitivity metrics, comparing distal versus proximal bins and linear positional trends, respectively.

### Species-held-out RNA-type classification

To evaluate the cross-species performance of k-mer-based RNA-type classification, a leave-one-species-out framework was utilized for the analysis. CDS, mRNA and ncRNA sequences from the nine-organism reference were used, with sequences shorter than 20 nt excluded.

In the binary task, ncRNA sequences were classified against a pooled coding-related class consisting of CDS and mRNA sequences. For each species and sampling repeat, up to 5,000 ncRNA sequences were sampled, and an equal number of coding-related sequences was sampled with approximately balanced CDS and mRNA contributions. In the three-class task, CDS, mRNA and ncRNA were classified separately, with up to 3,000 sequences sampled per RNA type per species and the sample size limited by the least abundant RNA type. Species with fewer than 50 eligible sequences for the required class were excluded from the corresponding repeat. The sampling procedure was repeated 20 times using predefined random seeds, with each repeat using an independent fixed seed. Sparse sliding k-mer frequency vectors were generated for each sequence at k = 1, 3, 5, 7 and 9. Individual k-mer scales were analyzed separately, and a multi-scale feature set was constructed by concatenating the 1-, 3-, 5-, 7-and 9-mer vectors. Counts were normalized within each k-mer scale by the total number of valid k-mer observations in the sequence before concatenation.

For each held-out species, an L2-regularized stochastic-gradient logistic classifier was trained on sequences from the remaining eight species and evaluated by balanced accuracy, using α = 10^-5^, max_iter = 1,000 and tol = 10^-4^. Multi-scale gain was calculated as the balanced-accuracy difference between the concatenated 1/3/5/7/9-mer and 3-mer-only feature sets for each held-out species. Paired comparisons across held-out species used one-sided paired Wilcoxon signed-rank tests.

### RiboGRAM architecture and pre-training

RiboGRAM is a self-supervised RNA language model pretrained on ncRNA sequences. The model comprises a 12-layer hybrid encoder with a hidden dimension of 1,280 and 20 attention heads. Each encoder layer contains a multi-head self-attention branch for modeling sequence-wide contextual dependencies and five parallel one-dimensional convolutional branches with kernel sizes of 1, 3, 5, 7 and 9 nt for multi-scale local sequence encoding. A learned position-wise gate assigns weights to the five convolutional outputs, which are aggregated and fused with the self-attention representation before processing by a SwiGLU^46^ feed-forward block. Rotary position embeddings (RoPE)^47^ are utilized to encode sequence position.

The pre-training corpus was assembled from RNAcentral 25.0^19^, Rfam 15.0^20^ and Ensembl release 115^21^. After deduplication at 100% sequence identity under a coverage rate of 0.8 using MMseqs2^48^, the corpus comprised approximately 40 million non-redundant ncRNA sequences. Sequences were tokenized at the single-nucleotide level, with <CLS> and <EOS> marking sequence boundaries, <UNK> representing unknown nucleotides and <PAD> used for padding. Sequences longer than 1,024 nt were randomly cropped to contiguous 1,024-nt segments during pre-training. For masked language modeling, 15% of nucleotide positions were selected as prediction targets. Among these masked nucleotide positions, 80% were replaced with <MASK>, 10% with a randomly selected nucleotide and 10% were left unchanged. Cross-entropy loss was calculated only at the selected positions. The model was optimized using AdamW^49^ with an initial learning rate of 1 × 10^-4^, *β*_1_ = 0.9, *β*_2_ = 0.98 and a weight decay of 0.01. The learning rate was reduced to 1 × 10^-5^ using a cosine schedule, and gradients were clipped to a maximum norm of 2.0. Pre-training was performed for approximately four weeks using distributed mixed-precision training on eight NVIDIA A800 GPUs with 80 GB of memory each.

### Architectural variants for RiboGRAM ablation analysis

To assess the contribution of contextual self-attention and multi-scale convolutional encoding, we pretrained 11 model variants: three CNN-only models, one Transformer-only model and seven hybrid Transformer–CNN models. The CNN-only variants omitted self-attention and used either a single 1-nt kernel, paired 1-and 3-nt kernels, or seven parallel kernels of 1, 3, 5, 7, 9, and 11-nt. Each convolution preserved sequence length and used 1,280 input and output channels. For variants with multiple kernels, a 1-nt gating convolution generated position-wise softmax weights across the convolutional branches. The weighted branch outputs were summed, combined with the layer input through a residual connection and linearly projected before entering a pre-normalized SwiGLU feed-forward block. The Transformer-only variant omitted the convolutional branches, convolutional gate and attention–convolution fusion projection, with the residual self-attention representation passed directly to the SwiGLU block.

The seven hybrid variants retained both self-attention and convolutional encoding and comprised two single-kernel modules using either a 1-or 3-nt kernel, one two-kernel model using 1-and 3-nt kernels, and four multi-kernel models using 1/3/5-, 1/3/5/7-, 1/3/5/7/9-or 1/3/5/7/9/11-nt kernels. Within each encoder layer, the self-attention and convolutional branches operated in parallel on separately normalized representations of the same layer input. Each branch output was independently combined with the input through a residual connection. The resulting representations were then concatenated, fused by a linear projection and passed through the SwiGLU block. The hybrid configuration with 1/3/5/7/9-nt kernels defined the full RiboGRAM architecture used in subsequent analyses. All variants comprised 12 encoder layers, a hidden dimension of 1,280, a SwiGLU intermediate dimension of 5,120 and the same masked language model prediction head. Variants containing self-attention used 20 attention heads and RoPE. All models were pretrained on the same corpus using the same sequence processing, masking scheme, reconstruction objective and learning rate schedule.

### Frozen-encoder cross-species evaluation of model architectural variants

To evaluate the pretrained architecture variants under a common RNA-type classification framework, two binary classification tasks were defined to distinguish ncRNA from CDS and from mRNA. For each task, 5,000 human sequences from each class were used for training, yielding a balanced training set of 10,000 sequences. The remaining human sequences and sequences from the other eight model organisms constituted the candidate evaluation dataset. To reduce sequence similarity between training and evaluation datasets, candidate sequences sharing at least 80% sequence identity with any training sequence from the same RNA class were removed using MMseqs2^48^. Following sequence-similarity filtering, 1,000 human sequences per class were assigned to the validation set. The remaining eligible sequences were organized into species-specific test sets across all nine species, with up to 5,000 sequences retained per class for each species (Supplementary Table 2).

Each pretrained CNN-only, Transformer-only and hybrid Transformer-CNN encoder was evaluated using the same frozen-encoder classification procedure. Hidden representations at non-padding nucleotide positions were mean-pooled to obtain a sequence-level representation and passed to an identical two-layer multilayer perceptron (MLP) classifier comprising a 2,048-unit hidden layer with ReLU activation and a two-unit output layer. Encoder parameters were kept fixed throughout classifier training, and only the MLP parameters were optimized. Model selection was performed on the human validation set, and classification performance was calculated separately for each species using macro-F1 and accuracy.

### Linear probing for ncRNA category and family classification

For category-level classification, we used the 13-class NCYPred^23^ benchmark (Supplementary Table 3), comprising 45,447 ncRNA sequences derived from Rfam 14.3^50^ and including 5S rRNA, 5.8S rRNA, C/D-box snoRNA, H/ACA-box snoRNA, tRNA, miRNA, riboswitch, ribozyme, leader RNA, group I intron, group II intron, Y RNA and Y RNA-like RNA. During construction of the benchmark, sequences sharing more than 80% sequence identity were removed using CD-HIT^51^. Following the partitioning scheme used in RNA-Scope^52^, training (28,626 sequences), validation (3,175 sequences), and test sets (13,646 sequences) were assigned.

For family-level classification, the Rfam-novel benchmark^53^ (Supplementary Table 4), which was derived from Rfam 14.1^50^ and contained 306,016 sequences from 88 ncRNA families before construction of the final balanced partitions, was utilized. Within each family, sequences were initially assigned to training, validation and test sets. Only validation and test sequences whose normalized Hamming similarity to every training sequence was below 0.50 were retained. After the class-balancing procedure defined for the benchmark, the resulting partitions contained 105,864 training, 17,324 validation and 25,342 test sequences.

For all linear-probing analyses, the parameters of the pretrained RNA model encoders were kept fixed. For comparison across models, token-level representations were extracted from the final encoder layer and mean-pooled over non-padding positions to obtain a fixed-length sequence representation. This representation was passed directly to a single trainable linear classification layer. Only the parameters of the linear probes were optimized, using cross-entropy loss and AdamW with a learning rate of 1 × 10^-3^ and a weight decay of 0.01. The same probe architecture and optimization settings were used for all models. For layer-wise analysis of RiboGRAM, separate linear probes were trained on mean-pooled representations extracted independently from each of the 12 encoder layers. Classification performance was evaluated using accuracy and macro-averaged precision, recall and F1.

### Layer-wise UMAP visualization of ncRNA class representations

To visualize the organization of ncRNA representations across encoder layers, token-level embeddings were extracted from each of the 12 layers of the frozen RiboGRAM encoder for all 45,447 sequences in the 13-class NCYPred dataset^23^. At each layer, embeddings from valid nucleotide positions were mean-pooled, excluding padding and special tokens, to obtain a 1,280-dimensional sequence-level representation. Representations from each layer were projected independently into two dimensions using uniform manifold approximation and projection (UMAP)^54^, with 100 nearest neighbors, a minimum distance of 1.0 and Euclidean distance as the distance metric.

### Layer-resolved profiling of APC-adjusted self-attention

Self-attention weights were extracted from each of the 12 RiboGRAM encoder layers. Within layer P, attention weights were averaged across heads to obtain a head-averaged matrix A(P) ∈ ℝ^L × L^, where A(P);ⱼ denotes attention from query position i to key position j, and L is the sequence length. Average product correction (APC) was applied to reduce row-and column-dependent background effects:

^∼^(P)

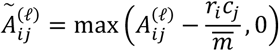

where *r;* and *cⱼ* are the mean attention values in row *i* and column *j*, respectively, and m̄ is the global mean of A(P).

Layer-specific attention properties were quantified using a class-balanced set of 1,300 NCYPred sequences, comprising 100 randomly sampled sequences from each of the 13 ncRNA classes. All metrics were calculated independently for each sequence before aggregation across sequences within each layer. To obtain a normalized measure of attention received by each nucleotide position, the APC-adjusted matrix was first normalized across key positions within each query row:

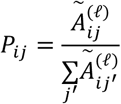

Rows with zero total APC-adjusted attention were retained as zero-valued rows. The total attention received by key position *j* was then calculated as:

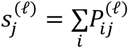

Attention concentration across nucleotide positions was quantified from the received-attention distribution using the Gini coefficient:

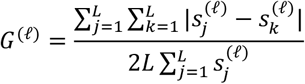

Lower Gini coefficients indicate a more even distribution of received attention across positions, whereas higher coefficients indicate greater concentration on a subset of positions. One Gini coefficient was calculated for each sequence and layer.

To quantify nucleotide-specific attention enrichment, received attention was converted into a positional attention fraction:

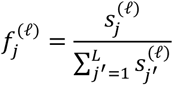

For each nucleotide *b* ∈ {*A*, *C*, *G*, *U*}, enrichment was defined as

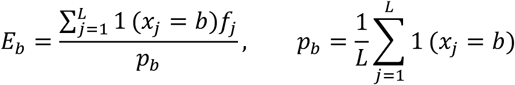

where *p*_b_ is the sequence-level abundance of nucleotide *b*. An enrichment value of 1 therefore indicates that the fraction of received attention assigned to a nucleotide type is proportional to its sequence abundance, whereas values above or below 1 indicate relative enrichment or depletion, respectively.

For position-resolved base attention, the positional attention fraction was normalized relative to the uniform allocation across nucleotide positions:

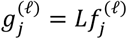

such that *g*^(P)^ = 1 corresponds to uniform received attention. Each sequence was mapped to a length normalized 5’-to-3’ coordinate and partitioned into 20 positional bins. For each base *b* within positional bin *β*, normalized attention was summarized as

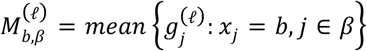

Sequence-level values were subsequently averaged across the balanced sequence set to obtain layer-, nucleotide-and position-specific attention profiles.

Representative analyses of conserved ncRNA features were performed using C/D-box snoRNAs, tRNAs and H/ACA-box snoRNAs. For layer-7, six representative sequences from each defined ncRNA category were aligned, and position-level APC-adjusted attention values were mapped to the corresponding alignment coordinates. Conserved C-and D-box regions in C/D-box snoRNA, the tRNA T-loop and the H-and ACA-box regions of H/ACA-box snoRNAs were annotated on the aligned profiles. For layer-12, two-dimensional APC-adjusted attention maps were generated for representative RNAs with nucleotide-level paired and unpaired annotations, and off-diagonal attention patterns were compared quantitatively with the corresponding Rfam-annotated secondary structures.

### Entropy-weighted masked-recovery scoring of contextual dependency

At the nucleotide level, contextual dependency was formulated as confident recovery of a masked wild-type nucleotide from its surrounding sequence context, combining a transformed recovery score with inverse predictive entropy:

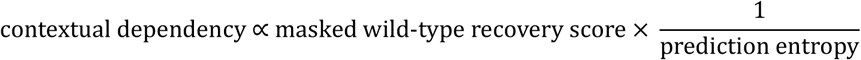

For each nucleotide position j in sequence n, the wild-type nucleotide *x*n,j was replaced with a mask token <MASK> while the remaining sequence was retained as context. The pretrained RiboGRAM model was then used to generate a probability distribution *p*n,j(*a*) over the nucleotide vocabulary *V*. The masked wild-type recovery score was defined as:

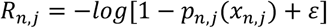

and the entropy of the predicted nucleotide distribution was calculated as:

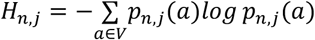

The nucleotide-level contextual dependency score was defined as:

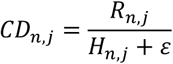

where ε = 10⁻⁸ was added for numerical stability. The negative-log transformation increased the separation among high-probability wild-type recoveries. Higher *CD*n,j values therefore indicate positions that were recovered with both high probability and low predictive uncertainty from the surrounding sequence context.

Seed sequences from all 4,178 RNA families in Rfam 15.0 were analyzed and organized according to the Rfam annotation hierarchy, including the three broad classes — gene, intron, and cis-regulatory — and their corresponding ncRNA subtypes (Supplementary Table 5). For a sequence of length *L*n, sequence-level contextual dependency was calculated as the mean position-specific scores:

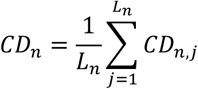

Within each family *r*, sequences were ranked by *CDn*, and up to 200 of the highest-scoring sequences were retained. The family-level contextual dependency score was calculated as

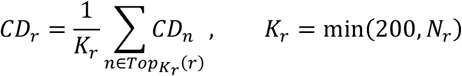

where *Nr* is the number of available seed sequences in family *r*. For ncRNA subtype *t*, contextual dependency was calculated as the arithmetic mean across its constituent family-level scores:

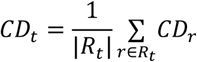

where *Rt* denotes the set of Rfam families assigned to subtype *t*. This two-stage aggregation assigned equal weight to each family, irrespective of the number of available seed sequences.

### Family-stratified contextual dependency motif enrichment

To identify local sequence patterns associated with high contextual dependency, we performed a family-stratified 6-mer enrichment analysis. Within each Rfam family, seed sequences were ranked by their mean nucleotide-level contextual-dependency scores, and up to the 200 highest-scoring sequences were retained. Position-specific scores were subsequently smoothed and normalized within each sequence to reduce differences in overall score scales. Each 6-mer occurrence was weighted by the mean normalized contextual dependency score across its six positions. For each family, enrichment of each 6-mer was calculated as the log2 ratio between its fraction of the total contextual dependency signal and its fraction of all 6-mer occurrences.

To prevent families with larger sample sizes from dominating the analysis, family-level enrichment values were aggregated with equal weights across families. Candidate 6-mers were ranked by mean enrichment, the number of supporting families, and family-level coverage. For each ncRNA subtype, up to the two highest-ranking non-redundant 6-mers were selected as candidate seed motifs. Sequence windows with a Hamming distance of one from each seed motif were then collected and weighted by their local contextual-dependency scores. Nucleotide frequency matrices were calculated separately within each family and averaged with equal family weights to generate balanced position weight matrices for sequence logo visualization.

### Zero-shot attention-based ncRNA base-pair prediction

A training-free attention probe was utilized to assess whether individual attention heads encoded RNA base-pairing information without updating model parameters. The analysis included RiboGRAM, RiNALMo10, RNA-FM13, RNAErnie212, 3UTRBERT15, SpliceBERT9, UTR-LM16, together with two architecture-matched PLM-CDS and PLM-mRNA controls pretrained on CDS and mRNA sequences, respectively. For sequence *n*, attention matrices *A*^(P,ℎ)^ were extracted separately from each layer P and attention head ℎ. After removal of special token positions, each matrix was symmetrized and subjected to APC adjustment:

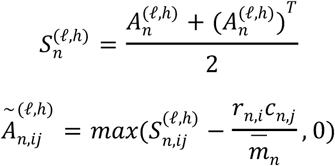

where *rn*,*i*, *cn*,*j* and *m*n are the row, column and global means of the symmetrized matrix, respectively. An annotated binary contact matrix *Y*n was constructed for each sequence, with *Yn*,*ij* = 1 when positions *i* and *j* formed an annotated base pair and *Yn*,*ij* = 0 otherwise.

Model-specific structure heads were selected using the 1,300-sequence VL0 validation partition of bpRNA-1m25, 26 (Supplementary Table 7), derived from Rfam 12.255. For each layer–head combination, Spearman correlation between the upper-triangular elements of the APC-adjusted attention and annotated contact matrix was calculated separately for each sequence and then averaged across VL0:

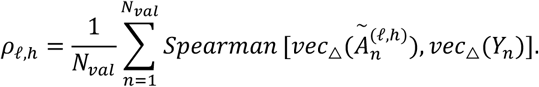

The layer–head combination with the highest mean correlation was designated the model-specific structure head:

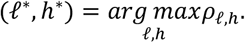

Using this fixed head, attention thresholds between 0 and 1 were evaluated on VL0. At each threshold, pair-level F1 was calculated separately for each validation sequence, and the threshold maximizing the macro-averaged F1 across sequences was retained:

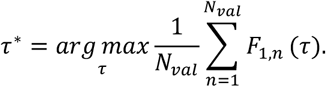

Base-pair predictions were then defined as

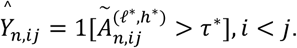

The structure head and threshold were determined independently for each model and fixed before downstream evaluation. Zero-shot performance was evaluated on the 10,814-sequence TR0 partition of bpRNA-1m, which was not used for model fitting, structure-head selection or threshold calibration. Pair-level micro-averaged precision, recall and F1 were calculated by pooling predicted and reference pairs across sequences. Macro-averaged metrics were obtained by averaging the corresponding sequence-level metrics.

Pair-type-specific performance on TR0 was evaluated separately for canonical and wobble pairs, comprising A–U, G–C and G–U, and for all remaining non-canonical nucleotide combinations, with micro-precision, micro-recall and micro-F1 calculated separately for each category. For pseudoknots, evaluation was restricted to base pairs involved in at least one crossing interaction, defined by *i* < *k* < *j* < *l*. Reference crossing pairs were present in 999 of the 10,814 TR0 sequences, and true-positive, false-positive and false-negative counts were pooled across these sequences to calculate the micro-averaged metrics.

RNA-type-level generalization was additionally evaluated using ArchiveII29,30 (Supplementary Table 10), comprising 3,865 RNA entries from nine ncRNA types: 5S rRNA (*n* = 1,283), signal recognition particle (SRP) RNA (*n* = 918), tRNA (*n* = 557), tmRNA (*n* = 462), RNase P RNA (*n* = 454), group I intron (*n* = 74), 16S rRNA (*n* = 67), telomerase RNA (*n* = 35) and 23S rRNA (*n* = 15). The VL0-selected structure head and threshold were applied without further adjustment. Within each ncRNA type, micro-averaged metrics were calculated by pooling pair-level counts across sequences, whereas macro-averaged metrics were calculated from sequence-level precision, recall and F1 (Supplementary Tables 11 and 12).

RNAfold56, RNAstructure57, LinearFold58 and EternaFold59 were included as thermodynamic or statistical folding baselines. We additionally evaluated two pseudoknot-capable structure predictors: EternaFold with ThreshKnot decoding (EternaFold+TK)59, 60 and the structure-supervised RibonanzaNet-SS (RNet-SS)27, 61. For EternaFold+TK, EternaFold-derived base-pair probabilities were decoded using ThreshKnot60, which retains reciprocally maximal pairs above a probability threshold while permitting crossing pairs. RNet-SS is a secondary structure-supervised adaptation of RibonanzaNet61, which was trained on large-scale RNA chemical-mapping data, and outputs a pairwise base-pairing confidence matrix. This matrix was decoded using the Hungarian algorithm62 to obtain a one-partner-per nucleotide pairing assignment while permitting crossing pairs. All predicted structures were converted to base-pair sets and evaluated on the same sequences using the same pair-level micro-and macro-averaged metrics.

### 5S rRNA representation and photosynthetic plastid 5S-like candidate analyses

Records from the Rfam 5S rRNA family (RF00001) were obtained from Rfam 15.020. The seed alignment initially contained 712 records. After removing records containing ambiguous nucleotides or lacking a resolved NCBI taxonomic assignment, 677 records remained: 290 canonical eukaryotic cytoplasmic, 10 rhodophyte cytoplasmic, 5 photosynthetic plastid, 333 bacterial and 39 archaeal 5S rRNAs. Groups were assigned according to Rfam annotations and NCBI taxonomic lineages. After removing special tokens, including <CLS>, <EOS> and padding tokens, RiboGRAM nucleotide embeddings were extracted and mean-pooled across nucleotide positions to obtain sequence-level representations. The 677 seed records were visualized using UMAP, and a neighbor-joining tree was constructed from pairwise cosine distances between the sequence representations. The association between representation cosine distance and patristic distance in the phylogenetic tree corresponding to the RF00001 seed alignment was assessed using Spearman correlation, with the *P*-value calculated using a two-sided Mantel permutation test. The bacterial reference centroid was obtained by averaging the normalized representations of bacterial 5S rRNAs. Position-specific contextual-dependency and pairing signals were calculated using the masked-nucleotide recovery and pairing-partner ranking procedures described above.

Within the UMAP of the complete RF00001 collection, records were defined as photosynthetic plastid 5S-like candidates if they occupied the same region as annotated photosynthetic plastid 5S rRNAs, belonged to Viridiplantae and were originally annotated as canonical eukaryotic cytoplasmic 5S rRNAs. This identified 2,734 candidate records, corresponding to 1,246 distinct sequences from 196 TaxIDs. Sequence similarity was calculated using RapidFuzz, a normalized Indel similarity ranging from 0 to 100, with higher values indicating greater similarity. Each candidate was compared with 672 non-organellar RF00001 seed sequences and 96 annotated photosynthetic plastid/chloroplast 5S rRNAs, and the maximum similarity to each reference set was retained.

A plastid reference genome was assigned to each candidate TaxID. A complete RefSeq plastid (chloroplast) genome from the same TaxID was used when available; otherwise, the closest available reference from the same genus or family was selected. Candidate-flanking sequences were examined at ±0.5-, ±2.5-and ±5-kb scales. The candidate 5S rRNA sequence was excluded from the ±0.5-kb window. For the ±2.5-and ±5-kb windows, the candidate sequence and an additional 100 nt on either side were excluded to reduce the contribution of immediately adjacent sequences to the larger-scale estimates. Valid 18-mers were restricted to sequences containing only A, C, G and T. Plastid-reference matching was defined as the proportion of valid 18-mers that matched the corresponding plastid genome exactly.

Nuclear-background controls were selected from plant nuclear chromosomes or long scaffolds without organellar annotations, preferentially using genomes from the same or the closest available taxon as the corresponding candidate. Control windows were positioned away from RF00001 hits and matched to candidate windows by GC content. Thirteen records were excluded because their contigs were too short or contained insufficient valid 18-mers, leaving 2,721 candidate–control pairs with measurements at all three scales. Associations among window scales were summarized using Spearman correlations. For three representative loci, candidate-containing genomic intervals were globally aligned to the corresponding plastid reference intervals, and reference gene annotations were projected onto the candidate regions through the alignment coordinates. Global alignment identity and exact 18-mer matching were calculated and reported separately.

### Human 5S rRNA variant-effect analysis

Human 5S rRNA variant data were obtained from the saturation-mutagenesis study by Sengl *et al*63. The original dataset contained 247 single-nucleotide variants at 114 positions. After excluding 41 variants without measurements of incorporation into the 60S ribosomal subunit, 206 variants at 110 positions were retained. Experimental effects were defined as 1 − relative 60S-subunit incorporation.

Zero-shot model scoring used the 120-nt human 5S rRNA consensus sequence employed in the assay. For each variant, the target position was replaced by <MASK>, and the wild-type and mutant nucleotide probabilities were calculated from the same masked sequence context. The mutation score was defined as

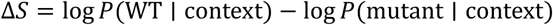

such that a larger positive value indicates a stronger model preference for the wild-type nucleotide.

RNAfold was run with the default ViennaRNA56 parameters to calculate the minimum free energy of each wild-type and mutant 5S rRNA. The folding-energy change was defined as ΔMFE = MFEmutant − MFEWT, with positive values indicating predicted destabilization. The Rfam tree-parsimony score quantified evolutionary variability at each position. For each alignment column, Fitch parsimony was used to estimate the minimum number of nucleotide changes on the reference phylogenetic tree associated with the RF00001 seed alignment, normalized by the number of non-gap sequences at that position; lower values therefore indicate greater conservation. The Rfam allele log-odds score was calculated from the wild-type and alternative nucleotide frequencies at the corresponding alignment position as log(*f*WT/*f*alt), with larger values indicating that the alternative nucleotide is less frequently observed.

RNA language models were evaluated using zero-shot masked-nucleotide scoring. Local adaptation used the six deduplicated RF00001 seed sequences most similar to the human wild-type 5S rRNA. During LoRA adaptation64, the RiboGRAM backbone was frozen and only the low-rank adapters were trained, using random single-nucleotide masking and recovery as the training objective. Variant effects after adaptation were calculated using the same zero-shot scoring procedure. Associations between predicted and experimental variant effects were assessed using Spearman correlations. For structural analysis, the 120-nt assay sequence was mapped to 5S rRNA chain L7 in the human 80S ribosome structure65 (PDB 6QZP). uL18/RPL5 and uL5/RPL11 corresponded to chains LD and LJ, respectively. A 5S rRNA nucleotide was defined as protein-contacting when its minimum heavy-atom distance to either protein was ≤ 4 Å.

### MLP probing for nucleotide-level pairing states in ncRNA

To evaluate the secondary-structure information captured by different RNA language models, the pretrained encoders of RiboGRAM, 3UTRBERT, UTR-LM, SpliceBERT, RNA-FM, RNAErnie2, RiNALMo, PLM-mRNA and PLM-CDS were kept frozen throughout training. Each nucleotide was labelled as paired if it participated in at least one annotated base pair and unpaired otherwise. For each model, the *d*-dimensional nucleotide representation, where *d* denotes the model-specific embedding dimension, was passed to an identically configured two-layer MLP with a ReLU activation. Only the MLP parameters were optimized using binary cross-entropy loss and AdamW with a weight decay of 0.01. Padding positions and special tokens were excluded from training and evaluation. Output logits were converted to probabilities using a sigmoid function and thresholded at 0.5. Performance was assessed by micro-averaged precision, recall and F1 across all valid nucleotide positions.

Two complementary evaluation settings were used. Intra-family performance was evaluated on bpRNA-1m25, derived from Rfam 12.255, using the established TR0, VL0 and TS0 partitions for training, validation and testing, respectively. Inter-family generalization was evaluated using the TrainSetA-to-TestSetB protocol28. TrainSetA66, comprising eight commonly studied RNA families, was divided into training and validation subsets, whereas TestSetB, comprising 22 structurally distinct Rfam families with less than 70% sequence identity to TrainSetA, was reserved for cross-family testing (Supplementary Table 13). The same probe architecture, optimization procedure and validation-F1 model-selection criterion were applied to all models.

### RNA structural element annotation and k-mer profiling

Structural elements were derived from the annotated base-pair assignments in the bpRNA-1m TR0 dataset and classified into six categories: stems, hairpin loops, internal loops, bulges, multiloops and exterior loops. Consecutive nested base pairs were grouped into stem segments. Contiguous unpaired regions were classified according to their enclosing helix topology, while unpaired regions outside any enclosing base pairs were assigned as exterior regions. Stem length was defined as the number of consecutive base pairs, while the lengths of unpaired structural elements were defined by the number of constituent nucleotides. Contiguous positions sharing the same structural annotation were treated as individual structural element segments for subsequent sequence analysis.

For element-specific k-mer profiling, overlapping 4-mers were enumerated within each structural element category. The frequency of each 4-mer within an element type was compared with its corresponding frequency across pooled full-length TR0 sequences, and enrichment was defined as the log2 ratio of the element-specific to background frequency. For length-specific analysis, structural-element segments of 3–7 nt were represented at their corresponding k-mer scale and compared with same-length k-mer frequencies derived from the pooled full-length TR0 sequences.

For position-resolved analysis, each contiguous structural element segment was partitioned into 5’-end, middle and 3’-end regional contexts. Within each context, 4-mer enrichment was calculated relative to the expected frequency under the corresponding nucleotide composition background. Boundary-centered profiles were generated by aligning the eight nucleotide positions upstream and downstream of each structural element boundary to a common relative coordinate. Nucleotide frequencies were calculated at each relative position. For each structural element type, position-specific nucleotide enrichment was defined by the log2 ratio of its nucleotide frequency to the corresponding pooled frequency across all structural element types at the same relative positions.

### Architecture-matched CDS-and mRNA-pretrained controls

To assess the contribution of the pre-training sequence corpus independently of model architecture and training configuration, we additionally pretrained two architecture-matched control models, PLM-CDS and PLM-mRNA. CDS and mRNA sequences were obtained from Ensembl release 11221 and deduplicated at 100% sequence identity with a coverage rate of 0.8 using MMseqs248, yielding approximately 13 million non-redundant CDS sequences and 14 million non-redundant mRNA sequences, respectively (Supplementary Table 6).

PLM-CDS and PLM-mRNA were pretrained using the same model architecture, tokenization and sequence-processing procedure, masking scheme, masked language modeling objective, optimization settings and learning-rate schedule as RiboGRAM. The models therefore differed in the RNA sequence corpus used for pretraining while retaining the same model and training configuration as RiboGRAM.

### RiboGRAM-3D architecture and training

RiboGRAM-3D was implemented as an MSA-free RNA tertiary structure prediction framework comprising a frozen RiboGRAM encoder and a trainable structure-prediction network. For each input RNA sequence, RiboGRAM was run once in inference mode with gradient computation disabled and all encoder parameters fixed. Final-layer nucleotide embeddings were linearly projected to 384-dimensional single representations, whereas multi-layer, multi-head self-attention features were projected to 128-dimensional pair representations.

The initial pair representation combined an MLP projection of the attention-derived features, a SequenceToPair transformation of the single representations and a learned relative positional embedding:

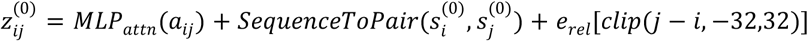

where *aij* denotes the attention-derived features for nucleotide pair (*i*, *j*), s*_i_*^(0)^ and s*_j_*^(0)^ denote the corresponding single representations, and *erel* is a learned embedding of signed sequence separation. Relative sequence separations were clipped to the interval [−32, 32].

Single and pair representations were jointly processed by a 12-block Pairformer31 trunk with two recycling iterations. The resulting representations were then used to condition on an all-atom diffusion structure module31 comprising atom-level encoders and decoders and a 24-block diffusion transformer, which generated RNA atomic coordinates.

All RiboGRAM encoder parameters remained frozen throughout training. Trainable components comprised the single-and pair-representation projection layers, the SequenceToPair module, Pairformer trunk, diffusion structure module and auxiliary prediction heads. Training examples were cropped to a maximum length of 256 nucleotides. Contiguous sequence and spatial cropping were sampled with probabilities of 0.25 and 0.75, respectively. Optimization used AdamW with a peak learning rate of 1 × 10-4, a weight decay of 0.01 and 500 warm-up steps, followed by cosine learning-rate decay to 1 × 10-5.

The training objective combined the diffusion loss with auxiliary objectives for geometry, nucleotide interactions, secondary structure, and prediction confidence:

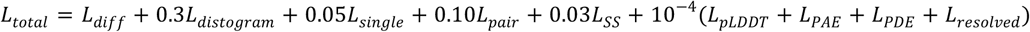

The diffusion loss was defined as:

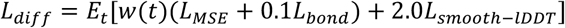

with

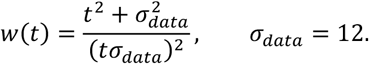

Here, *LMSE* quantified coordinate error between the denoised prediction and the experimentally determined structure after weighted rigid-body alignment, *Lbond* penalized deviations from covalent bond geometry, and *Lsmoot*ℎ−*lDDT* supervised local interatomic distance. The distogram, single-representation, pair-representation and secondary-structure losses provided auxiliary structural supervision. Separate confidence heads were trained for predicted local distance difference test (pLDDT), predicted aligned error (PAE), predicted distance error (PDE) and atom-resolution status. Confidence predictions were additionally used to rank independently generated diffusion samples.

### RNA tertiary structure dataset construction and benchmark evaluation

The RiboGRAM-3D training dataset was derived from the RNA tertiary structure collection released with DRfold233, with experimentally determined structures originating from the Protein Data Bank (PDB)32. Structures were processed in mmCIF format and separated into individual RNA chains. For each chain, nucleotide sequence and resolved atomic coordinates were extracted, and sequences were standardized to the RNA alphabet, with thymine converted to uracil where appropriate. Records lacking an identifiable RNA chain or valid nucleotide coordinates were excluded. Of the 10,432 RNA-chain records in the DRfold2 collection, 9,446 were converted for subsequent processing.

Following the temporal-split strategy used in DRfold2, the training set was restricted to structures released before 2024. To reduce sequence homology leakage, training RNAs sharing more than 80% sequence identity with any benchmark target were excluded. The held-out benchmark comprised 41 RNA chains ranging from 27 to 363 nt. Benchmark targets included recent RNA-Puzzles targets67, CASP15 RNA targets68 and RNA-only PDB structures released after the temporal cutoff and collected through 25 March 2025 (Supplementary Table 14).

For each benchmark target, RiboGRAM-3D generated 25 diffusion samples using independent initializations. Auxiliary confidence heads assigned model-derived ranking scores to the generated samples. When best-of-25 performance was reported, the sample with the lowest root-mean-square deviation (RMSD) to the experimentally determined structure was selected. The results were treated as an oracle assessment of sampling capacity and reported separately from confidence-ranked predictions.

RiboGRAM-3D was benchmarked against representative RNA tertiary structure prediction methods spanning fragment assembly, restraint-based modeling, and end-to-end learning approaches, including trRosettaRNA69, RNAComposer70, DeepFoldRNA71, RoseTTAFoldNA72, AlphaFold 331, RhoFold73 and DRfold233. All methods were evaluated on the same set of 41 benchmark targets using the official predictions reported by DRfold2.

Structural accuracy was evaluated using TM-score and RMSD. TM-score measured global structural similarity, with higher values indicating greater similarity to the experimentally determined structure74. For RMSD calculation, phosphate atoms from corresponding nucleotides were matched using the Kabsch algorithm. RMSD was calculated as:

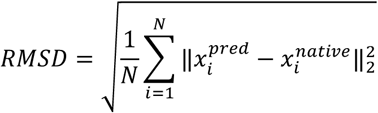

where *N* is the number of matched phosphate atoms, and *^pred^* and *x*^native^ denote the superimposed predicted and experimental coordinates of the *i*-th matched phosphate atom, respectively. Lower RMSD values indicate closer structural agreement.

For length-stratified evaluation, the 41 benchmark targets were divided into short (27–55 nt; n = 14), medium-length (57–118 nt; n = 15) and long (134–363 nt; n = 12) groups. For homolog-depth-stratified evaluation, MSAs were generated using the AlphaFold3 RNA data pipeline, which utilized nhmmer from the HMMER suite to search the NT-RNA, Rfam and RNAcentral databases. RNA MSA depth (*D*MSA) was defined for each target as the number of non-query homologous sequences in the resulting MSA. Targets for which no non-query homolog was detected (*D*MSA = 0; n = 14) formed the non-homolog group. The remaining 27 targets were ranked by *D*MSA and divided into low-depth (n = 13; observed *D*MSA = 1-11) and high-depth (n = 14; observed *D*MSA = 12-371) groups. These ranges reflect the observed MSA depths within the rank-defined groups rather than pre-specified biological thresholds. MSA information was used only for benchmark stratification and was not provided as input to RiboGRAM-3D. Performance was evaluated for individual targets and summarized across the benchmark using RMSD and TM-score distributions and corresponding summary statistics.

### RiboGRAM post-training for RNA–target interaction prediction

The pretrained RiboGRAM was post-trained for RNA–protein, RNA–RNA and RNA–chromatin interaction prediction using a shared concatenated-input framework. Each input began with a classification <CLS> token, followed by the RNA sequence and its interaction partner, with individual sequence segments terminated by <EOS> tokens. RNA–RNA inputs used the original RiboGRAM nucleotide vocabulary. For RNA–protein and RNA–chromatin inputs, the vocabulary was extended with amino-acid-and DNA-specific tokens, respectively. RNA– chromatin inputs additionally included both the forward DNA sequence and its reverse complement. Embeddings for newly introduced tokens were trainable during post-training.

All sequence segments within an interaction pair were processed jointly by the RiboGRAM encoder without modification of the encoder architecture. The final hidden representation of the <CLS> token was passed to a linear classification head to obtain a scalar interaction logit, which was converted to an interaction probability using a sigmoid function. The pretrained encoder, token embeddings and classification head were optimized end-to-end.

Post-training jointly optimized masked language modeling and binary interaction classification:

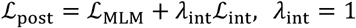

where ℒMLM denotes cross-entropy loss over selected masked-token positions and ℒint denotes binary cross-entropy loss for the interaction label. Approximately 15% of positions not occupied by special tokens were selected for masked-token prediction. Of these positions, 80% were replaced with the <MASK> token, 10% with a randomly selected token from the= vocabulary corresponding to the same sequence modality and 10% were left unchanged. For RNA–chromatin inputs, corresponding positions in the forward and reverse-complement DNA segments were masked jointly. Optimization used AdamW with a learning rate of 1 × 10-4.

Post-training was performed independently on the task-specific datasets. The resulting models are referred to as RiboGRAM–RPI, RiboGRAM–RRI and RiboGRAM–RCI for RNA–protein, RNA–RNA and RNA–chromatin interaction prediction, respectively. These models utilized the same core RiboGRAM encoder architecture and post-training objective while differing in partner modality, input composition and task-specific training data.

### RNA–protein interaction modeling

Two datasets were used for RNA–protein interaction post-training and evaluation (Supplementary Table 15). The primary dataset, RPI-Core, was derived from the processed RBP-binding data released with HDRNet34, which integrates POSTAR275 CLIP-seq and ENCODE40 eCLIP data. The released 101-nt positive and negative RNA sequences and their binary labels were retained and balanced with equal numbers. RPI-Core comprised 112 RBP-specific datasets from K562 cells, each partitioned into training, validation and test sets at a ratio of 7:1:2. Positive test sequences sharing at least 80% sequence identity with positive training sequences were removed using CD-HIT-EST-2D51, followed by class rebalancing. The resulting K562 training, validation and test sets contained 781,870, 111,694 and 144,576 RNA– protein pairs, respectively.

Cross-cell evaluation was performed on the 62 RBPs represented in both K562 and HepG2. For each RBP, HepG2 positive sequences sharing at least 80% sequence identity with the corresponding K562 positive training sequences were removed using the same filtering procedure, and the retained positives were combined with an equal number of negative sequence pairs. HepG2 RNA–protein sequence pairs were not used for model training, hyperparameter optimization or model selection. The resulting HepG2 evaluation set contained 523,712 RNA–protein pairs with equal representation of positive and negative classes.

To examine the effect of eCLIP data scale on post-training performance, an expanded dataset, RPI-Scale, was constructed from ENCODE40 eCLIP peaks for 138 RBPs in K562 and 105 RBPs in HepG2. Peaks from experimental replicates were pooled, overlapping intervals on the same chromosome and strand were merged, and each interval was converted into a peak-centered 101-nt positive sequence. Negative sequences were sampled from transcript regions, matched to the corresponding positive sequences by chromosome, strand and length, and excluded if located within 500 nt of any positive interval. Based on peak enrichment scores provided in the ENCODE eCLIP narrowPeak-format bigBed files, K562 subsets were constructed with per-RBP limits of 5,000, 50,000, 100,000 and 200,000 positive–negative pairs, together with an uncapped full set. The resulting K562 subsets were designated RPI-Scale-K562-5K, RPI-Scale-K562-50K, RPI-Scale-K562-100K, RPI-Scale-K562-200K and RPI-Scale-K562-All, respectively, and were used independently for continued post-training of RiboGRAM-RPI. For cell-line-specific motif analyses, the corresponding 100K subsets from K562 and HepG2 were referred to as RPI-Scale-K562-100K and RPI-Scale-HepG2-100K, respectively, and were used to construct attention-derived position weight matrices (PWMs) for RBP motif analysis.

The pretrained RiboGRAM model was first post-trained on the RPI-Core K562 training set to obtain RiboGRAM-RPI. Continued post-training was then performed independently on the RPI-Scale-K562-5K, RPI-Scale-K562-50K, RPI-Scale-K562-100K, RPI-Scale-K562-200K and RPI-Scale-K562-All subsets. The RPI-Scale-K562-100K variant was selected based on the highest accuracy and area under the receiver operating characteristic curve (AUC) and designated RiboGRAM-RPIplus for subsequent analyses. Baseline models included the sequence-based convolutional model DeepBind76, the self-attention-based SA-Net77, the sequence-and structure-aware PrismNet41, and HDRNet34 which combines contextual sequence and *in vivo* structure representations. All models were evaluated on the fixed RPI-Core K562 test set and, for the 62 RBPs shared between K562 and HepG2, on the independently held-out HepG2 evaluation set using AUC and accuracy.

### Attention-derived RBP motif discovery and peak-centered positional profiling

Attention-derived RBP motif analyses were performed using the RPI-Scale-K562-100K and RPI-Scale-HepG2-100K datasets, comprising 138 K562 and 105 HepG2 RBPs, respectively. For each RBP and cell type, up to 100,000 positive 101-nt RNA sequences centered on eCLIP peaks were retained; when fewer eligible peaks were available, all sequences were used. Each positive set was accompanied by an equal number of matched non-peak sequences. RNA sequences were oriented in the 5’-to-3’ direction according to the annotated strand and paired with the corresponding protein amino-acid sequence using the joint RNA–protein input format described above. During attention extraction and all subsequent motif analyses, the parameters of the post-trained RiboGRAM-RPIplus model were fixed.

For each RNA–RBP input, self-attention matrices were averaged across encoder layers and attention heads. Special token positions were excluded, and APC was applied to the resulting attention matrix to reduce row-and column-dependent background. APC-adjusted attention received by each RNA nucleotide was then aggregated across query positions to obtain a position-specific attention score. The 101 RNA-position scores were z-score normalized within each sequence. All overlapping 6-mers were assigned the mean standardized attention score across their six constituent nucleotides, and the highest-scoring 25% of 6-mer windows within each sequence were retained as candidate motif instances.

Candidate 6-mers were grouped when their Hamming distance was no greater than one. Within each resulting cluster, the nucleotide frequencies at the six aligned positions were aggregated across supporting RNA sequences to construct a 6-mer position PWM. Matched non-peak sequences were not used for PWM construction but were used to quantify enrichment of each candidate motif relative to background. Candidate PWMs were ranked according to their coverage among positive RNAs, enrichment relative to matched non-peak sequences and the associated statistical significance. A significantly peak-enriched candidate PWM was retained as the representative motif for each RBP.

Reference RBP motifs were obtained primarily from cell-type-matched records in mCrossBase35. When no corresponding mCrossBase motif was available, CisBP-RNA36, 37 was queried. RBPs without a motif in either resource were designated as lacking a reference motif. For RBPs with an available reference, the RiboGRAM-RPIplus-derived and reference PWMs were compared using TOMTOM37. Motif similarity was reported using the unadjusted raw TOMTOM *P* value.

For peak-centered positional profiling, the representative PWM for each RBP was used to scan the corresponding positive and matched non-peak 101-nt sequences. For each sequence, the three highest-scoring non-overlapping motif windows were retained. At each nucleotide position, motif occupancy was defined as the fraction of sequences for which at least one retained motif window overlapped that position. The positional occupancy profile for each RBP was calculated as motif occupancy in positive sequences minus that in matched non-peak sequences at each position across the 101-nt window.

### RNA–RNA interaction modeling

RNA–RNA interaction prediction was evaluated using the MirTarRAW78, 79 benchmark and a RISE80-derived dataset comprising 15 RNA-type combinations (Supplementary Table 16). MirTarRAW78 combines DeepMirTar79 and miRAW81 and contains 27,720 miRNA-mRNA pairs, with equal numbers of positive and negative examples. Label-stratified partitioning at a ratio of 1:1:8 yielded 2,772 training, 2,772 validation and 22,176 test pairs. For the RISE-derived dataset, experimentally supported human RNA–RNA interactions were used as positive pairs, with both interacting regions restricted to less than 500 nt. For each positive pair, up to four background-matched negative candidates were generated by retaining one RNA and replacing its partner with equal-length, same-chromosome and same-strand flanking segments, separated from the original interval by exactly 500 nt. Positive pairs were stratified by RNA-interaction class and partitioned into training, validation and test sets at an approximate ratio of 7:1:2, with negative pairs generated independently within each partition. Pairs overlapping those represented in the training partition were removed from the validation and test partitions, and the test sequences were additionally filtered using an 80% sequence identity threshold to reduce redundancy. The resulting training, validation and test sets contained 261,997, 34,184 and 28,551 RNA pairs, respectively.

RiboGRAM was post-trained separately on the MirTarRAW and RISE-derived datasets, generating two dataset-specific RiboGRAM-RRI models for binary RNA–RNA interaction prediction. For the RISE-derived dataset, each training pair was presented in both RNA orders as data augmentation. Masked language modeling and binary interaction classification objectives were optimized jointly with equal weighting, with approximately 15% of nucleotide positions selected for masking during training. RNAErnie2, RNA-FM and RiNALMo were included as comparison models. For each model, a linear classification head was added to the pretrained encoder, and the encoder and classification head were fine-tuned end-to-end using a model-specific learning rate. Performance was evaluated using the area under the receiver operating characteristic curve (AUC) and accuracy.

### RNA–chromatin interaction modeling

Six human RNA–DNA contact datasets curated in the RNA-Chrom database82 were used to evaluate RNA–chromatin interaction prediction (Supplementary Table 17). These comprised iMARGI datasets from HEK293T and HUVEC cells, Red-C datasets from fibroblasts and K562 cells, and GRID-seq datasets from MDA-MB-231 and MM.1S cells. Experimentally detected RNA–DNA contacts were treated as positive pairs; these labels represent assay-defined associations and do not distinguish direct RNA–DNA contacts from interactions mediated by proteins or other chromatin components. For each positive contact, two candidate negative pairs were generated while retaining the interacting DNA interval. The original RNA interval was replaced with an equal-length genomic fragment on the same strand, located either upstream or downstream of the original RNA interval, with a gap of at least 500 bp between the two intervals. Negative examples were sampled from these locally generated candidates to construct class-balanced datasets.

To construct evaluation sets enriched for strongly supported contacts, positive contacts within each dataset were independently ranked by the background-normalized contact strengths reported by RNA-Chrom. For the HEK293T dataset, the 100,000 highest-ranked positive contacts and 100,000 constructed negatives were assigned to the test set. The next 100,000 positive contacts and 100,000 constructed negatives were assigned to the validation set, and the remaining 4.8 million positive contacts and 4.8 million negative pairs were used for training. The other five datasets were reserved for external evaluation across cell types and methods, with each evaluation set comprising the 100,000 highest-ranked positive contacts and 100,000 constructed negatives.

Each RNA–DNA pair was encoded using the shared concatenated-input framework described above. The RNA sequence was oriented according to its strand and combined with the interacting DNA sequence in reference-genome orientation and its reverse complement. A sequence-level classification token was added at the beginning of the input, and boundary tokens were inserted to distinguish the three sequence components. The RiboGRAM vocabulary was extended with DNA-specific nucleotide tokens, allowing RNA and DNA nucleotides to be represented by distinct token sets. RiboGRAM was subsequently post-trained on the HEK293T training set for binary RNA–chromatin interaction classification to obtain the RiboGRAM-RCI model. Masked language modeling and binary interaction classification objectives were optimized jointly with equal weighting. Approximately 15% of RNA and DNA nucleotide positions were selected for masking. Corresponding positions in the reference-orientation and reverse-complement DNA representations were masked jointly. Model training was restricted to the HEK293T training set. The HEK293T validation set was used for model validation, whereas the HEK293T test set and the five external evaluation sets were held out from training and model selection. The sequence-only configuration of DeepRCI83, an interpretable deep learning framework for modeling variable-length RNA and DNA sequences, was used as the comparison method. Performance was evaluated using AUC and accuracy.

## Data availability

All datasets compiled and curated for this study are publicly accessible from the Zenodo repository (https://zenodo.org/records/23201637) and the RiboGRAM resource portal (https://www.ultrarnalab.com/ribogram.php).

## Code availability

The source code and analysis scripts are publicly accessible through the Zenodo repository (https://zenodo.org/records/23201637) and mirrored on the RiboGRAM resource portal at https://www.ultrarnalab.com/ribogram.php. The executable Code Ocean capsule is provided (https://codeocean.com/capsule/2602506/tree/v1) to facilitate reproduction. RiboGRAM is also freely available as a web server at https://www.ultrarnalab.com/ribogram_usage.php.

## Author contributions

Y.Z. and H.W. conceptualized the study. H.W. and Y.Z. performed the modeling and data analyses, with H.H. performing the three-dimensional structure prediction analyses. Y.Z., Z.C., H.W., Q.Y. and A.J. contributed to data interpretation. Y.Z. and Q.Y. supervised this study. H.W. and Y.Z. wrote the original draft. All authors discussed the results and revised the manuscript.

## Acknowledgements

We thank the BAAI-Health Center and the Jäschke lab members for discussions, P. González Jabalera (Heidelberg University) for helpful comments, Y. Wu (Heidelberg University) for figure preparation, BAAI-JIUDING for GPU cluster computing resources, bwForCluster Helix, bwUniCluster 3.0 and HoreKa at NHR@KIT for CPU computing resources and the SDS@hd service for data storage.

## Competing Interests

The authors declare no competing interests.

## Supplementary Materials

**Supplementary Fig. 1.**
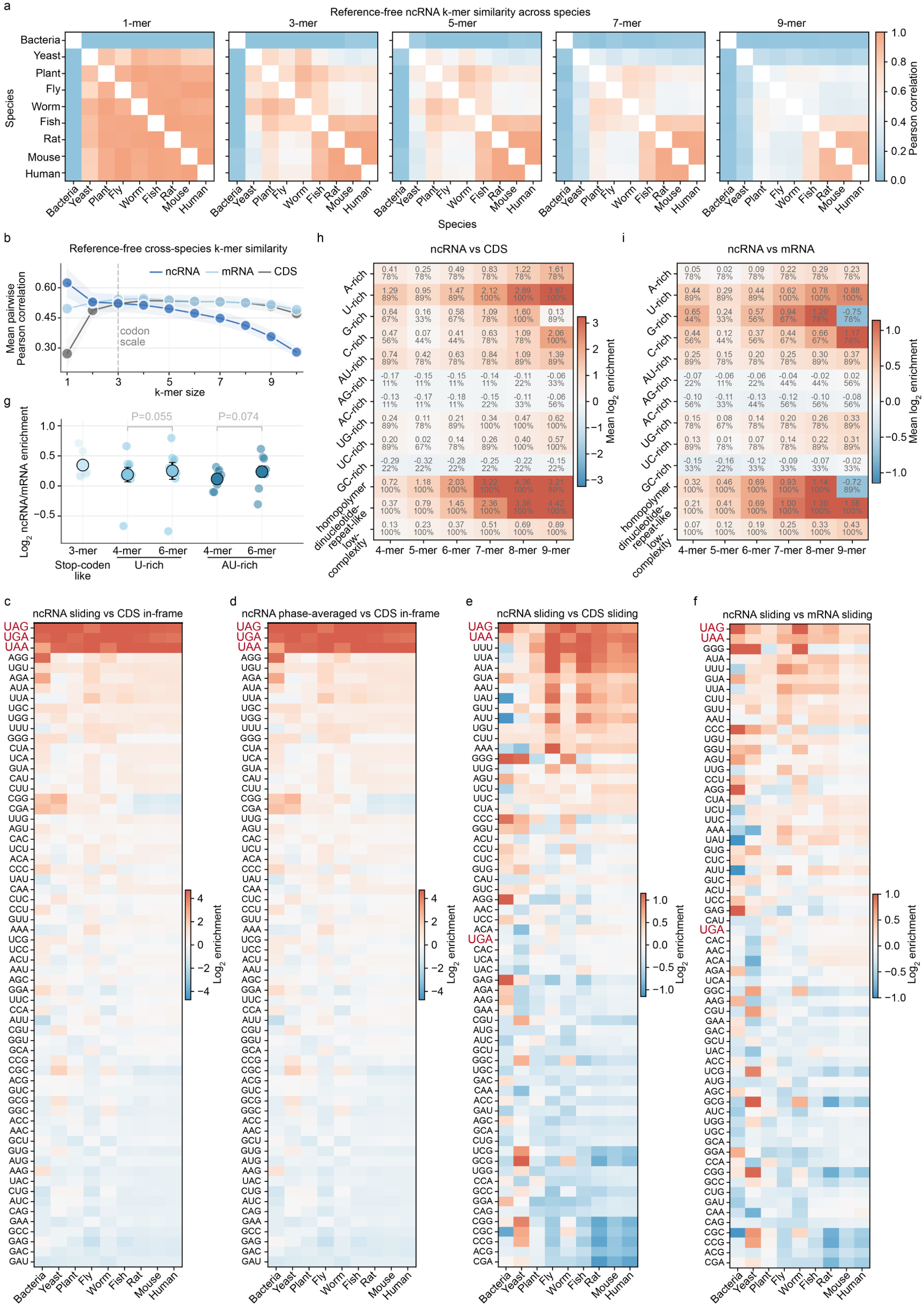
Multi-scale organization of ncRNA local sequence patterns across species. a, Pairwise ncRNA k-mer profile similarity across nine model organisms. For each k-mer size of 1, 3, 5, 7 and 9 nt, species-level k-mer counts were normalized as log2 (CPM +1), where CPM denotes counts per million, and Pearson correlations were calculated between each pair of species. Diagonal self-comparisons are masked. **b,** Cross-species k-mer profile similarity for ncRNA, mRNA and CDS. For each RNA type and k-mer size from 1 to 10 nt, Pearson correlations were calculated between all 36 species pairs using the species-level profile vector defined in panel a. Points show the mean pairwise correlation, and shaded bands show s.e.m. across species pairs. The dashed vertical line marks the 3-mer scale. **c-f**, ncRNA triplet enrichment across counting frames and reference RNA types. For each species, the frequencies of all 64 triplets were calculated separately for ncRNA and the indicated CDS or mRNA reference. Enrichment was defined as the log2 ratio of ncRNA to reference frequency, with a small constant added to both frequencies to accommodate zero values. In panel **c**, overlapping ncRNA triplets were enumerated using a 3-nt sliding window with a one-nucleotide step and compared with in-frame codons from annotated CDS. In panel **d**, non-overlapping ncRNA triplet frequencies were calculated at each of the three possible offsets and averaged to generate a phase-averaged pseudo-frame profile, which was compared with in-frame CDS codons. In panels e and f, the same 3-nt sliding-window procedure was applied to both sides of each comparison, with ncRNA compared against CDS in panel **e** and mRNA in panel **f**. The stop codon triplets UAA, UAG, and UGA are highlighted in red. **g,** ncRNA-enriched local sequence patterns relative to mRNA. Selected stop-like triplets and U-rich and AU-rich 4-and 6-mer classes are shown as log2 enrichment relative to same-species mRNA. Colored dots denote species-level values, and black circles and error bars show cross-species mean ± s.e.m. across nine species. Brackets indicate paired 4-versus 6-mer comparisons within the U-rich and AU-rich classes; two-sided Wilcoxon signed-rank test; n = 9. **h,i**, Multi-scale enrichment of local sequence-pattern classes in ncRNA. For each species and k-mer size from 4 to 9 nt, the frequency of each class was obtained by summing the frequencies of all k-mers assigned to that class. Enrichment was calculated as the log2 ratio of the ncRNA class frequency to the corresponding same-species CDS frequency in panel **h** or mRNA frequency in panel **i**. A-, U-, G-, and C-rich classes comprised k-mers in which the indicated nucleotide accounted for at least 80% of positions. AU-, AG-, AC-, UG-, UC-and GC-rich classes comprised k-mers in which the indicated nucleotide pair jointly accounted for at least 80% of positions, excluding k-mers already assigned to a single-nucleotide-rich class. Homopolymer, dinucleotide-repeat-like and low-complexity classes comprised k-mers containing one repeated nucleotide, a period-2 repeat or no more than two distinct nucleotides, respectively. These three classes were defined independently of the nucleotide-rich classes and were therefore not mutually exclusive. Cell color shows mean log2 enrichment across nine species. Cell annotations report the mean enrichment and the percentage of species with positive enrichment.

**Supplementary Fig. 2.**
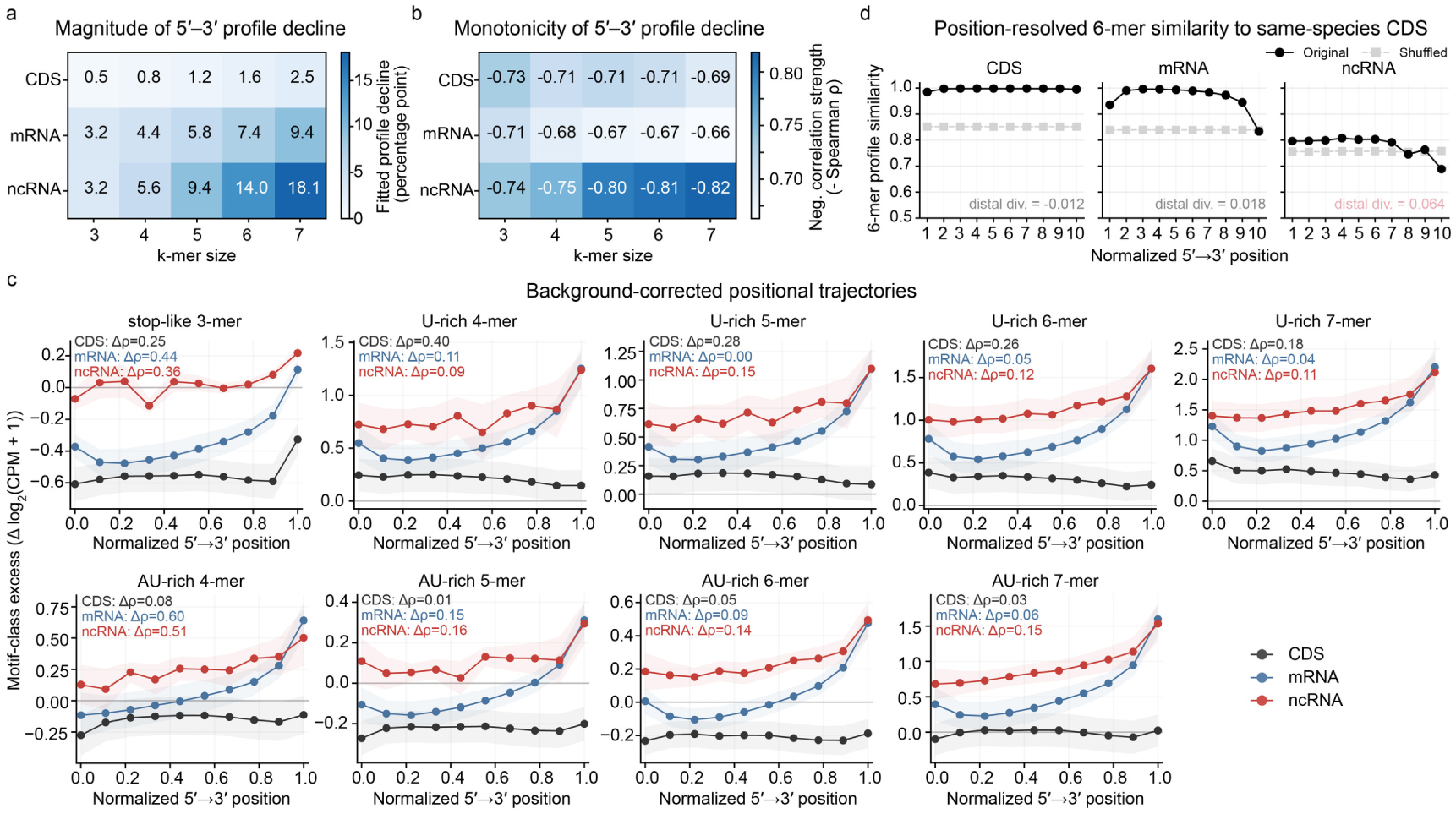
Position-resolved directional organization of ncRNA sequence patterns along the 5’-to-3’ axis. a,b, Magnitude and monotonicity of position-resolved k-mer profile divergence. For each species and RNA type, RNAs of at least 60 nt were divided into ten length-normalized 5’-to-3’ bins. Each bin was represented by its k-mer frequency profile for k-mer sizes from 3 to 7 nt, and cosine similarity to the bin-1 profile from the same species and RNA type was calculated. **a,** Profile decline from bin 1 to bin 10 was estimated from the linear fit across bins. **b,** Positional monotonicity was quantified by the Spearman correlation between bin position and profile similarity. Cells show means across nine species; more negative ρ values indicate a stronger monotonic decline toward the 3’-end. **c,** Background-corrected positional trajectories of selected motif classes. For each species, RNA type and position bin, motif-class excess was calculated as the mean log2 (CPM + 1) of class k-mers minus that of composition-matched background k-mers. Stop-like 3-mers comprised UAA, UAG and UGA. U-rich and AU-rich classes were defined using the prespecified k-specific composition criteria, with U-rich k-mers excluded from the corresponding AU-rich classes. Lines and shaded bands show mean ± s.e.m. across nine species. Annotations report matched-background Spearman excess, Δρ, defined as the positional Spearman correlation ρ of the motif class minus that of its matched background. Positive values indicate a stronger increase toward the 3’-end than the matched background. **d,** Position-resolved 6-mer profile similarity to same-species CDS. Using the same positional binning in panel a, each bin-specific 6-mer frequency profile was compared by cosine similarity with a fixed same-species CDS profile pooled across all CDS bins. Solid and dashed lines show means across nine species for original sequences and per-sequence shuffled controls, respectively. Lower-right annotations report the mean distal divergence of original sequences, defined within each species as the similarity of bin 1 minus the mean similarity across bins 8-10 and then averaged across species.

**Supplementary Fig. 3.**
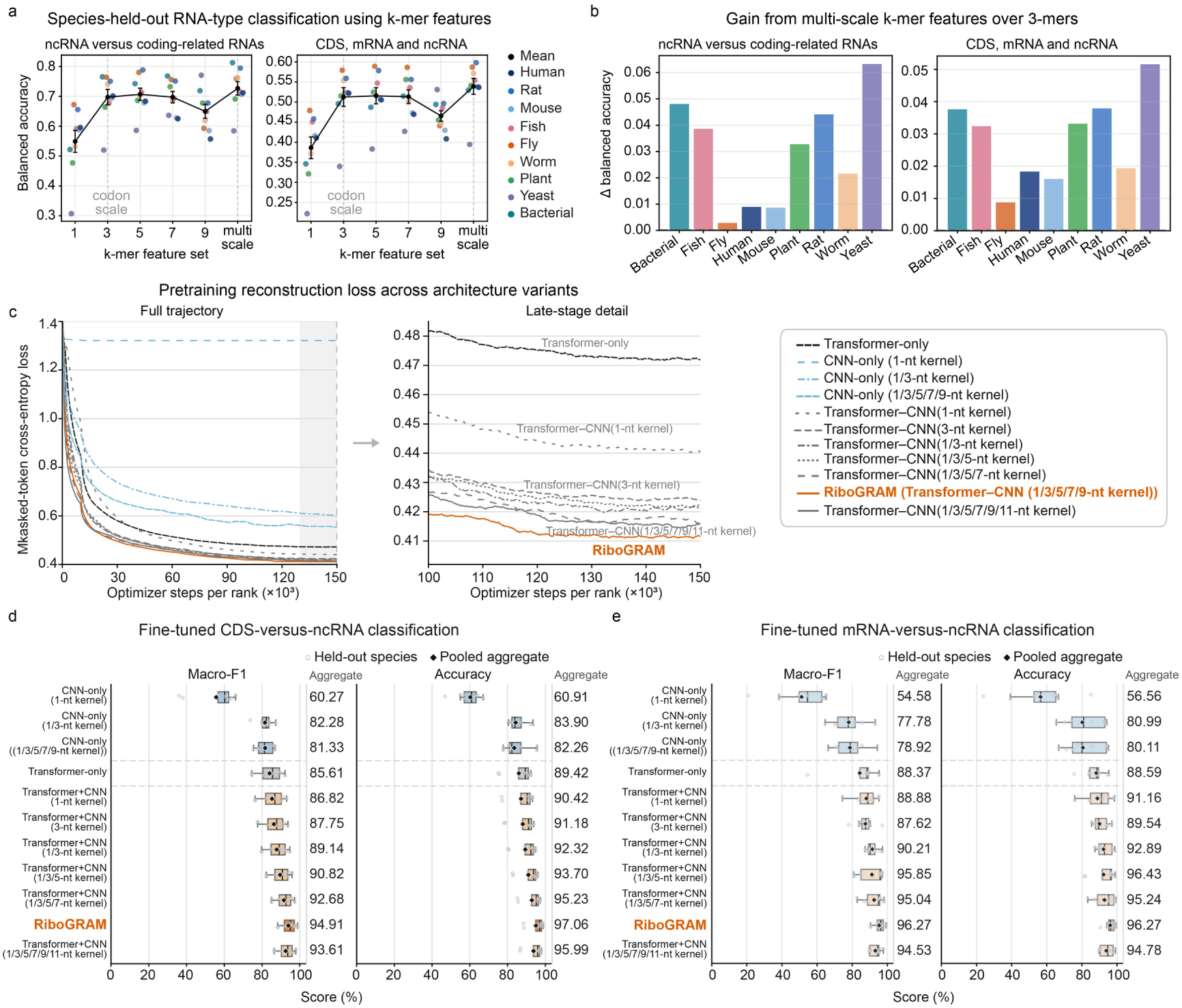
Cross-species RNA-type classification and model optimization analyses. a, Species-held-out RNA-type classification from k-mer composition. Linear classifiers with logistic loss were trained on sequences from eight species and evaluated on the remaining species in a leave-one-species-out design. Inputs comprised per-sequence frequencies of 1-, 3-, 5-, 7-, or 9-mers, or the concatenated 1/3/5/7/9-mer feature set, with each k-mer block normalized separately. Left, binary classification distinguished ncRNA from a coding-related group containing equal numbers of CDS and mRNA sequences and matched in total sample size to ncRNA. Right, three-class classification distinguished CDS, mRNA and ncRNA using equal class sampling within each species. Colored points show the mean balanced accuracy for each held-out species across 20 sequence-resampling runs, and black points and lines show means across the nine held-out species. **b,** Species-level gain from multi-scale k-mer features over 3-mer alone. Bars show, for each held-out species, the difference in balanced accuracy between the concatenated 1/3/5/7/9-mer feature set and the 3-mer feature set for binary classification of ncRNA versus a coding-related class composed of equal numbers of CDS and mRNA sequences (left) and for three-class classification of CDS, mRNA and ncRNA (right). Positive values indicate improved performance with the multi-scale feature set. **c,** Pretraining loss across architecture variants. Masked-token reconstruction loss was monitored during pretraining for the indicated Transformer-only, CNN-only and hybrid Transformer–CNN architectures. Left, complete optimization trajectories as a function of optimizer steps per rank; right, late-stage trajectories shown on an expanded scale. The full RiboGRAM architecture, which integrated Transformer-based contextual modeling with convolutional kernels of 1-, 3-, 5-, 7-, and 9-nt, is highlighted in orange. **d,e,** Frozen-encoder RNA-type classification across architecture variants. Each pretrained architecture in panel c was evaluated using the same frozen-encoder procedure for binary classification of ncRNA versus CDS (panel **d**) or mRNA (panel **e**). Encoder parameters were held fixed, and only the identical MLP classifier was trained. Macro-F1 and accuracy are reported for each model. Boxplots summarize performance across nine species, with points denoting individual species, and the aggregate columns report the corresponding median scores across species.

**Supplementary Fig. 4.**
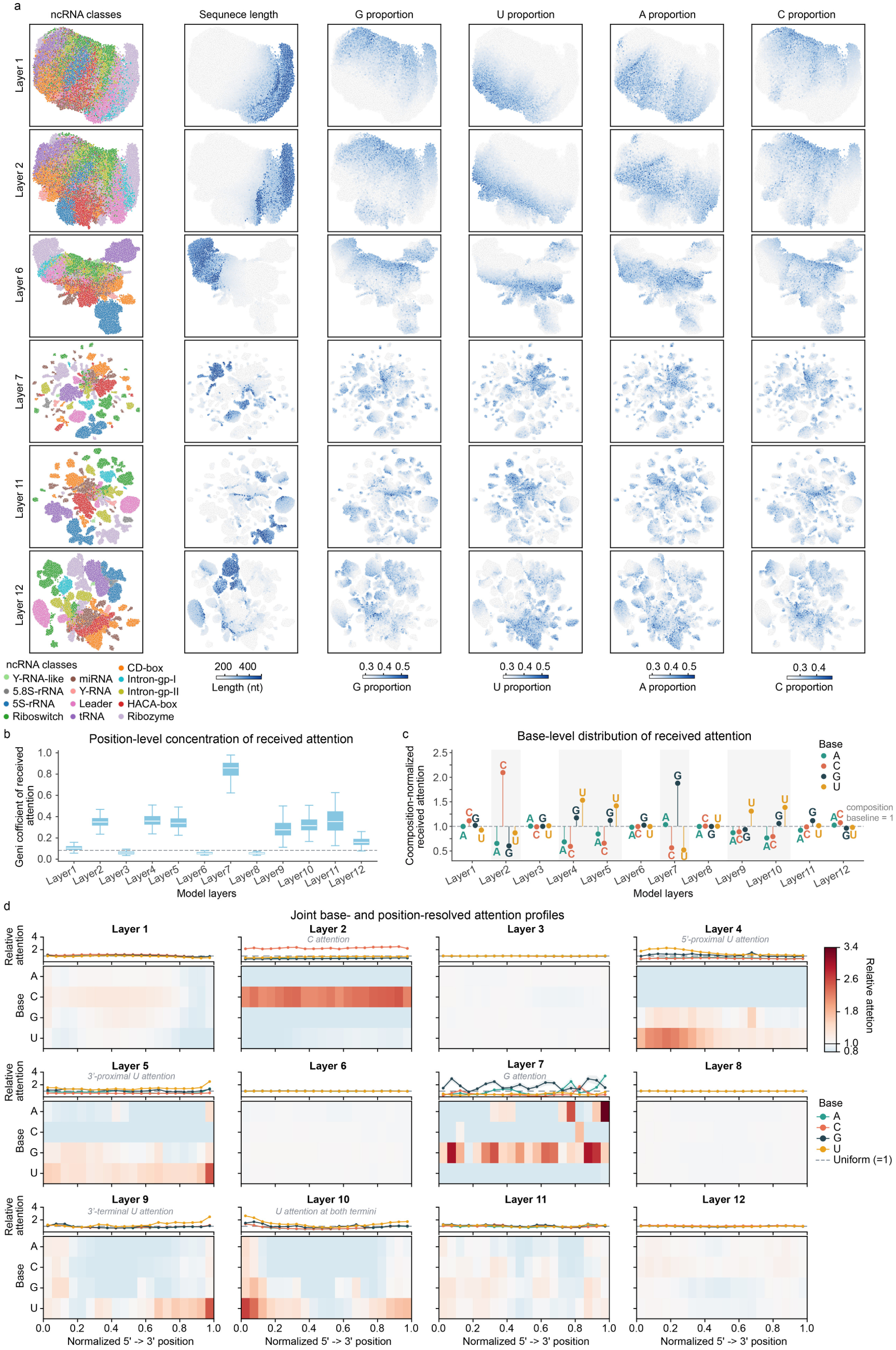
RiboGRAM layers reveal ncRNA class organization and distinct attention patterns. a, Layer-resolved UMAPs of mean-pooled sequence embeddings colored by ncRNA category and sequence features. The analysis includes all 45,447 sequences in the 13-class NCYPred dataset1 and six representative layers spanning early, intermediate, and late RiboGRAM representations. For each layer, the same UMAP coordinates are colored by ncRNA class, sequence length, or the sequence-wide proportions of G, U, A and C. For each feature, the color scale is fixed across layers. **b,** Position-level concentration of received attention across RiboGRAM layers. Following head-averaging, APC-adjustment and row normalization, the attention received at each nucleotide position was summed across query positions, and its concentration was quantified using the Gini coefficient. High values indicate attention concentrated on fewer positions. The analysis used a class-balanced set of 1,300 sequences, comprising 100 sequences from each ncRNA class. Boxes show the median and interquartile range, whiskers extend to 1.5 times the interquartile range, and outliers are not shown. **c,** Base-level distribution of received attention across RiboGRAM layers. For each base, the total attention received by the corresponding positions was normalized by its sequence-level abundance. A value of one indicates attention proportional to nucleotide composition, whereas values above and below one indicate greater or lower received attention than expected from composition alone. Letters show layer-wise means, and vertical lines connect each mean to the composition-proportional baseline of 1. Shaded bands mark selected layers with prominent deviations from this baseline. **d,** Joint base-and position-resolved profiles of received attention across RiboGRAM layers. Using the same class-balanced sequence set as in panels b and c, each sequence was divided into 20 length-normalized 5’-to-3’ bins, and received attention to A, C, G and U positions within each bin was normalized to the uniform per-position expectation. A value of one denotes uniform attention. Upper line plots show mean base-specific trajectories, and lower heat maps show the corresponding base-by-position pattern across layers. Annotations highlight prominent patterns in selected layers.

**Supplementary Fig. 5.**
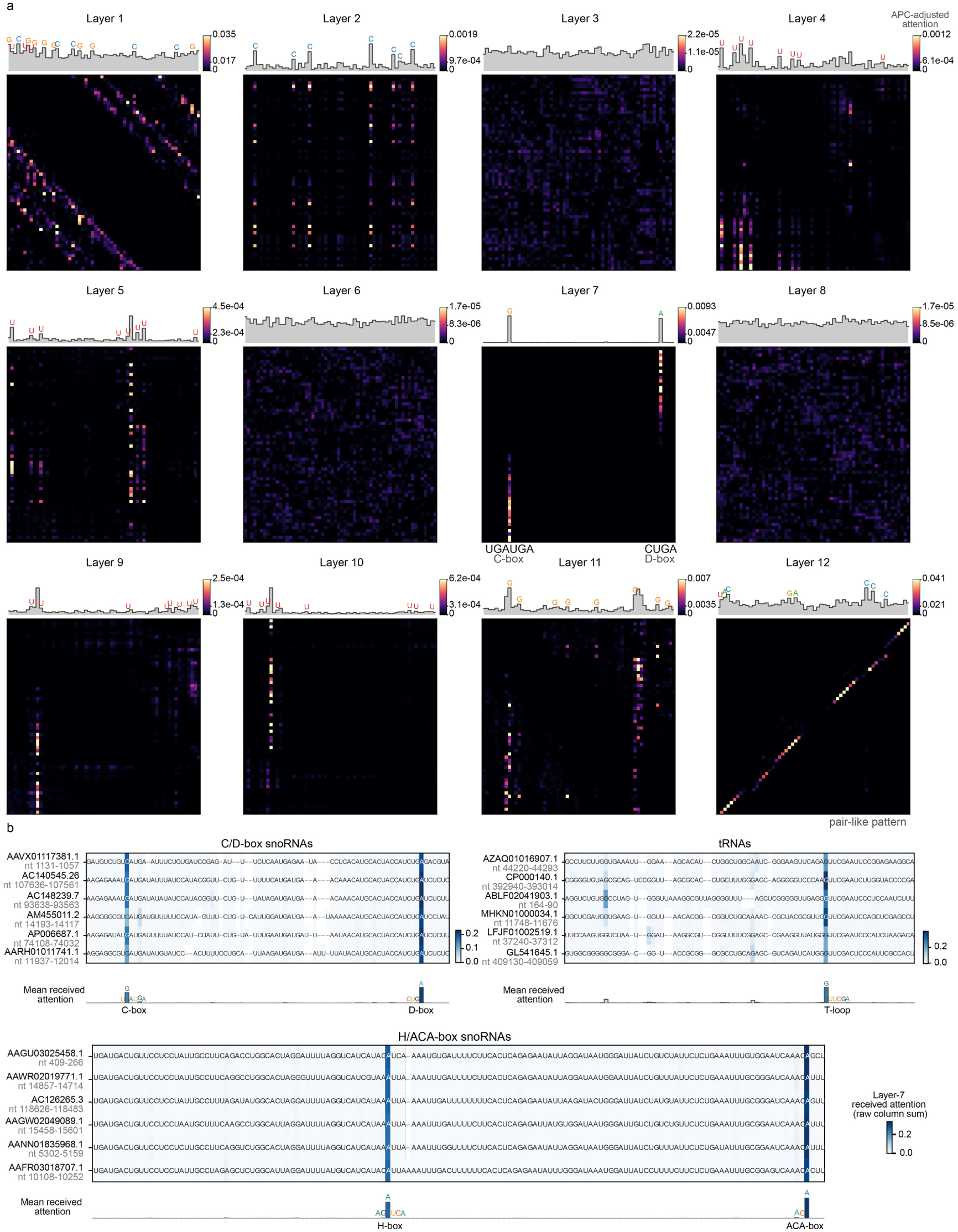
Layer-resolved RiboGRAM attention reveals motif-centered and pair-like organization. a, Layer-resolved self-attention patterns in the representative C/D-box snoRNA snoZ39 (AJ2400571.1, nt 1-60). Head-averaged APC-adjusted attention matrices are shown across all 12 RiboGRAM layers, with the profile above each matrix summarizing the total attention received at each nucleotide position. Prominent nucleotide positions are labeled in the profiles, and the canonical C-box (UGAUGA) and D-box (CUGA) are indicated below the layer-7 matrix. **b,** Cross-sequence organization of layer-7 attention relative to annotated conserved elements in C/D-box snoRNAs, tRNAs, and H/ACA-box snoRNAs. Within each RNA group, aligned sequences are displayed together with the annotated motif regions, with alignment gaps shown where required. Blue intensity represents the raw column-summed layer-7 attention received at each aligned nucleotide position, and the profile below each alignment summarizes the mean of this quantity at each alignment position. The C-and D-boxes, tRNA T-loop, and H-and ACA-boxes are indicated below the corresponding alignments.

**Supplementary Fig. 6.**
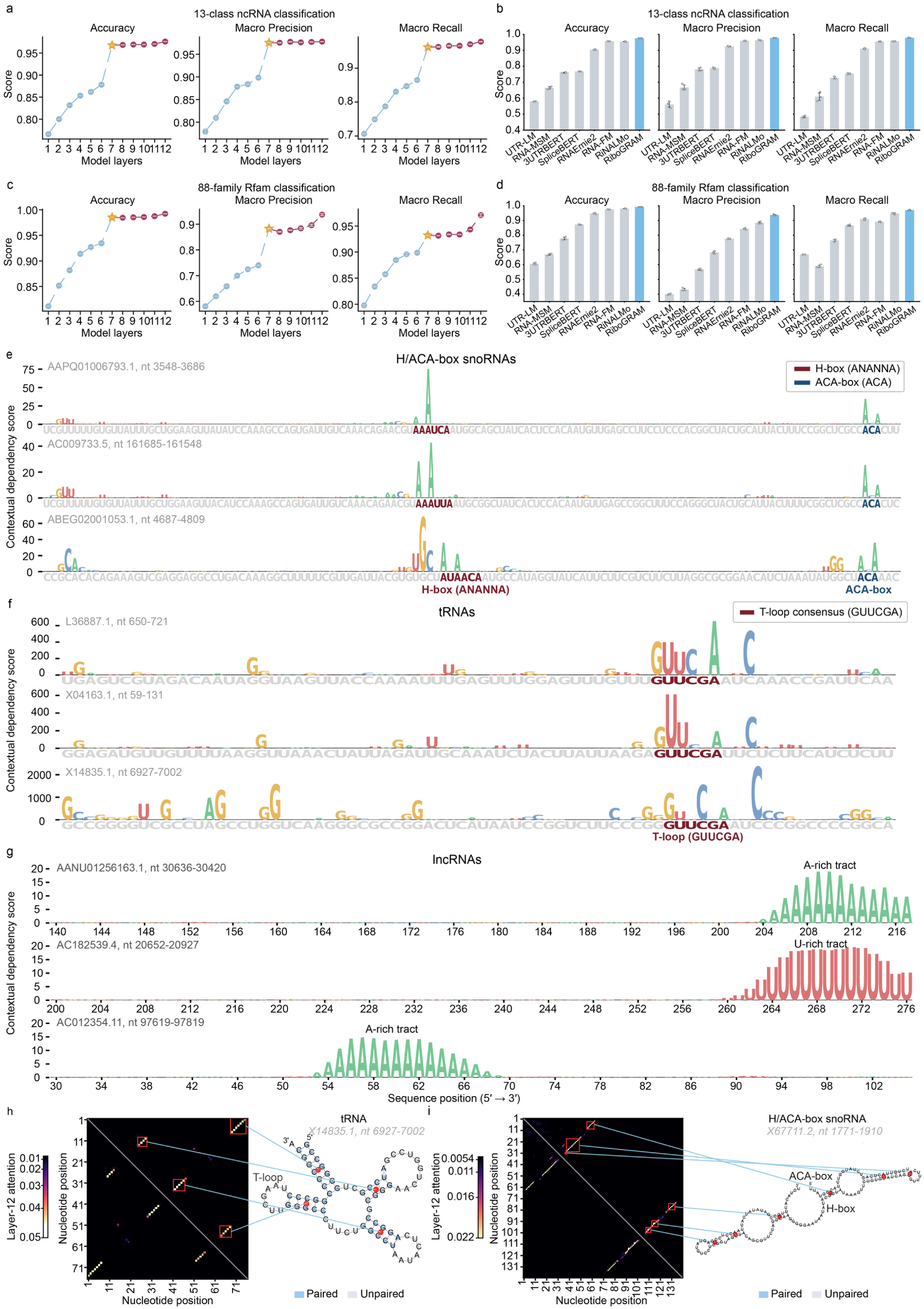
RiboGRAM representations resolve ncRNA identity and reveal local sequence features and pair-like signals. a, Layer-wise linear-probe performance for 13-class ncRNA classification. Linear probes were trained separately on sequence embeddings from each RiboGRAM layer and evaluated on the held-out NCYPred1 test set of 13,646 sequences. Accuracy, macro precision and recall are shown as mean ± s.d., with n = 3 random seeds. **b,** Cross-model comparison of layer-12 representations for 13-class ncRNA classification. Using the same linear probing as in panel a, RiboGRAM was compared with the indicated RNA language models on the held-out NCYPred1 test set. Accuracy, macro precision and recall are shown as mean ± s.d., with n = 3 random seeds. **c,** Layer-wise linear-probe performance for 88-family Rfam classification2. Linear probes were trained separately on sequence embeddings from each RiboGRAM layer and evaluated on the held-out Rfam test set of 25,342 sequences. Accuracy, macro precision and recall are shown as mean ± s.d., with n = 3 random seeds. **d,** Cross-model comparison of layer-12 representations for 88-family Rfam classification2. Using the same linear probing as in panel c, RiboGRAM was compared with the indicated RNA language models on the held-out Rfam test set. Accuracy, macro precision and recall are shown as mean ± s.d., with n = 3 random seeds. **e-g,** Position-resolved contextual dependency across H/ACA-box snoRNAs (**e**), tRNAs (**f**) and lncRNAs (**g**). Letter height denotes the nucleotide-level contextual dependency score, with higher values reflecting stronger and more confident recovery of the masked nucleotide from its surrounding sequence context. Annotated H-box, ACA-box regions and T-loop elements are indicated, together with the selected high-scoring A-or U-rich low-complexity tracts in the lncRNA examples. **h,i,** Layer-12 pair-like attention relative to Rfam-annotated secondary structures of the tRNA X14845.1 (nt 6927–7002; **h**) and H/ACA-box snoRNA X6711.2 (nt 1771–1910; **i**). Symmetrized, APC-adjusted attention maps are shown alongside the corresponding secondary-structure diagrams. Red boxes mark selected attention-map regions, with blue connectors indicating the corresponding annotated base-paired regions in the structure diagrams. The tRNA T-loop and snoRNA H-and ACA-boxes are annotated for reference.

**Supplementary Fig. 7.**
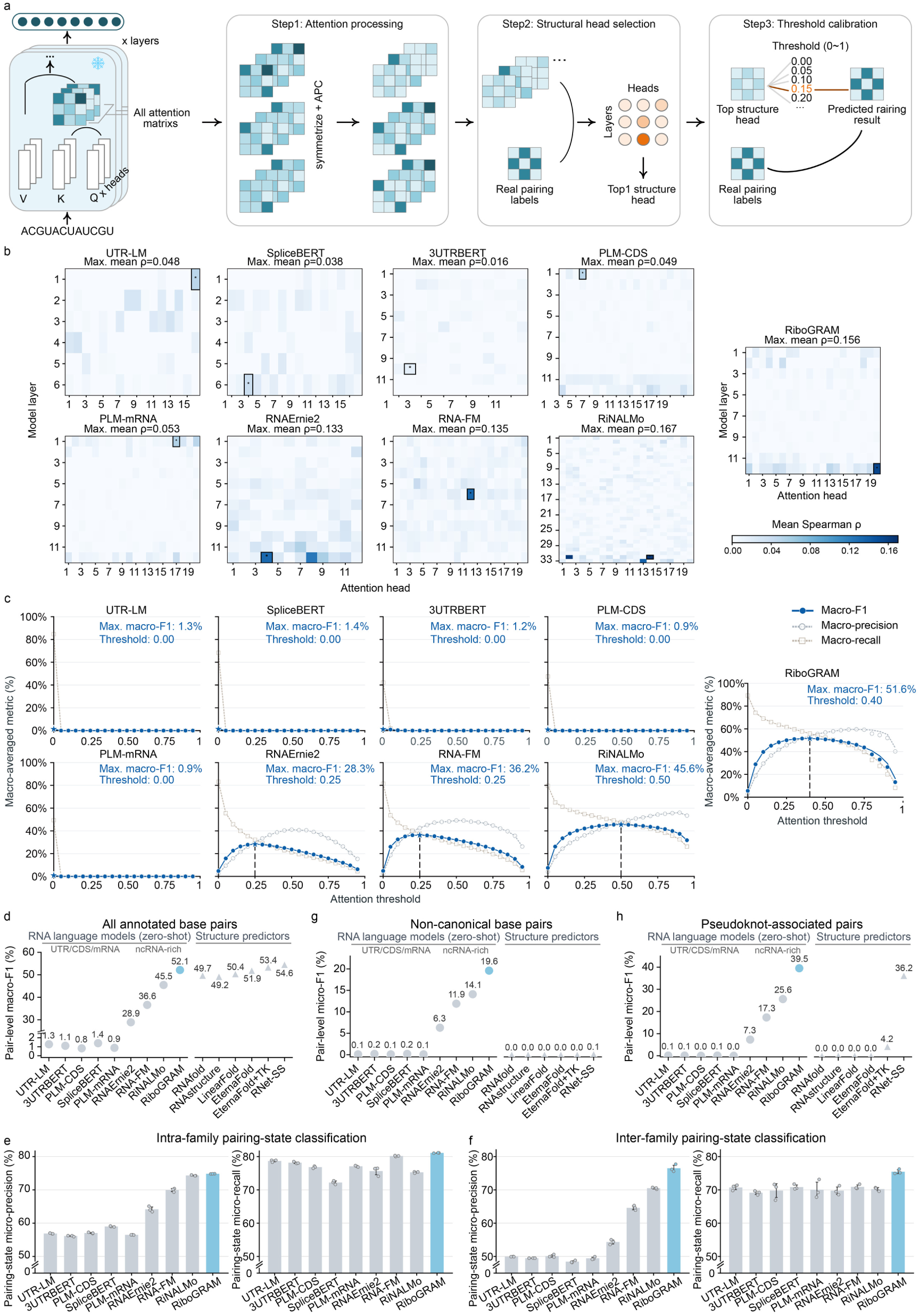
ncRNA-rich pretraining strengthens recoverable base-pairing signals. a, Training-free extraction of base-pairing signals from model self-attention. Attention maps from all layers and heads were symmetrized and APC-adjusted. On bpRNA-1m VL03, the layer-head combination with the highest mean sequence-level Spearman ρ to annotated base-pair maps was selected for each model, and the attention threshold maximizing macro-F1 was fixed before evaluation on held-out data. **b,** Mean attention-pairing Spearman ρ across layers and heads on bpRNA-1m VL0^3^ (n = 1,300 sequences). Black boxes mark the selected heads. **c,** Threshold dependence of macro-precision, macro-recall and macro-F1 on the same bpRNA-1m VL03 set in panel b. Dashed line and annotations indicate the selected thresholds and corresponding peak macro-F1. **d,** Held-out zero-shot macro-F1 on bpRNA-1m TR0^3^ (n = 10,814), using the head and threshold fixed on VL0 for each model. RNA language models are grouped by pretraining corpus, with secondary structure predictors shown for comparison. EternaFold+TK denotes EternaFold with ThreshKnot decoding^4, 5^ and RNet-SS is structure-supervised^6, 7^. Complete micro-and macro-averaged precision, recall and F1 scores are provided in Supplementary Table 8. **e,f,** Precision and recall for the intra-family (**e**) and the inter-family (**f**) nucleotide pairing-state classification, corresponding to the F1 results in Fig. 3b,c, respectively. Identically configured two-layer MLP probes were trained on frozen model embeddings using the bpRNA-1m TR0/VL0/TS03 splits for intra-family evaluation and the TrainSetA-to-TestSetB^8^ protocol for inter-family evaluation. TrainSetA contains eight RNA families, while TestSetB contains 22 unseen families with less than 70% sequence identity to TrainSetA. Bars show mean ± s.d. across three random seeds (n = 3). **g,** Zero-shot micro-F1 for non-canonical base pairs on bpRNA-1m TR0, defined as annotated pair types other than A-U, G-C, and G-U pairs, using the same model set as in panel d. **h,** Zero-shot micro-F1 for pseudoknot-associated crossing base pairs, evaluated on the 999 TR0 sequences containing at least one annotated crossing pair, using the same model set in panel d. Corresponding micro-precision, micro-recall and micro-F1, together with complete metrics for canonical and wobble base pairs, non-canonical and pseudoknot-associated pairs are provided in Supplementary Table 9.

**Supplementary Fig. 8.**
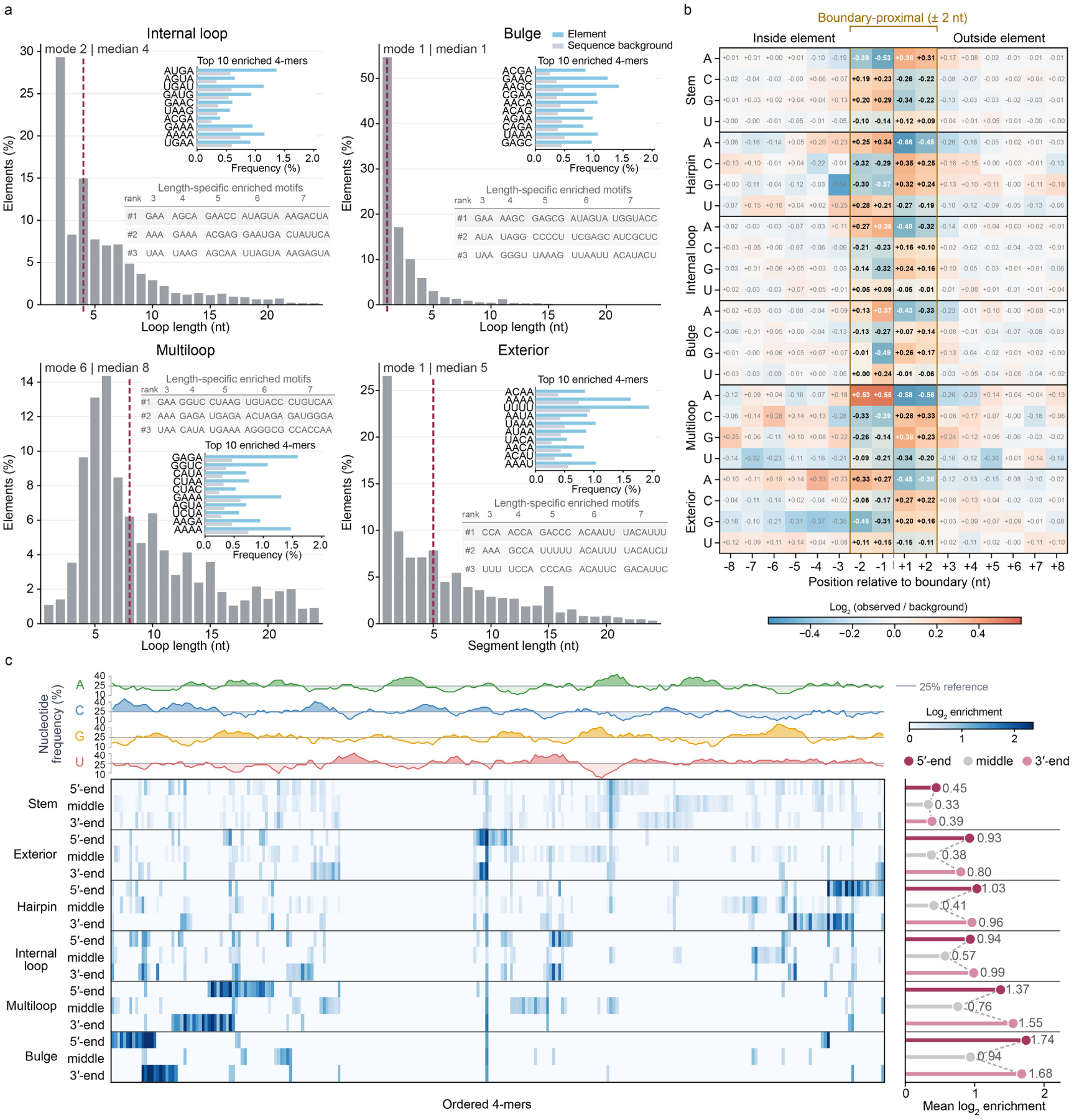
RNA structural elements show length-and position-resolved sequence patterns. a, Length-scale organization and local sequence patterns of annotated internal loops, bulges, multiloops and exterior regions. Histograms show the proportion of structural elements at each length with mode and median indicated. Insets compare the frequencies of the ten most enriched 4-mers within each structural element with their global frequencies across all full-length sequences. Lower tables show the three highest-ranked enriched motifs at element lengths of 3–7 nt. **b,** Position-resolved nucleotide patterns across structural element boundaries. Cells show relative nucleotide enrichment, quantified as log2 (observed/background) for A, C, G and U at positions-8 to-1 inside and +1 to +8 outside each structural element. The 5’ and 3’ boundaries were oriented to a common inside-to-outside coordinate before aggregation, with positions within ±2 nt of the boundary highlighted. **c,** 5’-to-3’ variation in 4-mer patterns within structural elements. Rows represent the 5’-end, middle and 3’-end regions of six structural elements, and columns show all 256 4-mers, ordered by hierarchical clustering of their enrichment profiles. Colored cells indicate positive log2 enrichment relative to whole-sequence background with Benjamini-Hochberg-adjusted q < 0.05 from one-sided binomial tests. Top profiles show A, C, G and U frequencies across the 4-mer order, with 25% marking the equal-base reference. Summary plots alongside the heatmap show mean log2 enrichment for each element region.

**Supplementary Fig. 9.**
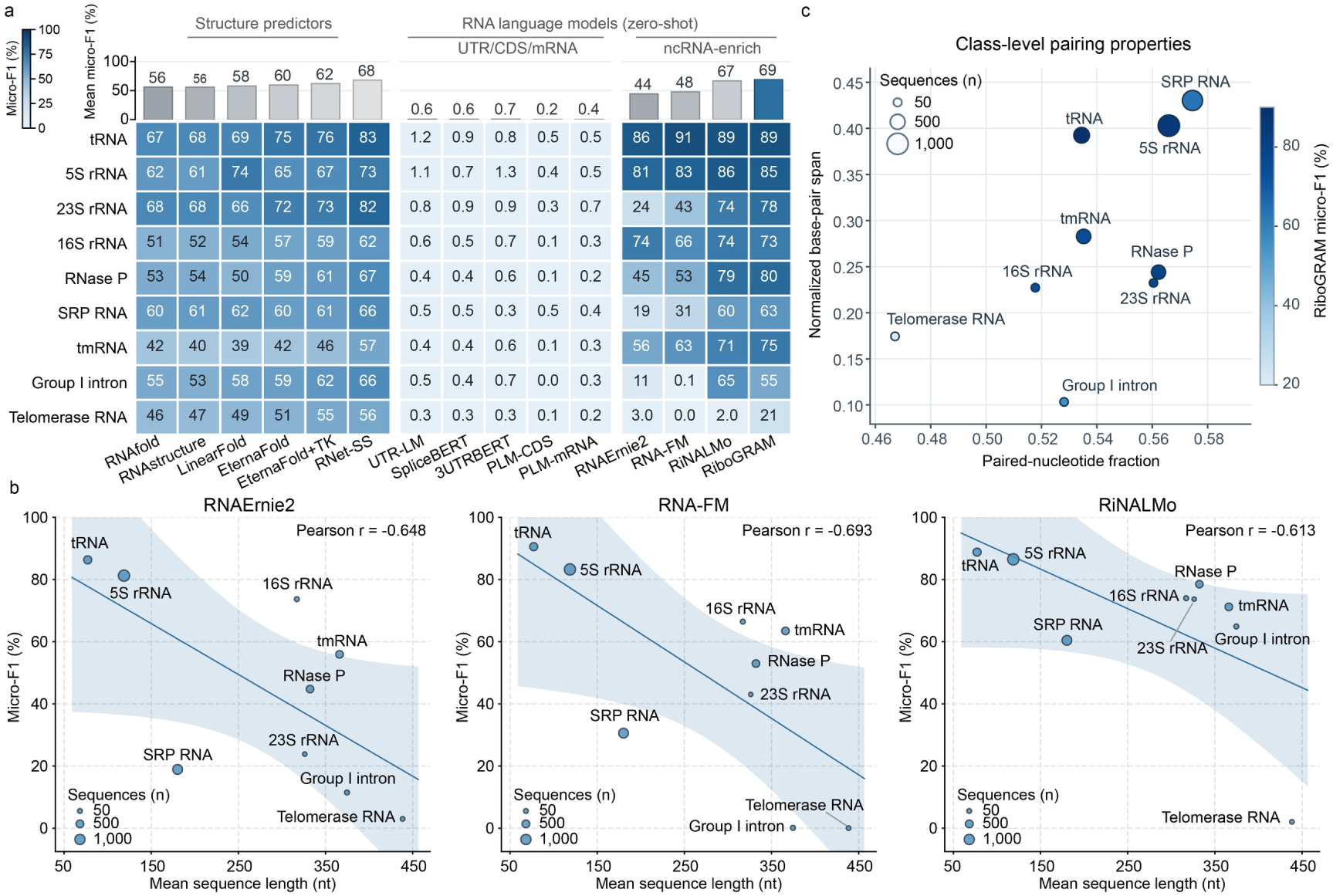
RNA-class variation in zero-shot base-pair recovery relates to length and pairing organization. a, Base-pair recovery across nine RNA classes in ArchiveII^9, 10^. Class-wise micro-F1 is shown for RNA language models and structure predictors, with the bars above indicating the unweighted mean across the nine classes for each method. EternaFold+TK denotes EternaFold with ThreshKnot decoding^4, 8^ and RNet-SS is structure-supervised^6, 7^. The benchmark comprises 3,865 sequences from 5S rRNA (n = 1,283), SRP RNA (n = 918), tRNA (n = 557), tmRNA (n = 462), RNase P RNA (n = 454), group I intron (n = 74), 16S rRNA (n = 67), telomerase RNA (n = 35) and 23S rRNA (n = 15). For RNA language models, the heads and thresholds selected on bpRNA-1m VL0 were kept fixed for ArchiveII evaluation. Corresponding full class-level micro-and macro-averaged precision, recall and F1 values are provided in Supplementary Tables 11 and 12. **b,** Base-pair recovery across RNA classes of different sequence lengths. For RNAErnie2, RNA-FM and RiNALMo, each RNA class is characterized by its micro-F1 and mean sequence length. Lines show ordinary least-squares fits, shaded bands indicate 95% confidence intervals and Pearson *r* values summarize the corresponding correlations. Bubble area indicates the number of sequences in each class. **c,** RiboGRAM base-pair recovery across RNA classes with varying pairing properties. RNA class positions are defined by paired-nucleotide fraction and mean base-pair span normalized by sequence length. Bubble color indicates RiboGRAM micro-F1 and bubble area indicates the number of sequences in each class.

**Supplementary Fig. 10.**
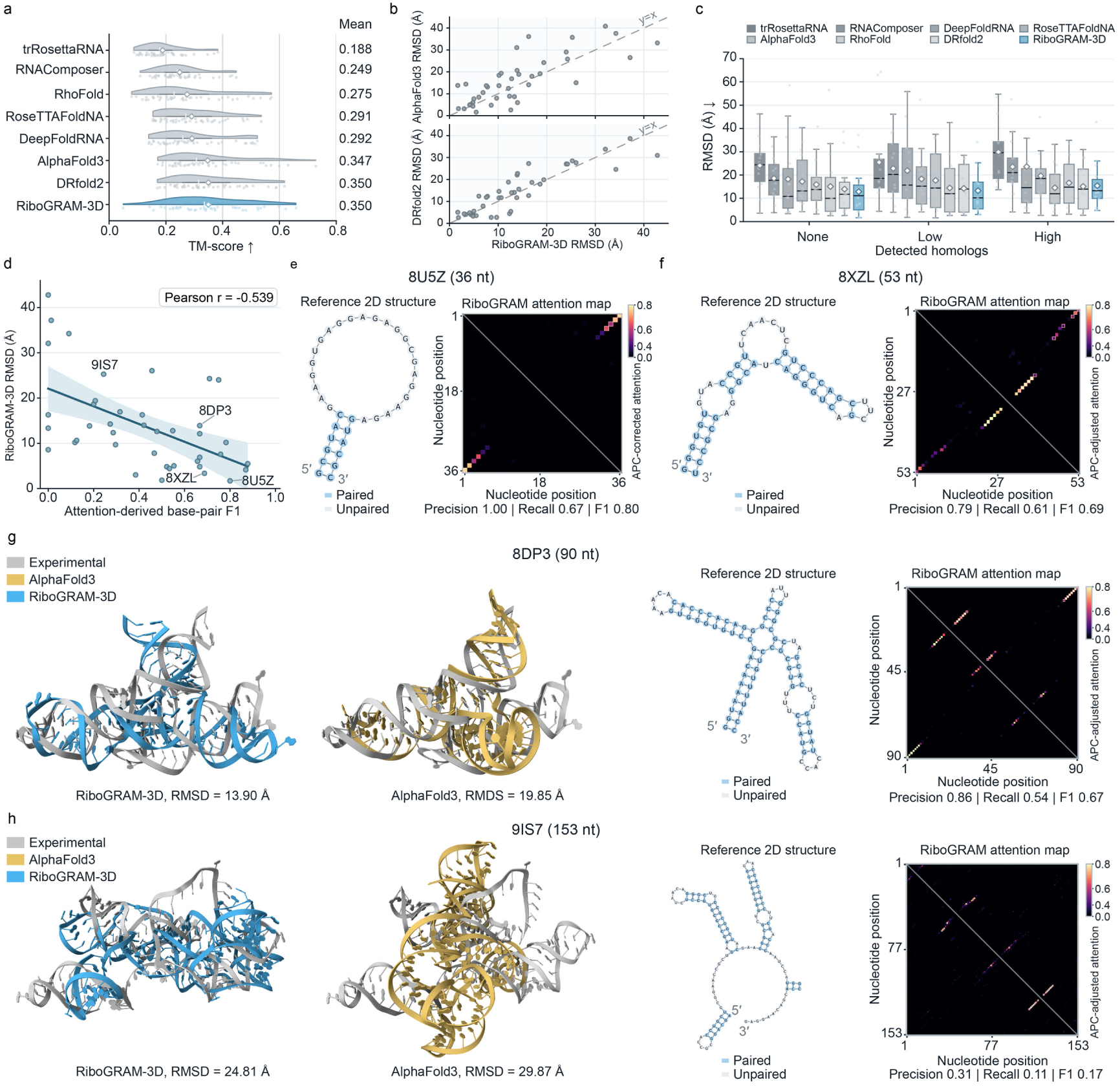
RiboGRAM-3D performance and its association with attention-derived base-pair recovery. a, TM-score distributions for eight tertiary structure prediction methods across 41 held-out RNA targets. Points denote individual targets. Horizontal bars, vertical ticks and diamonds indicate interquartile ranges, medians and means, respectively. Mean TM-scores are listed at right. **b,** Target-wise RMSD comparisons of RiboGRAM-3D with AlphaFold3 (top) and DRfold2 (bottom) across the same 41 targets in panel a. Pale blue marks the region corresponding to lower RMSD for RiboGRAM-3D. RiboGRAM-3D RMSD was lower than AlphaFold 3 for 29 of 41 targets and lower than DRfold2 for 28 of 41 targets. **c,** RMSD distributions for the same eight methods stratified by detected homologs: none (n = 14), low (n = 13) and high (n = 14). Boxes show medians and interquartile ranges, whiskers extend to 1.5 times the interquartile range, points denote individual targets and diamonds indicate means. **d,** Association between RiboGRAM attention-derived base-pair F1 and RiboGRAM-3D RMSD across the 41 held-out RNA targets. The line shows an ordinary least-squares fit, with the shaded band indicating the 95% confidence interval. **e,f**, Reference secondary structures and corresponding RiboGRAM attention maps for 8U5Z chain A (36 nt, panel **e**) and 8XZL chain A (53 nt; panel **f**). The reference secondary structures were derived from the experimentally resolved tertiary structures. For panels **e-h**, attention maps show symmetrized, APC-adjusted attention from RiboGRAM layer 12, head 20. Base pairs with attention scores above 0.4 were classified as predictions. Light-red outlines mark predicted pairs that match reference base pairs, and precision, recall and F1 are shown below each map. **g,h,** Structural superpositions, experimental secondary structures and corresponding RiboGRAM attention maps for 8DP3 chain R (90 nt; panel **g**) and 9IS7 chain A (153 nt; panel **h**). In the structural superpositions, grey denotes the experimental structure, blue for the RiboGRAM-3D prediction and ochre for the AlphaFold 3 prediction. RMSD to the experimental structure is shown for each prediction.

**Supplementary Fig. 11.**
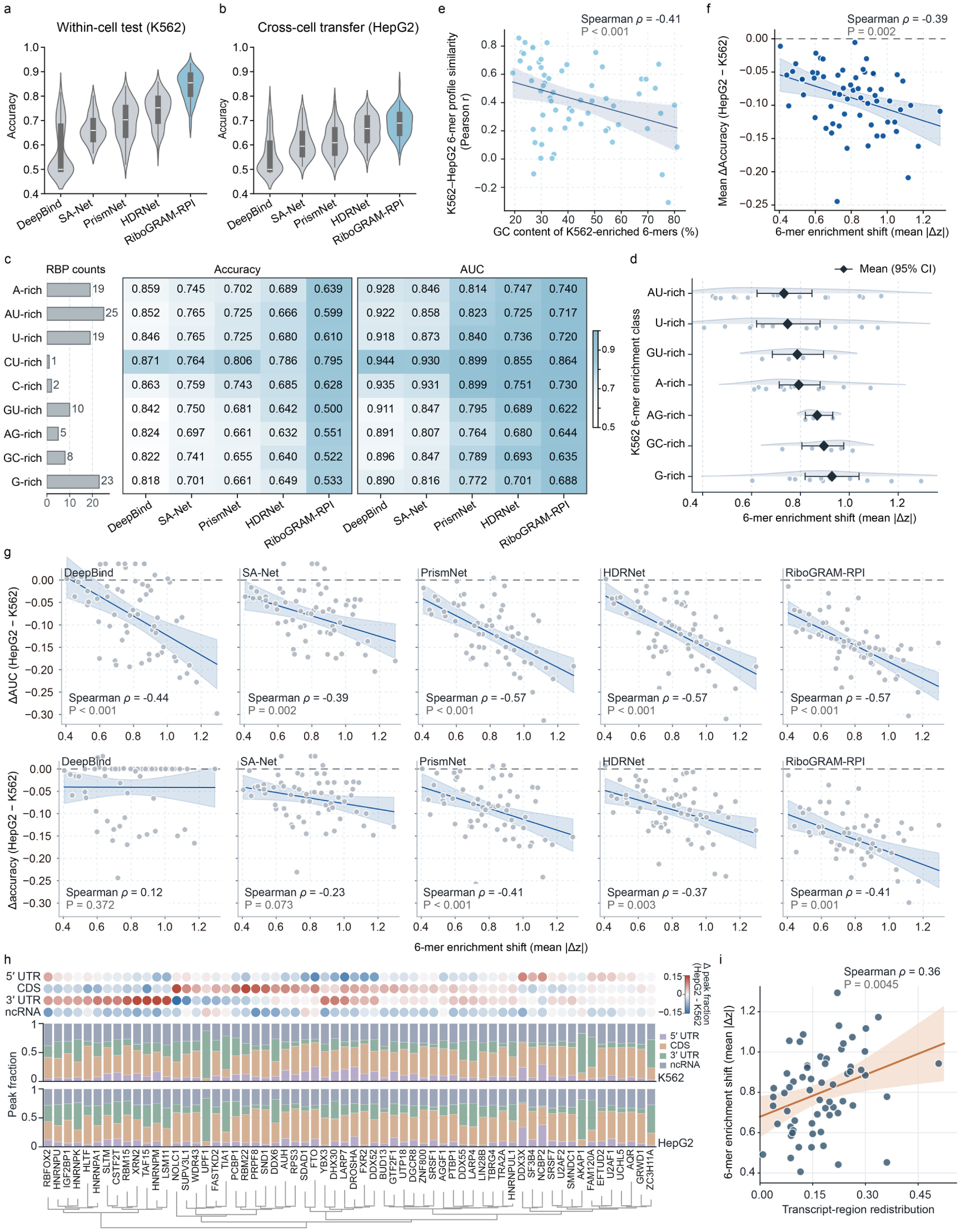
Cross-cell transfer of RBP-binding prediction is associated with 6-mer enrichment shifts and transcript-region redistribution. a,b, Accuracy distribution for DeepBind, SA-Net, PrismNet, HDRNet and RiboGRAM-RPI in the within-cell K562 test across 112 RBPs (**a**) and during cross-cell transfer to the 62 RBPs shared with HepG2 (**b**). HepG2 predictions were evaluated using the K562-trained models without cell-line-specific retraining. **c,** Predictive performance across K562-defined 6-mer enrichment classes. Mean accuracy and AUC of the five models are shown for RBPs grouped according to the nucleotide composition of their K562-enriched 6-mers (enrichment z score ≥ 1.5). Group sizes are shown at left. **d,** Cross-cell sequence-preference shifts across the same K562-defined enrichment classes. For each of the 62 RBPs shared between K562 and HepG2, the 6-mer enrichment shift was quantified as the mean absolute K562–HepG2 difference in enrichment z score across all 4,096 6-mers. Points denote individual RBPs. Diamonds and horizontal bars indicate class means and 95% confidence intervals (CI). Only classes represented by at least three shared RBPs are shown. **e,** Cross-cell 6-mer profile similarity varied with the GC content of K562-enriched 6-mers. Profile similarity was calculated as the Pearson correlation between the complete 4,096 6-mer enrichment profiles in K562 and HepG2 for each of the 62 shared RBPs. In this panel and the scatter plots in panels **f**, **g** and **i**, solid lines show fitted trends with shaded 95% CI, and Spearman ρ and associated P values are shown in each panel. **f,g,** Relationship between cross-cell 6-mer enrichment shifts and changes in predictive performance. **f,** The 6-mer enrichment shift is shown against the mean change in accuracy across the five models. **g,** The same relationship is shown separately for each model, with changes in AUC (upper row) and accuracy (lower row) for DeepBind, SA-Net, PrismNet, HDRNet and RiboGRAM-RPI. **h,i,** Transcript-region redistribution across K562 and HepG2 and its association with cross-cell 6-mer enrichment shifts. **h,** Positive eCLIP peaks for the 62 shared RBPs were distributed across the 5’ UTR, CDS, 3’ UTR and ncRNA regions. The upper matrix shows the signed change in peak fraction from K562 to HepG2, while the stacked bars show the corresponding transcript-region compositions in the two cell lines. **i,** Overall transcript-region redistribution, quantified as the total absolute change in peak fraction across the four regions, is shown against the cross-cell 6-mer shift. Each point represents one shared RBP.

**Supplementary Fig. 12.**
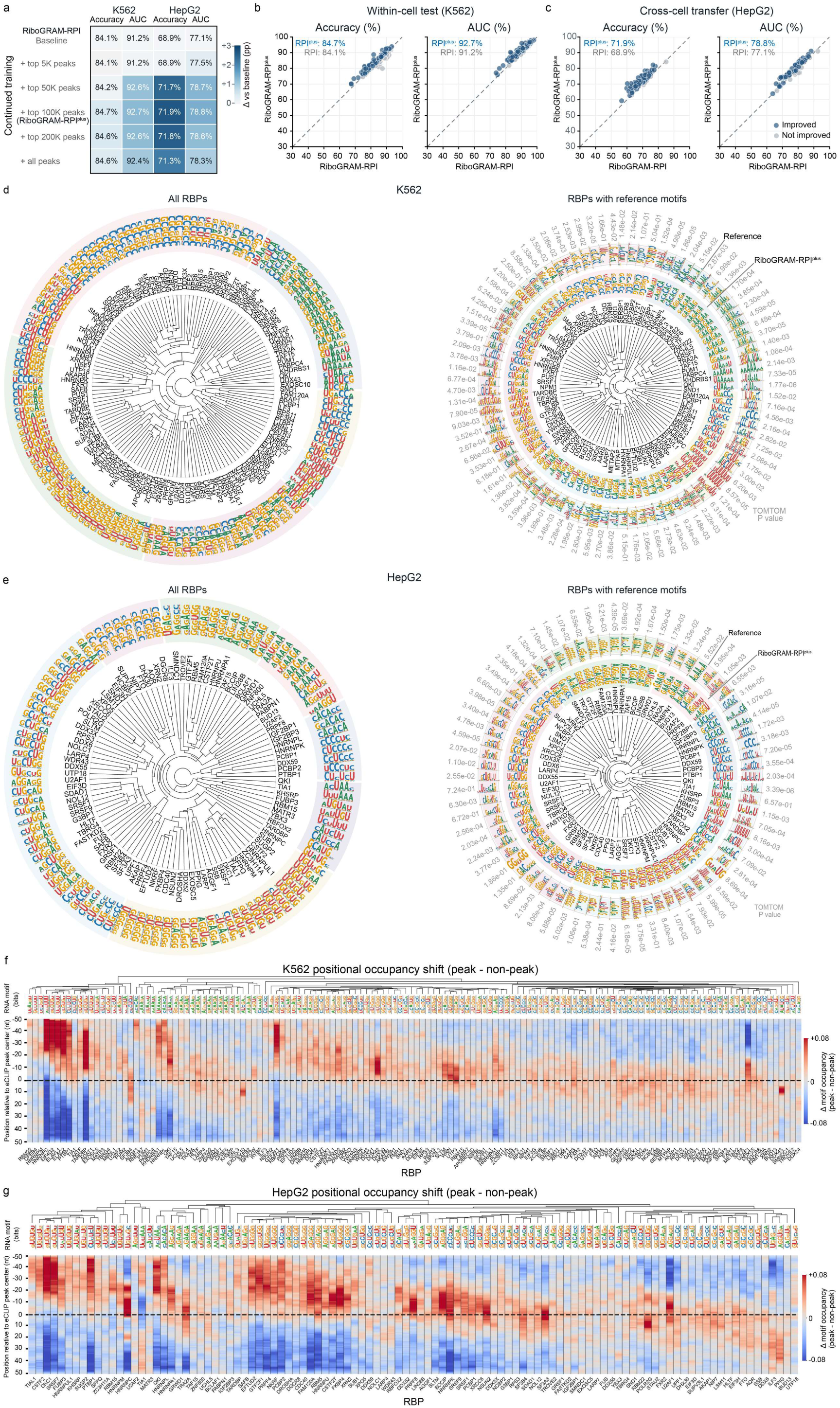
RiboGRAM-RPI^plus^ continued training and characterization of RBP-binding motifs in K562 and HepG2. a, Performance across continued training configurations of RiboGRAM-RPI. Mean accuracy and AUC across 112 K562 and 62 HepG2 RBP datasets are shown for the RiboGRAM-RPI baseline and the continued training with up to the indicated number of top-ranked peaks per RBP. Numerical values indicate absolute performance (%), while cell colors show changes relative to the corresponding baseline in percentage points (pp). The top 100K continued training configuration was designated RiboGRAM-RPIplus. **b,c,** Paired comparison of RiboGRAM-RPI and RiboGRAM-RPIplus in the within-cell K562 test (**b**) and cross-cell transfer to HepG2 (**c**). Each point represents one RBP dataset. The dashed identity line indicates equal performance between the two models, and blue and grey points denote datasets with and without improvement, respectively. Mean performance for each model is indicated in the panels. **d,e,** Organization of RBP-binding motifs and comparison with reference motifs in K562 (**d**) and HepG2 (**e**). Left, representative 6-mer motifs identified for 138 K562 and 105 HepG2 RBPs, respectively, organized according to motif similarity. Right, the corresponding subsets with available reference motifs comprise 89 K562 and 74 HepG2 RBPs. Inner and outer logos show the RiboGRAM-derived and reference motifs, respectively, and the outer labels show TOMTOM *P* values for each comparison. Reference motifs were obtained from collections of 1,071 mCrossBase motifs and 544 human CisBP-RNA motifs. TOMTOM *P* values were ≤ 0.05 for 70 of 89 K562 and 58 of 74 HepG2 motif comparisons. **f,g,** Peak-centered positional occupancy of representative RBP motifs in K562 (**f**) and HepG2 (g). For each RBP, its representative 6-mer motif was mapped across the corresponding 101-nt eCLIP peak-centered sequences and matched non-peak controls, retaining the three highest-scoring mutually non-overlapping matches per sequence. Motif occupancy at each position was calculated as the fraction of sequences covered by at least one retained match, and the heatmaps show peak-minus-control occupancy differences. Red and blue indicate greater motif occupancy in peak and matched non-peak control sequences, respectively. Columns represent RBPs, with representative motifs shown above and ordered by hierarchical clustering of their positional occupancy-difference profiles. The dashed line marks the eCLIP peak center. The analyses comprised 11,868,746 K562 and 9,864,959 HepG2 peak sequences, each with an equal number of matched non-peak controls. Sequence logos use green, blue, orange and red for A, C, G and U, respectively.

**Supplementary Fig. 13.**
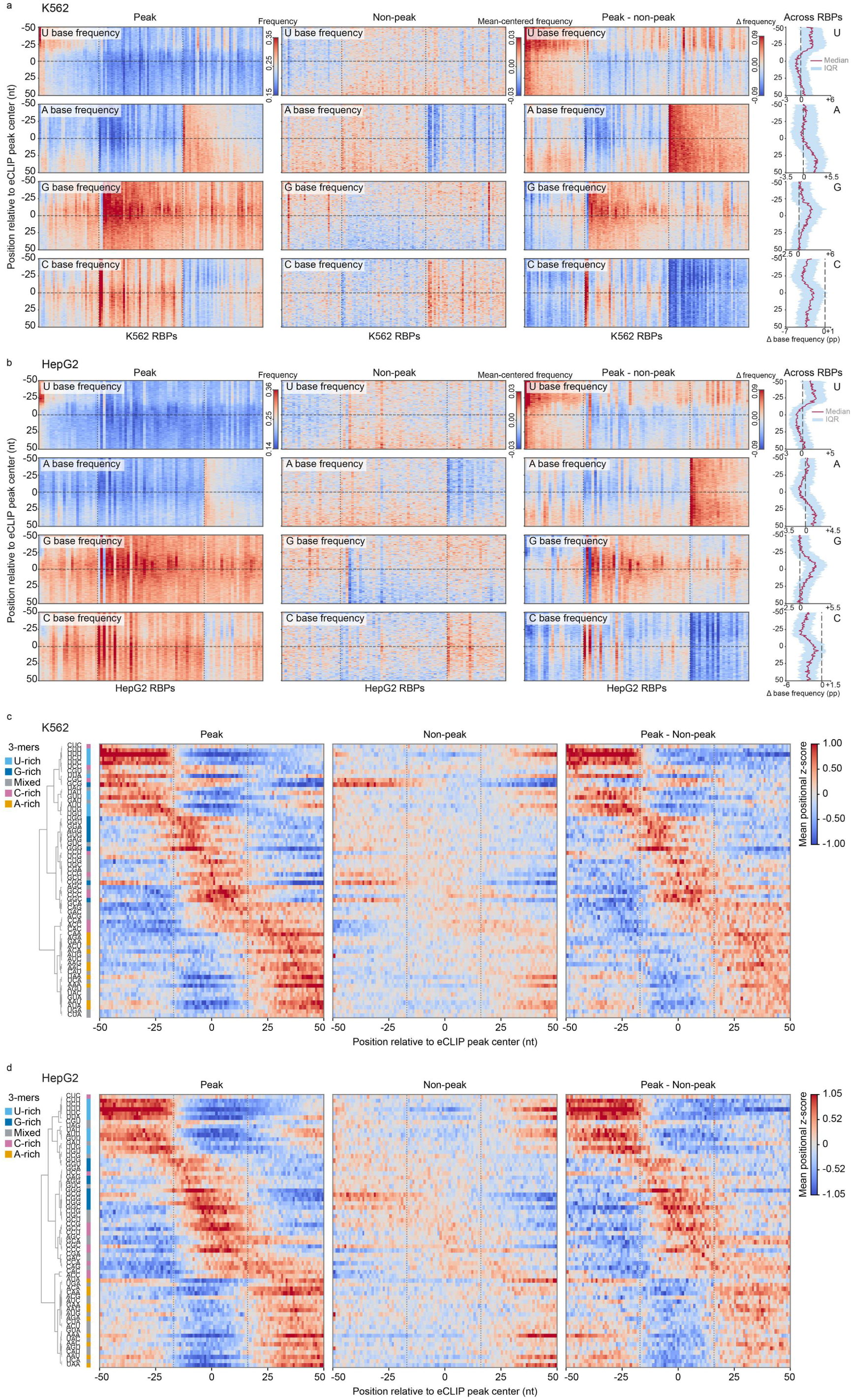
Position-resolved nucleotide composition and 3-mer organization around eCLIP peak centers in K562 and HepG2. a,b, Position-resolved base-frequency profiles in K562 (**a**) and HepG2 (**b**). The analysis used the complete HDRNet datasets^11^ for K562 (112 RBPs) and HepG2 (84 RBPs). For U, A, G and C, heatmaps show base frequencies in 101-nt peak-centered sequences, mean-centered frequencies in matched non-peak sequences and the corresponding peak-minus-non-peak differences. Right, median peak-minus-non-peak profiles across RBPs are shown with interquartile ranges, with frequency differences denoted in percentage points (pp). **c,d,** Position-resolved 3-mer profiles in K562 (**c**) and HepG2 (**d**). Heatmaps show mean positional z scores for all 64 3-mers across RBPs in peak and matched non-peak sequences, together with the corresponding peak-minus-non-peak differences. Rows are ordered by hierarchical clustering of the mean positional z-score profiles of the 64 3-mers in peak sequences, with adjacent color strips denoting U-rich, G-rich, C-rich, A-rich and mixed composition classes.

**Supplementary Fig. 14.**
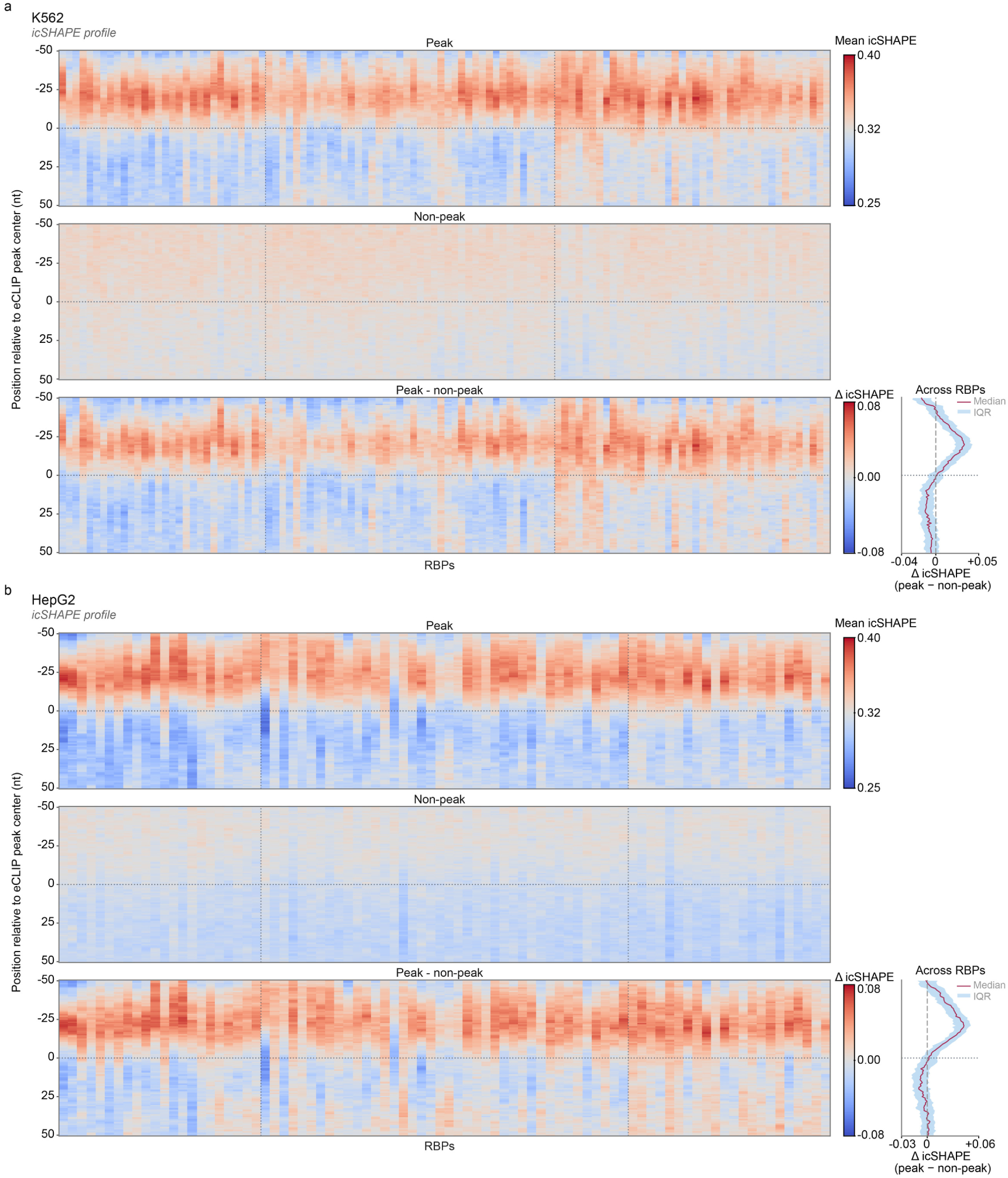
Position-resolved RNA accessibility profiles around eCLIP peak centers in K562 and HepG2.a,b, Position-resolved icSHAPE reactivity profiles in K562 (**a**) and HepG2 (**b**). The analyses include all available HDRNet RBP datasets for K562 (112 RBPs) and HepG2 (84 RBPs). For each RBP, heatmaps show mean icSHAPE reactivity across 101-nt peak-centered and corresponding non-peak sequences, together with the peak-minus-non-peak difference. Columns represent individual RBPs. Right, median peak-minus-non-peak profiles across RBPs are shown with interquartile ranges. Higher icSHAPE values indicate greater local nucleotide accessibility. The horizontal dotted line marks the eCLIP peak center.

---

**Supplementary Table 1.**
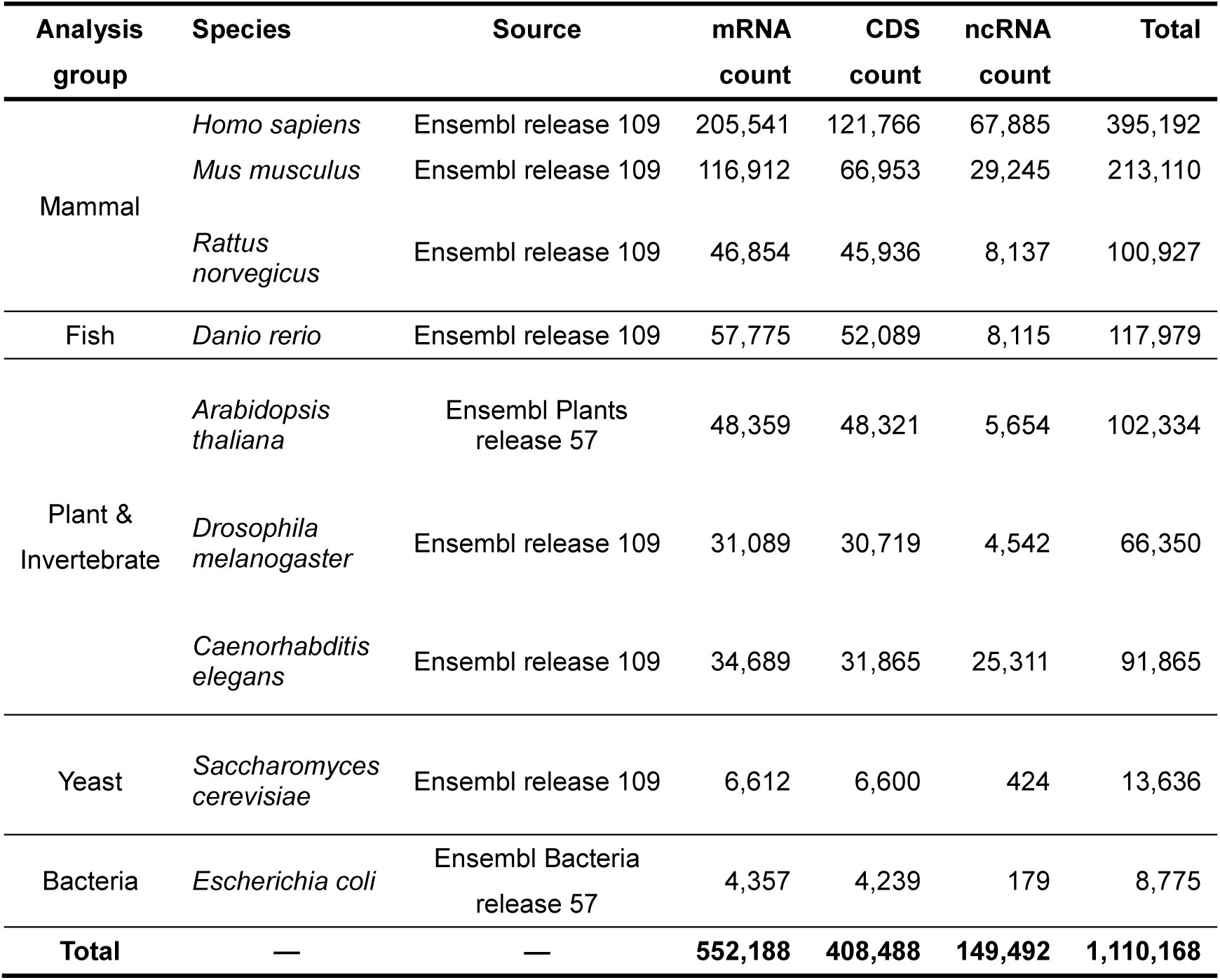
mRNA, CDS and ncRNA sequence collection across nine model organisms.

**Supplementary Table 2.** Cross-species ncRNA–CDS and ncRNA–mRNA binary classification datasets.

| Analysis group | Species | Split | ncRNA–mRNA Binary Classification |  |  | ncRNA–CDS Binary Classification |  |  |
| --- | --- | --- | --- | --- | --- | --- | --- | --- |
|  |  |  | ncRNA | mRNA | Total | ncRNA | CDS | Total |
| Mammal | *Homo sapiens* | Training | 5,000 | 5,000 | 10,000 | 5,000 | 5,000 | 10,000 |
|  |  | Validation | 1,000 | 1,000 | 2,000 | 1,000 | 1,000 | 2,000 |
|  |  | Test | 5,000 | 5,000 | 10,000 | 5,000 | 5,000 | 10,000 |
|  | *Mus musculus* | Test | 5,000 | 5,000 | 10,000 | 5,000 | 5,000 | 10,000 |
|  | *Rattus norvegicus* | Test | 5,000 | 5,000 | 10,000 | 5,000 | 5,000 | 10,000 |
| Fish | *Danio rerio* | Test | 5,000 | 5,000 | 10,000 | 5,000 | 5,000 | 10,000 |
| Plant & Invertebrate | *Arabidopsis thaliana* | Test | 5,000 | 5,000 | 10,000 | 5,000 | 5,000 | 10,000 |
|  | *Drosophila melanogaster* | Test | 4,478 | 5,000 | 9,478 | 4,265 | 5,000 | 9,265 |
|  | *Caenorhabditis elegans* | Test | 5,000 | 5,000 | 10,000 | 5,000 | 5,000 | 10,000 |
| Yeast | *Saccharomyces cerevisiae* | Test | 420 | 5,000 | 5,420 | 416 | 5,000 | 5,416 |
| Bacteria | *Escherichia coli* | Test | 179 | 4,357 | 4,536 | 162 | 3,871 | 4,033 |
| **Total** | **—** | **—** | **41,077** | **50,357** | **91,434** | **40,843** | **49,871** | **90,714** |

**Supplementary Table 3.**
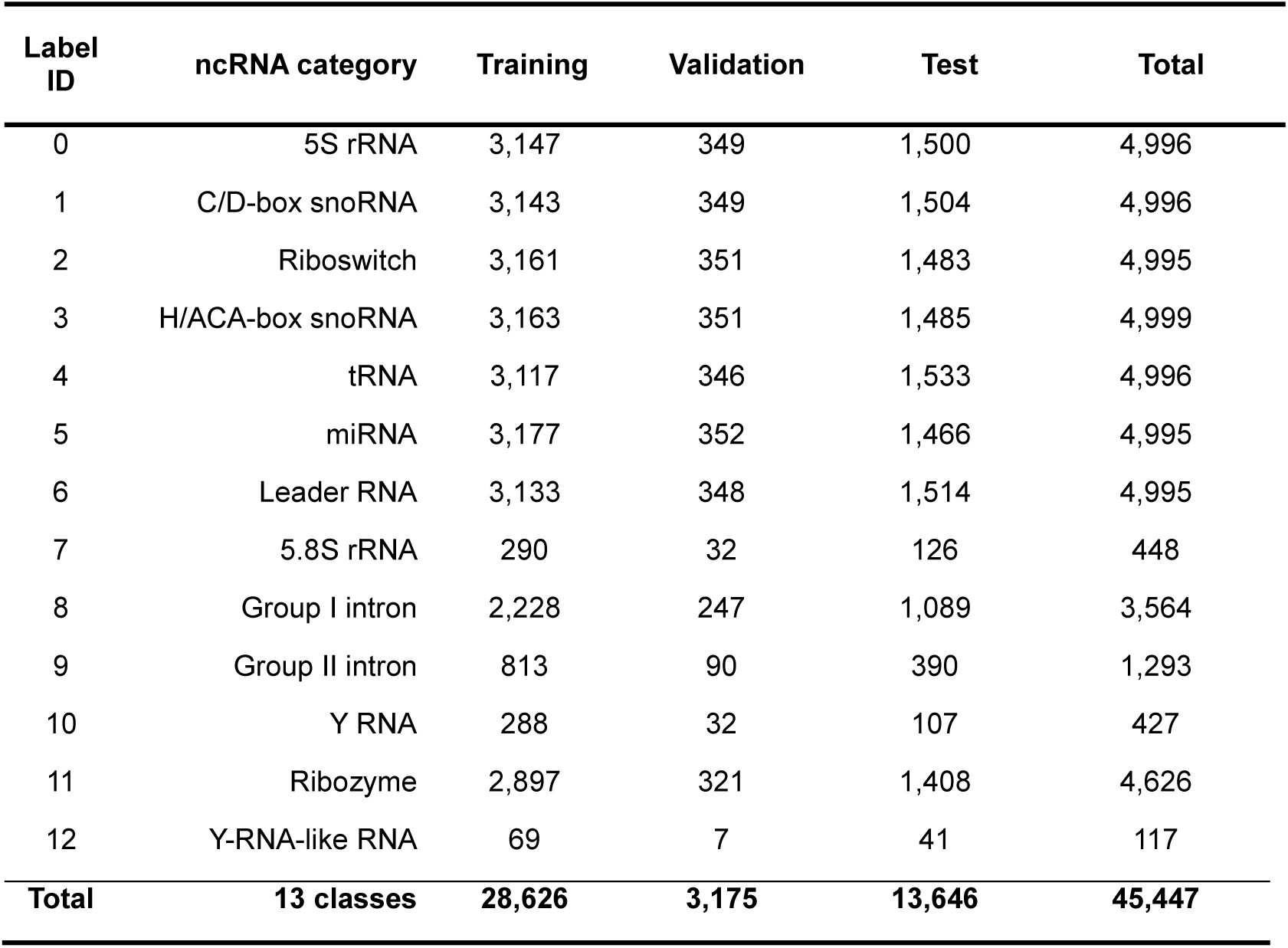
NCYPred 13-class ncRNA classification dataset.

| **Label ID** | **ncRNA category** | **Training** | **Validation** | **Test** | **Total** |
| --- | --- | --- | --- | --- | --- |
| 0 | 5S rRNA | 3,147 | 349 | 1,500 | 4,996 |
| 1 | C/D-box snoRNA | 3,143 | 349 | 1,504 | 4,996 |
| 2 | Riboswitch | 3,161 | 351 | 1,483 | 4,995 |
| 3 | H/ACA-box snoRNA | 3,163 | 351 | 1,485 | 4,999 |
| 4 | tRNA | 3,117 | 346 | 1,533 | 4,996 |
| 5 | miRNA | 3,177 | 352 | 1,466 | 4,995 |
| 6 | Leader RNA | 3,133 | 348 | 1,514 | 4,995 |
| 7 | 5.8S rRNA | 290 | 32 | 126 | 448 |
| 8 | Group I intron | 2,228 | 247 | 1,089 | 3,564 |
| 9 | Group II intron | 813 | 90 | 390 | 1,293 |
| 10 | Y RNA | 288 | 32 | 107 | 427 |
| 11 | Ribozyme | 2,897 | 321 | 1,408 | 4,626 |
| 12 | Y-RNA-like RNA | 69 | 7 | 41 | 117 |
| **Total** | **13 classes** | **28,626** | **3,175** | **13,646** | **45,447** |

**Supplementary Table 4.** Rfam 88-family ncRNA classification dataset.

| ncRNA macro-class | Rfam families | Training | Validation | Test | Total |
| --- | --- | --- | --- | --- | --- |
| snRNA/snoRNA | 22 | 26,466 | 3,732 | 5,434 | 35,632 |
| Cis-regulatory RNA | 32 | 38,496 | 7,318 | 9,358 | 55,172 |
| miRNA | 17 | 20,451 | 1,669 | 1,657 | 23,777 |
| sRNA | 7 | 8,421 | 1,304 | 1,298 | 11,023 |
| Intron | 6 | 7,218 | 488 | 486 | 8,192 |
| rRNA | 2 | 2,406 | 1,387 | 4,362 | 8,155 |
| tRNA | 2 | 2,406 | 1,426 | 2,747 | 6,579 |
| **Total** | **88** | **105,864** | **17,324** | **25,342** | **148,530** |

**Note.** The Rfam-novel benchmark was derived from Rfam 14.1 and contains 88 RNA families, which are presented as seven ncRNA macro-classes following the original dataset definition.

**Supplementary Table 5.** Summary of Rfam v15.0 seed sequences used for family-level contextual-dependency analysis.

| **Rfam class** | **Number of RNA subtypes** | **Number of Rfam families** | **Sequences analyzed** |
| --- | --- | --- | --- |
| Gene | 16 | 3,644 | 50,352 |
| Cis-regulator | 6 | 525 | 21,861 |
| Intron | 1 | 9 | 1,246 |
| **Total** | **23** | **4,178** | **73,459** |

**Supplementary Table 6.** Pretraining corpora for the architecture-matched PLM-CDS and PLM-mRNA controls.

| **Pretraining corpus** | **Corresponding model** | **Source** | **Sequence type** | **Number of sequences (million)** | **Total nucleotides (billion nt)** |
| --- | --- | --- | --- | --- | --- |
| CDS corpus | PLM-CDS | Ensembl release 112 | CDS | 13.43 | 24.41 |
| mRNA corpus | PLM-mRNA | Ensembl release 112 | mRNA (spliced transcript) | 14.30 | 35.22 |

**Supplementary Table 7.**
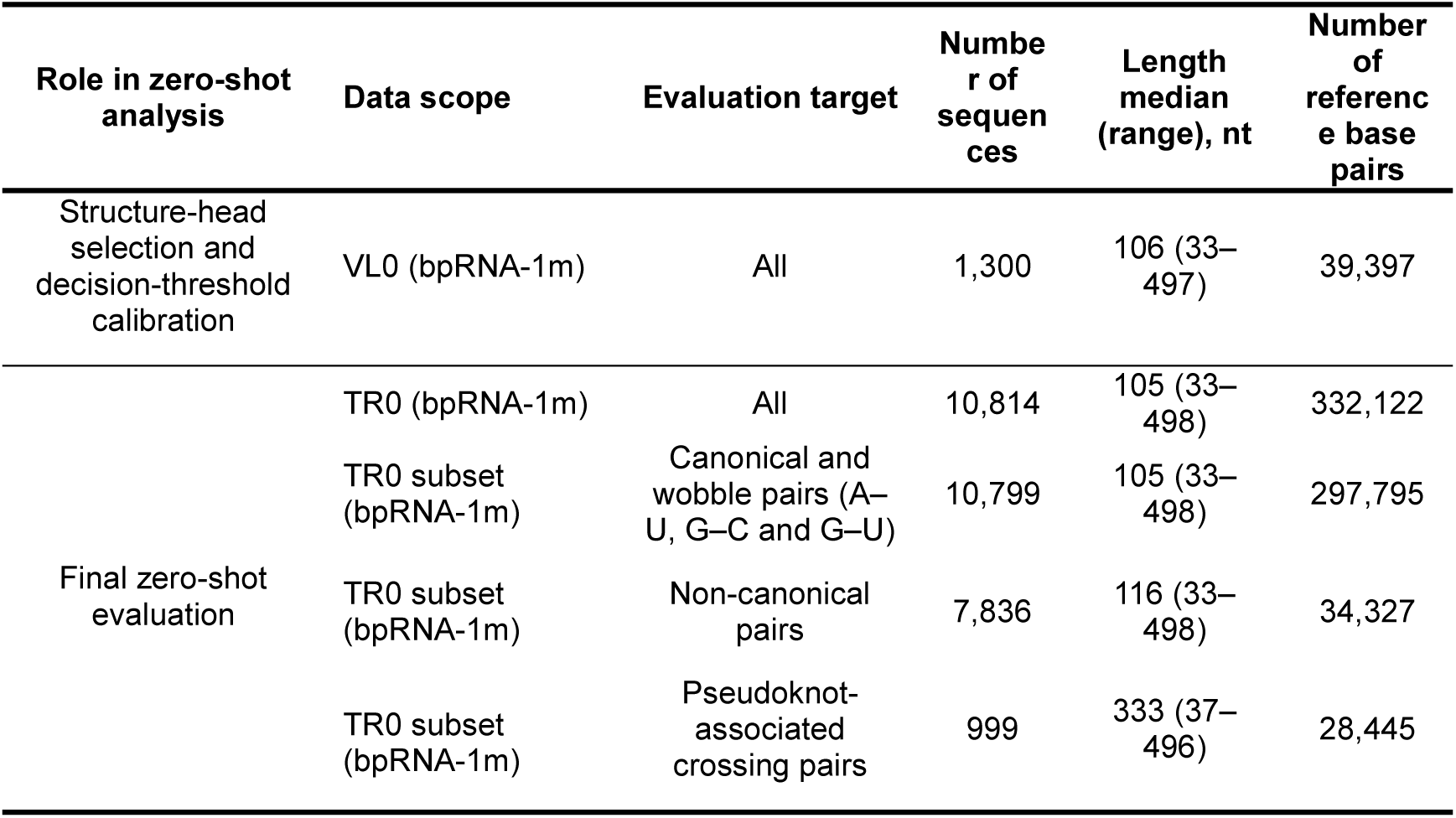
bpRNA-1m dataset for zero-shot ncRNA base-pair analysis.

**Supplementary Table 8.** Micro-and macro-averaged recovery of all annotated base pairs on the bpRNA-1m TR0 subset.

| Method group | Model | All annotated base pairs |  |  |  |  |  |
| --- | --- | --- | --- | --- | --- | --- | --- |
|  |  | Micro-averaged |  |  | Macro-averaged |  |  |
|  |  | P (%) | R (%) | F1 (%) | P (%) | R (%) | F1 (%) |
| Structure predictors | RNAfold | 42.8 | 56.8 | 48.8 | 45.4 | 58.5 | 49.7 |
|  | RNAstructure | 42.5 | 55.9 | 48.3 | 45.1 | 57.6 | 49.2 |
|  | LinearFold | 50.5 | 51.9 | 51.2 | 51.6 | 51.4 | 50.4 |
|  | EternaFold | 45.8 | 59.0 | 51.6 | 47.6 | 60.7 | 51.9 |
|  | EternaFold+TK | 53.6 | 55.2 | 54.4 | 53.6 | 57.0 | 53.4 |
|  | RNet-SS | 50.8 | 61.2 | 55.6 | 51.0 | 62.7 | 54.6 |
| UTR/CDS/mRNA | UTR-LM | 0.4 | 83.6 | 0.7 | 0.7 | 84.3 | 1.3 |
|  | 3UTRBERT | 0.5 | 42.9 | 0.9 | 0.6 | 40.8 | 1.1 |
|  | PLM-CDS | 0.2 | 43.1 | 0.4 | 0.4 | 46.2 | 0.8 |
|  | SpliceBERT | 0.4 | 69.8 | 0.8 | 0.7 | 68.9 | 1.4 |
|  | PLM-mRNA | 0.3 | 48.3 | 0.5 | 0.5 | 49.5 | 0.9 |
| ncRNA-rich | RNAErnie2 | 36.9 | 27.8 | 31.7 | 30.7 | 32.7 | 28.9 |
|  | RNA-FM | 56.1 | 32.6 | 41.2 | 43.4 | 38.1 | 36.6 |
|  | RiNALMo | 57.1 | 46.2 | 51.1 | 47.9 | 47.7 | 45.5 |
|  | **RiboGRAM** | **62.3** | **54.5** | **58.2** | **54.6** | **56.5** | **52.1** |

**Note.** Values are percentages. P, R and F1 denote pair-level precision, recall and F1, respectively. Micro metrics were calculated by pooling base-pair-level TP, FP and FN across all 10,814 TR0 sequences. Macro metrics were calculated for each sequence and then averaged across sequences.

**Supplementary Table 9.** Zero-shot pair-level recovery of canonical and wobble, non-canonical, and pseudoknot-associated base pairs.

| Method group | Model | Canonical/wobble (A–U, G–C, G–U) |  |  | Non-canonical |  |  | Pseudoknot-associated crossing pairs |  |  |
| --- | --- | --- | --- | --- | --- | --- | --- | --- | --- | --- |
|  |  | P (%) | R (%) | F1 (%) | P (%) | R (%) | F1 (%) | P (%) | R (%) | F1 (%) |
| Structure predictors | RNAfold | 42.8 | 63.3 | 51.1 | 0.0 | 0.0 | 0.0 | 0.0 | 0.0 | 0.0 |
|  | RNAstructure | 42.5 | 62.3 | 50.5 | 0.0 | 0.0 | 0.0 | 0.0 | 0.0 | 0.0 |
|  | LinearFold | 50.5 | 57.9 | 54.0 | 0.0 | 0.0 | 0.0 | 0.0 | 0.0 | 0.0 |
|  | EternaFold | 45.8 | 65.8 | 54.0 | 0.0 | 0.0 | 0.0 | 0.0 | 0.0 | 0.0 |
|  | EternaFold+TK | 53.6 | 61.6 | 57.3 | 0.0 | 0.0 | 0.0 | 25.3 | 2.3 | 4.2 |
|  | RNet-SS | 50.9 | 68.3 | 58.3 | 25.9 | 0.1 | 0.1 | 35.0 | 37.5 | 36.2 |
| UTR/CD S/mRNA | UTR-LM | 1.1 | 1.6 | 1.3 | 0.1 | 0.8 | 0.1 | 0.0 | 0.9 | 0.1 |
|  | 3UTRBERT | 1.2 | 1.1 | 1.2 | 0.1 | 1.3 | 0.2 | 0.0 | 0.8 | 0.1 |
|  | PLM-CDS | 0.7 | 0.7 | 0.7 | 0.1 | 0.9 | 0.1 | 0.0 | 0.4 | 0.0 |
|  | SpliceBERT | 1.5 | 1.3 | 1.4 | 0.1 | 1.4 | 0.2 | 0.0 | 0.8 | 0.1 |
|  | PLM-mRNA | 0.8 | 0.8 | 0.8 | 0.1 | 1.0 | 0.1 | 0.0 | 0.4 | 0.0 |
| ncRNA-rich | RNAErnie2 | 55.9 | 28.2 | 37.5 | 4.5 | 10.2 | 6.3 | 4.4 | 20.9 | 7.3 |
|  | RNA-FM | 69.7 | 33.9 | 45.6 | 11.4 | 12.3 | 11.9 | 13.2 | 25.0 | 17.3 |
|  | RiNALMo | 67.4 | 49.2 | 56.9 | 12.2 | 16.8 | 14.1 | 17.6 | 47.2 | 25.6 |
|  | **RiboGRAM** | **72.4** | **57.7** | **64.2** | **16.5** | **24.1** | **19.6** | **31.1** | **54.1** | **39.5** |

**Note.** Values are percentages. P, R, and F1 denote pair-level micro-precision, micro-recall, and micro-F1, respectively. For each pair category, metrics were calculated by pooling base-pair-level TP, FP, and FN across the bpRNA-1m TR0 test set. Each base pair was counted once ( $i < j$ ). Canonical/wobble pairs comprise A–U, G–C, and G–U; all other pairs were classified as non-canonical. Pseudoknot-associated pairs were defined as base pairs involved in at least one crossing interaction ( $i < k < j < l$ ) and were identified separately in the reference and predicted structures. Metrics with a zero denominator were set to 0.

**Supplementary Table 10.** ArchiveII dataset for zero-shot ncRNA base-pair analysis across nine RNA classes.

| RNA class | Number of sequences | Sequence length, median (range), nt | Number of reference base pairs | Number of pseudoknot-associated reference base pairs |
| --- | --- | --- | --- | --- |
| 5S rRNA | 1,283 | 119 (102–135) | 43,124 | 0 |
| SRP RNA | 918 | 117 (28–509) | 49,838 | 0 |
| tRNA | 557 | 76 (54–93) | 11,445 | 0 |
| tmRNA | 462 | 363 (102–437) | 45,332 | 26,153 |
| RNase P RNA | 454 | 330 (120–486) | 42,717 | 10,133 |
| Group I intron | 74 | 395 (210–510) | 7,321 | 907 |
| 16S rRNA | 67 | 353 (73–512) | 5,582 | 198 |
| Telomerase RNA | 35 | 444 (382–508) | 3,578 | 969 |
| 23S rRNA | 15 | 347 (242–380) | 1,370 | 17 |
| **Total** | **3,865** | **120 (28–512)** | **210,307** | **38,377** |

**Note.** Pseudoknot-associated reference base pairs were defined as unique annotated base pairs participating in at least one crossing interaction, $i < k < j < l$ . Each base pair was counted only once. Zero indicates that no pseudoknot-associated reference base pairs were identified.

**Supplementary Table 11.**
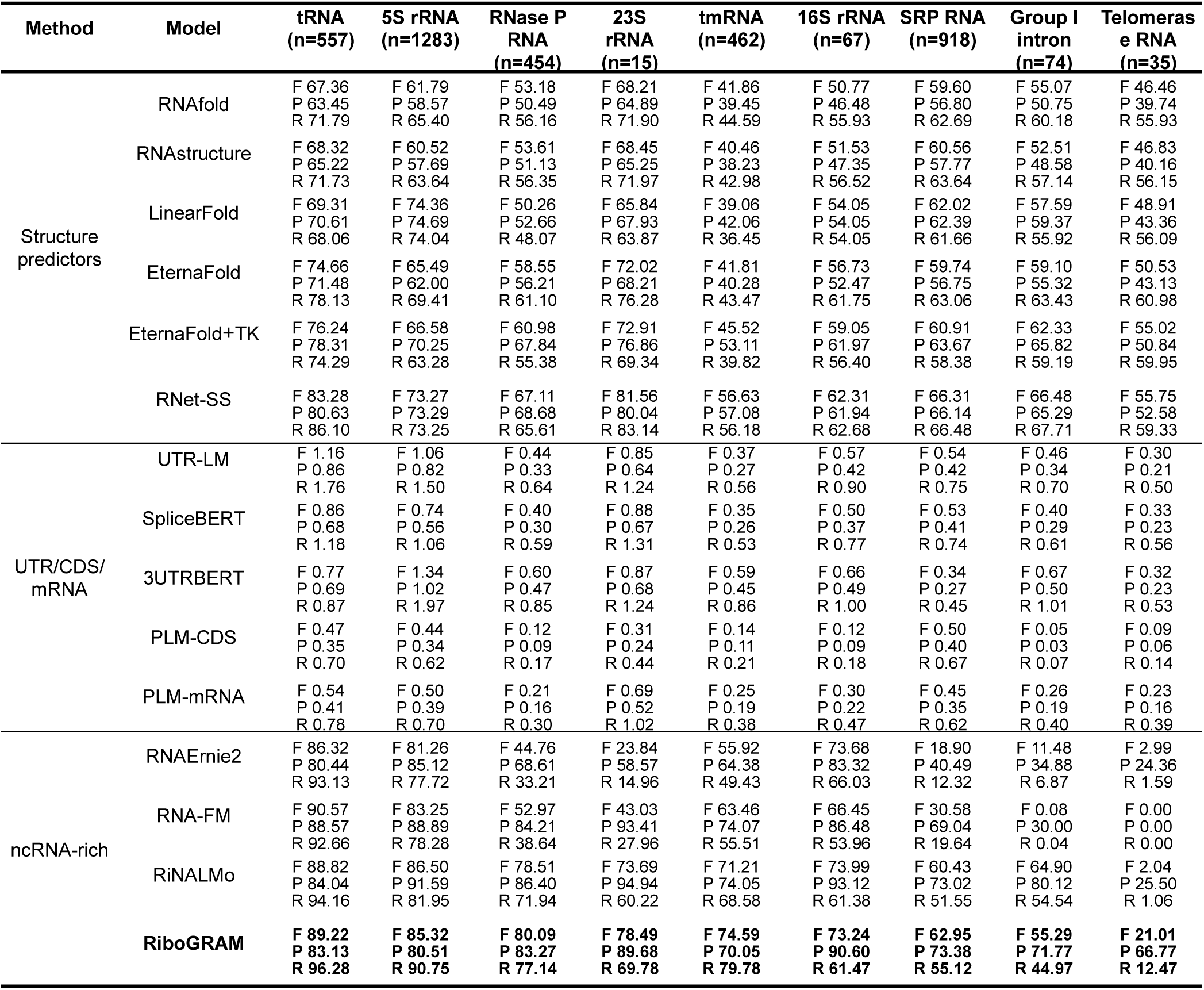
Micro-averaged base-pair recovery performance across nine ArchiveII RNA classes.

**Supplementary Table 12.**
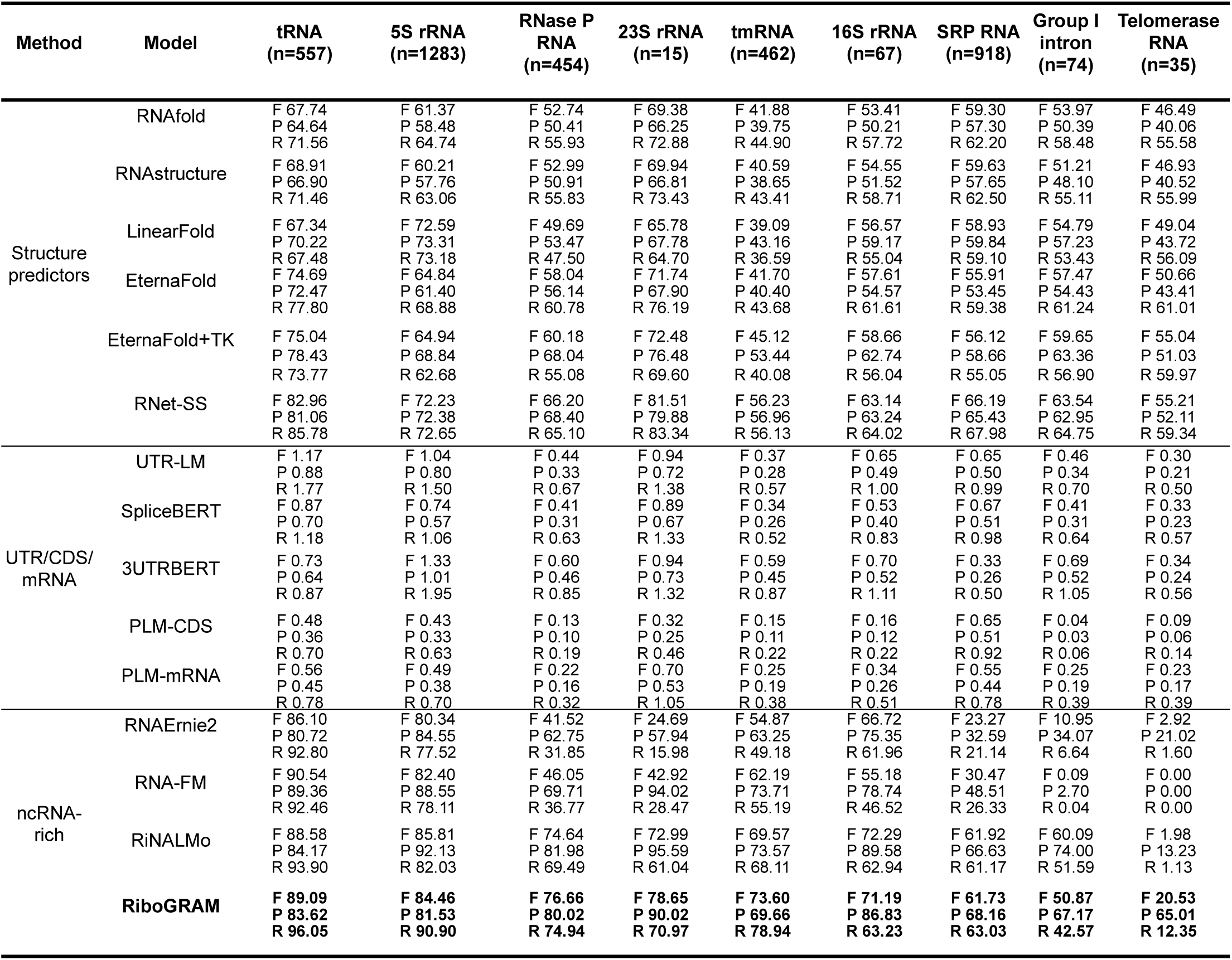
Macro-averaged base-pair recovery performance across nine ArchiveII RNA classes.

**Supplementary Table 13.** Datasets for intra-family and inter-family nucleotide pairing-state probing.

| Probing setting | Dataset | Split | Number of sequences | Sequence length, median (range), nt | Paired nucleotides | Unpaired nucleotides |
| --- | --- | --- | --- | --- | --- | --- |
| Intra-family probing on bpRNA-1m | TR0 | Training | 10,814 | 105 (33–498) | 664,244 | 782,415 |
|  | VL0 | Validation | 1,300 | 106 (33–497) | 78,794 | 92,635 |
|  | TS0 | Test | 1,305 | 109 (22–499) | 81,570 | 95,977 |
| Inter-family probing (TrainSetA-to-TestSetB) | TrainSetA subset | Training | 2,850 | 115 (10–734) | 302,538 | 266,690 |
|  | TrainSetA subset | Validation | 316 | 107.5 (10–695) | 32,102 | 28,949 |
|  | TestSetB | Test | 430 | 109 (27–244) | 22,858 | 29,239 |

**Supplementary Table 14.**
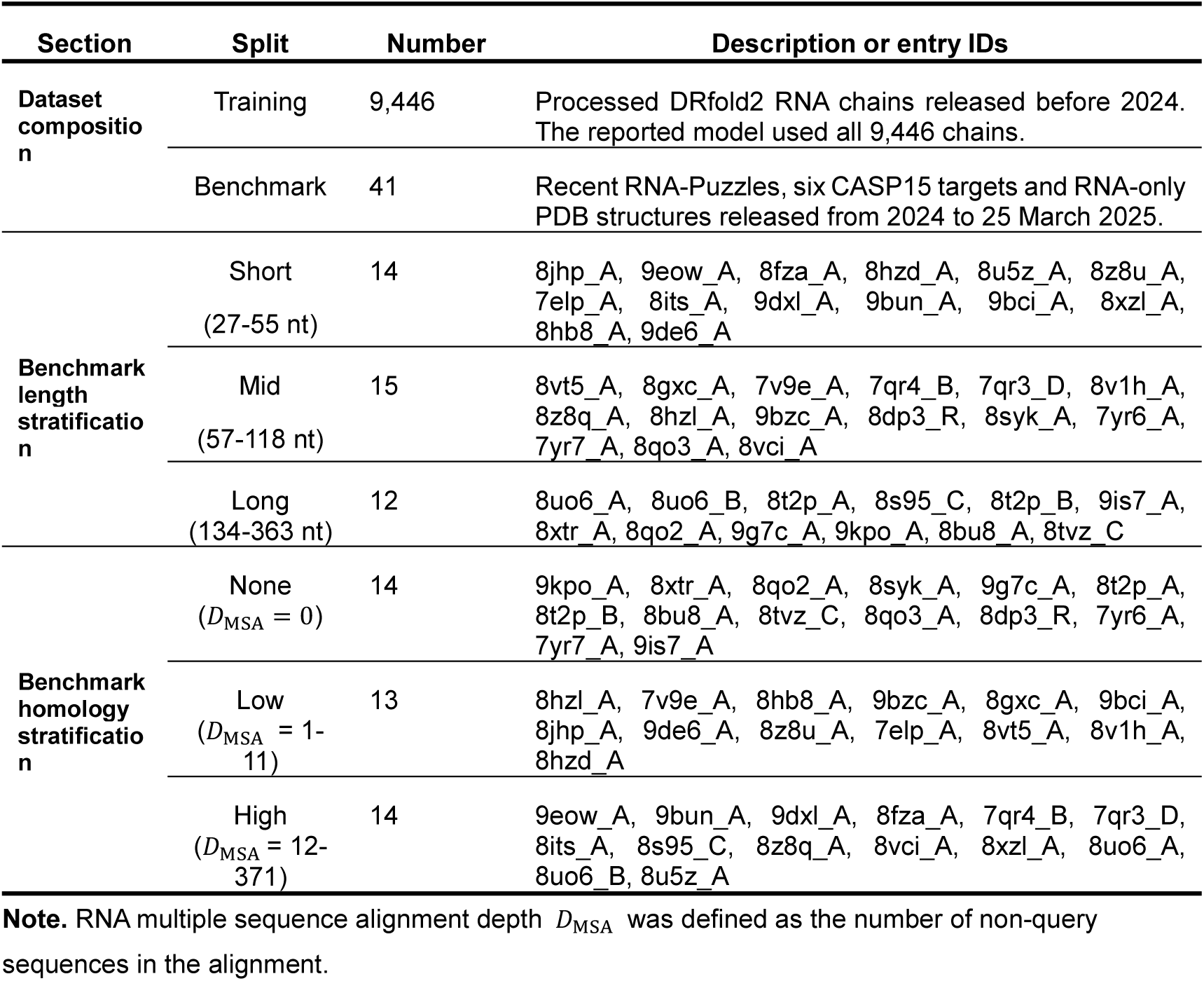
RiboGRAM-3D training data and stratification of the 41-target benchmark by sequence length and homolog depth.

**Supplementary Table 15.**
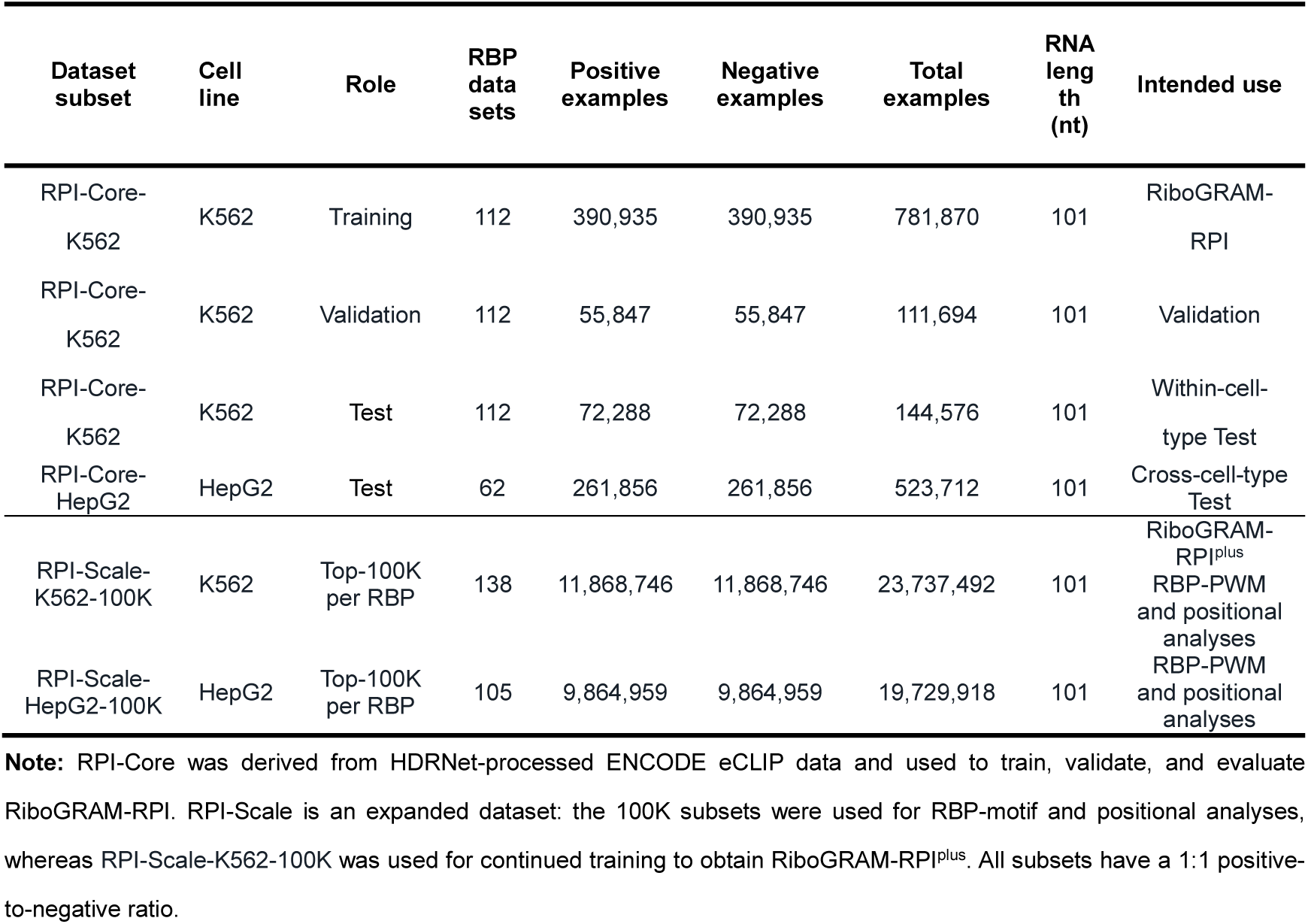
RNA–protein interaction datasets for RiboGRAM-RPI training, evaluation and downstream analysis.

**Supplementary Table 16.** RNA–RNA interaction datasets for RiboGRAM-RRI training and evaluation.

| **Dataset** | **Data split** | **Positive**<br>**pairs** | **Negative**<br>**pairs** | **Total**<br>**pairs** |
| --- | --- | --- | --- | --- |
| MirTarRAW | Training | 1,386 | 1,386 | 2,772 |
|  | Validation | 1,386 | 1,386 | 2,772 |
|  | Test | 11,088 | 11,088 | 22,176 |
|  | **Total** | **13,860** | **13,860** | **27,720** |
| RISE-derived | Training | 59,834 | 202,163 | 261,997 |
|  | Validation | 7,858 | 26,326 | 34,184 |
|  | Test | 6,622 | 21,929 | 28,551 |
|  | **Total** | **74,314** | **250,418** | **324,732** |

**Supplementary Table 17.** RNA–chromatin interaction datasets for RiboGRAM-RCI training and cross-context evaluation.

| Cell type | Assay | GEO accession(s) | Data split | Positive pairs | Negative pairs |
| --- | --- | --- | --- | --- | --- |
| HEK293T | iMARGI | GSE122690;<br>GSM3478205 | Training | 4,800,000 | 4,800,000 |
|  |  |  | Validation | 100,000 | 100,000 |
|  |  |  | Test<br>(same cell type,<br>same assay) | 100,000 | 100,000 |
| HUVEC | iMARGI | GSE135357;<br>GSM4006840;<br>GSM4729222 | Test<br>(cross-cell-type,<br>same assay) | 100,000 | 100,000 |
| Fibroblasts | Red-C | GSE136141 | Test<br>(cross-cell-type,<br>cross-assay) | 100,000 | 100,000 |
| K562 | Red-C | GSE136141 | Test<br>(cross-cell-type,<br>cross-assay) | 100,000 | 100,000 |
| MDA-MB-231 | GRID-seq | GSE82312 | Test<br>(cross-cell-type,<br>cross-assay) | 100,000 | 100,000 |
| MM.1S | GRID-seq | GSE82312 | Test<br>(cross-cell-type,<br>cross-assay) | 100,000 | 100,000 |

**Supplementary Note 1. RiboGRAM representations of lineage, structural constraints and functional variation in 5S rRNA.** 5S rRNA retains a highly conserved structural scaffold across Bacteria, Archaea and Eukaryota with sequence variation across taxonomic lineages and cellular compartments. We therefore used 5S rRNA as a within-family system to examine how RiboGRAM organizes this sequence variation within a shared structural framework and whether it relates to functional variation. Among 677 curated records from the Rfam 5S rRNA seed alignment (RF00001)^12^, RiboGRAM representations separated canonical eukaryotic cytoplasmic 5S rRNAs from bacterial and archaeal sequences, whereas rhodophyte cytoplasmic and photosynthetic plastid 5S rRNAs localized closer to the prokaryotic region (Note Fig. 1a). A neighbor-joining tree constructed from pairwise cosine distances in the original high-dimensional representation space showed the same broad organization. Pairwise representation distances were positively correlated with patristic distances in the phylogenetic tree associated with the RF00001 seed alignment (Spearman *ρ* = 0.735; two-sided Mantel permutation-test *P* = 0.001; **Note Fig. 1b**). Relative to a bacterial 5S representation centroid, the rhodophyte cytoplasmic and photosynthetic plastid 5S rRNAs showed substantially higher mean cosine similarity than canonical eukaryotic cytoplasmic 5S rRNAs (0.927 and 0.921 versus 0.676; **Note Fig. 1c**), consistent with their more prokaryotic-like organization in representation space. At the sequence level, the five groups shared highly conserved core positions and a common secondary-structure scaffold, while showing localized nucleotide variation associated with lineage and cellular compartment (**Note Fig. 1d**). RiboGRAM contextual dependency signals varied along the sequence, and partner-ranking signals identified annotated pairing partners across multiple consensus-paired positions. When extended to 576,674 records from the complete RF00001 collection, the broad separation between eukaryotic and prokaryotic 5S rRNAs was retained, with additional organization corresponding to major lineages within each domain (**Note Fig. 1e**). Within this large-scale representation map, a subset of plant-derived records without explicit photosynthetic organellar annotations localized near annotated photosynthetic organellar 5S rRNAs (**Note Fig. 2a**). This region contained 2,734 candidate records, representing 1,246 distinct sequences from 196 plant TaxIDs. Their median cosine similarity to the bacterial 5S representation centroid was 0.938, close to that of annotated photosynthetic organellar 5S rRNAs (0.941) and substantially higher than that of other plant cytoplasmic 5S rRNAs (0.594; **Note Fig. 2b**). We next asked whether this relationship in representation space was also reflected in the underlying 5S rRNA sequences. For each candidate, we compared its maximum sequence similarity to annotated photosynthetic organellar and non-organellar 5S rRNAs. Similarity was consistently higher to the photosynthetic organellar set, with a median difference of 19.1 percentage points (**Note Fig. 2c**). To determine whether the organellar-like 5S signal in these plant records extended to plastid-like genomic context, we quantified the percentage of valid 18-mers exactly matching the corresponding plastid (chloroplast) reference genome within ±0.5-, ±2.5-and ±5-kb windows across 2,721 candidate–control pairs from 196 TaxIDs. Candidate loci showed median plastid-matching proportions of 76.7%, 62.4% and 54.8% across the three scales, respectively, whereas matched nuclear-background controls had median values of 0% at all three scales (**Note Fig. 2d**). Candidate measurements were strongly correlated across window scales (Spearman *ρ* = 0.936, 0.914 and 0.986). The 5S hit was excluded from the ±0.5-kb measurement, whereas the 5S hit plus 100 nt on each side was excluded from the ±2.5-and ±5-kb measurements. Seven initially sampled controls exceeding 5% plastid-reference matching were replaced; the displayed controls therefore provide an outcome-screened descriptive background. The persistence of plastid-matching sequence across increasing window size was consistent with these features extending beyond the 5S locus into the surrounding context. Three representative loci further showed varying extents of plastid-like genomic context (**Note Fig. 2e**). The *Microthlaspi erraticum* candidate showed 98.4% identity across a 10.9-kb global alignment to the *Microthlaspi perfoliatum* plastid reference interval and showed extensive reference-mapped correspondence across the flanking region. The *Lupinus albus* locus showed 71.7% identity across a 13.4-kb global alignment, with partial correspondence of the surrounding gene organization, whereas the *Capsicum annuum* candidate showed a more localized pattern of similarity (41.5% identity across a 16.5-kb global alignment). These global-alignment identities are distinct from the 18-mer matching percentages. Together, these observations showed concordant organellar-like representation and 5S sequence features alongside plastid-like flanking genomic context, without by themselves establishing the physical origin or integration mechanism of individual loci. We then examined whether the sequence constraints learned by RiboGRAM were related to the effects of human 5S rRNA variants on ribosome assembly. The analysis included 206 single-nucleotide variants at 110 positions^13^, with reduced incorporation of mutant 5S rRNA into the 60S ribosomal subunit indicating greater impairment of 5S rRNA function. Without task-specific training, RiboGRAM zero-shot mutation-effect predictions were positively correlated with the experimentally measured functional effects (Spearman *ρ* = 0.273), exceeding RNAfold^14^ ΔMFE (*ρ* = 0.162), Rfam-derived conservation scores (*ρ* = 0.182-0.189) and RNA-MSM^15^, the best-performing alternative RNA language model (*ρ* = 0.244; **Note Fig. 2f**). To test whether closely related natural sequence information could improve the correspondence with experimental variant effects, we used six natural 5S rRNA sequences with high similarity to human 5S rRNA for Low-Rank Adaptation (LoRA)^16^ of RiboGRAM. Following adaptation, the correlation with experimental variant effects increased from *ρ* = 0.273 to *ρ* = 0.339. Mapping the experimental and predicted effects onto the human 80S ribosome structure^17^ (PDB 6QZP) enabled their spatial comparison. Experimentally measured functional impairment was enriched at positions contacting uL5 or uL18, whereas neither the RiboGRAM zero-shot nor the LoRA-adapted zero-shot scores showed corresponding significant enrichment (**Note Fig. 2g**). These results indicate that RiboGRAM captures part of the functional variation in human 5S rRNA, with closely related natural sequences further improving agreement with experimentally measured variant effects. Together, RiboGRAM representations organized sequence variation across taxonomic lineages and cellular compartments within the conserved 5S rRNA scaffold, highlighted plant records with organellar-like 5S sequence features and plastid-like flanking genomic context, and captured part of the experimentally measured functional variation in human 5S rRNA.

**Note Fig. 1.**
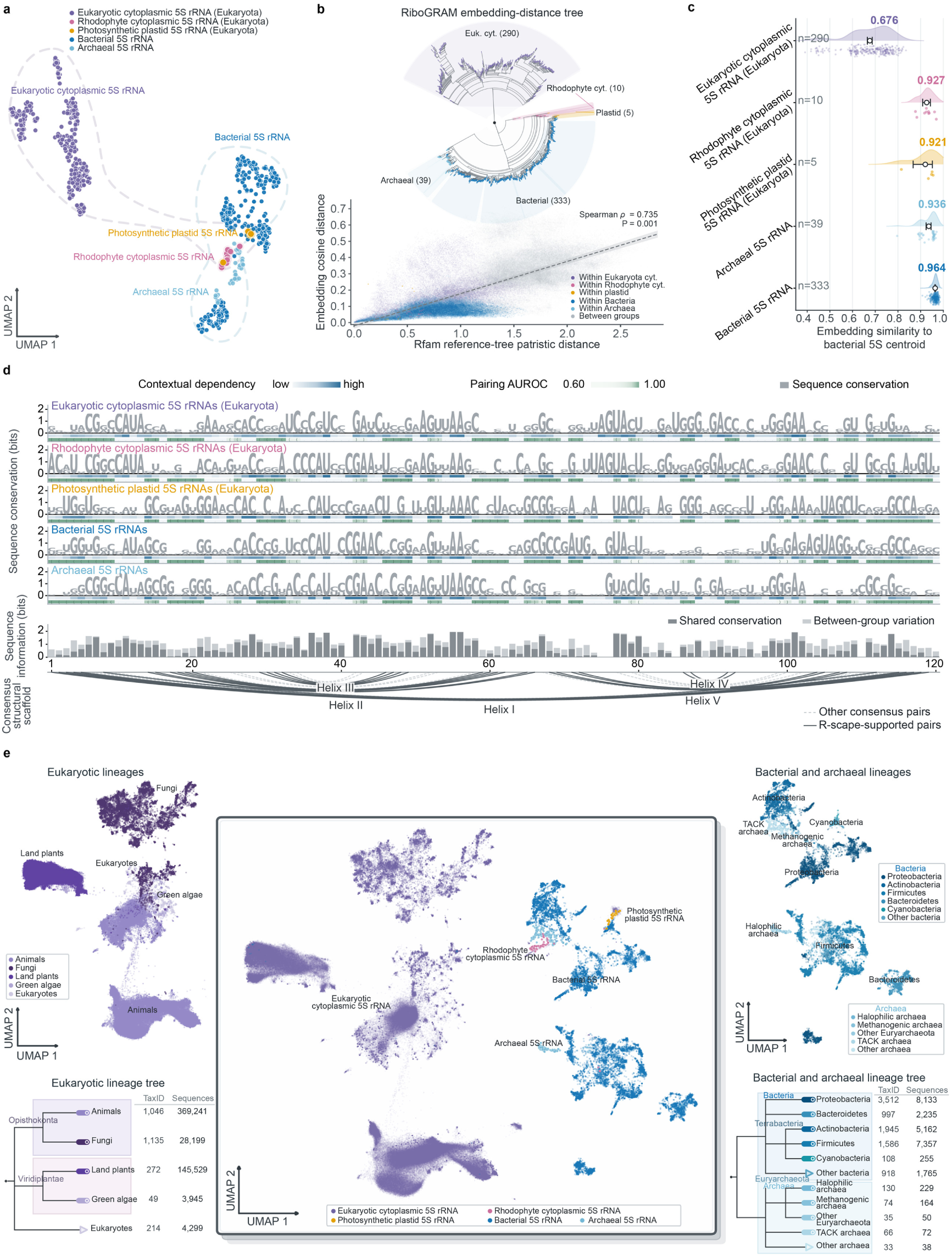
Lineage organization and sequence constraints across the 5S rRNA family. a, UMAP projection of mean-pooled RiboGRAM representations for 677 curated records from the Rfam 5S rRNA seed alignment (RF00001), including canonical eukaryotic cytoplasmic (n = 290), rhodophyte cytoplasmic (n = 10), photosynthetic plastid (n = 5), bacterial (n = 333) and archaeal (n = 39) 5S rRNAs. Colors denote the same five groups. **b**, The neighbor-joining tree (top) was constructed from pairwise cosine distances between the original high-dimensional RiboGRAM representations of the same 677 records shown in panel a. Colored tip markers and semitransparent sectors indicate the five groups. The scatterplot (bottom) compares representation cosine distances for all unique record pairs with their patristic distances in the phylogenetic tree associated with the Rfam 5S rRNA seed alignment (RF00001). The two distance measures showed a significant positive correlation (Spearman’s *ρ* = 0.735), supported by a two-sided Mantel permutation test (*P*-value = 0.001). The dashed line indicates the least-squares fit, and the shading represents the 95% confidence interval; colored and gray points denote within-group and between-group comparisons, respectively. **c,** Cosine similarity between the RiboGRAM representation of each 5S rRNA and a bacterial 5S rRNA reference centroid for the five groups shown in panel a. Points denote individual sequences, while open circles indicate the means. Thick and thin lines denote the 50% and 95% bootstrap intervals, respectively. **d,** Sequence conservation, contextual dependency and pairing signals across Rfam 5S rRNA seed alignment (RF00001) positions 1–120 for 574 representative sequences drawn from the five groups shown in panel a (one sequence per group–TaxID combination). Gray sequence logos show nucleotide composition and conservation within each group, with total logo height representing corrected nucleotide information at each position. Blue tracks show group-median RiboGRAM contextual dependency scores based on masked-nucleotide recovery, and teal tracks show partner-ranking AUROC at consensus-paired positions. The lower bar tracks distinguish sequence information shared across groups from lineage-associated nucleotide variation. Arcs denote the 34 consensus base pairs in the Rfam RF00001 secondary-structure annotation. Dark solid arcs indicate the 22 pairs with significant R-scape18 covariation support (*E* ≤ 0.05), whereas pale dashed arcs indicate the remaining annotated pairs. Helices I–V denote the five canonical helices of 5S rRNA. **e,** UMAP projections of mean-pooled RiboGRAM representations for 576,674 records from the complete RF00001 collection, including canonical eukaryotic cytoplasmic (n = 550,832), rhodophyte cytoplasmic (n = 287), photosynthetic plastid (n = 96), bacterial (n = 24,907), and archaeal (n = 553) 5S rRNAs. The central view shows all records, whereas the flanking views use the same UMAP coordinates to display the major eukaryotic lineages on the left and the major bacterial and archaeal lineages on the right. The diagrams below provide rank-based NCBI taxonomic summaries of 2,716 eukaryotic and 9,404 bacterial or archaeal TaxIDs.

**Note Fig. 2.**
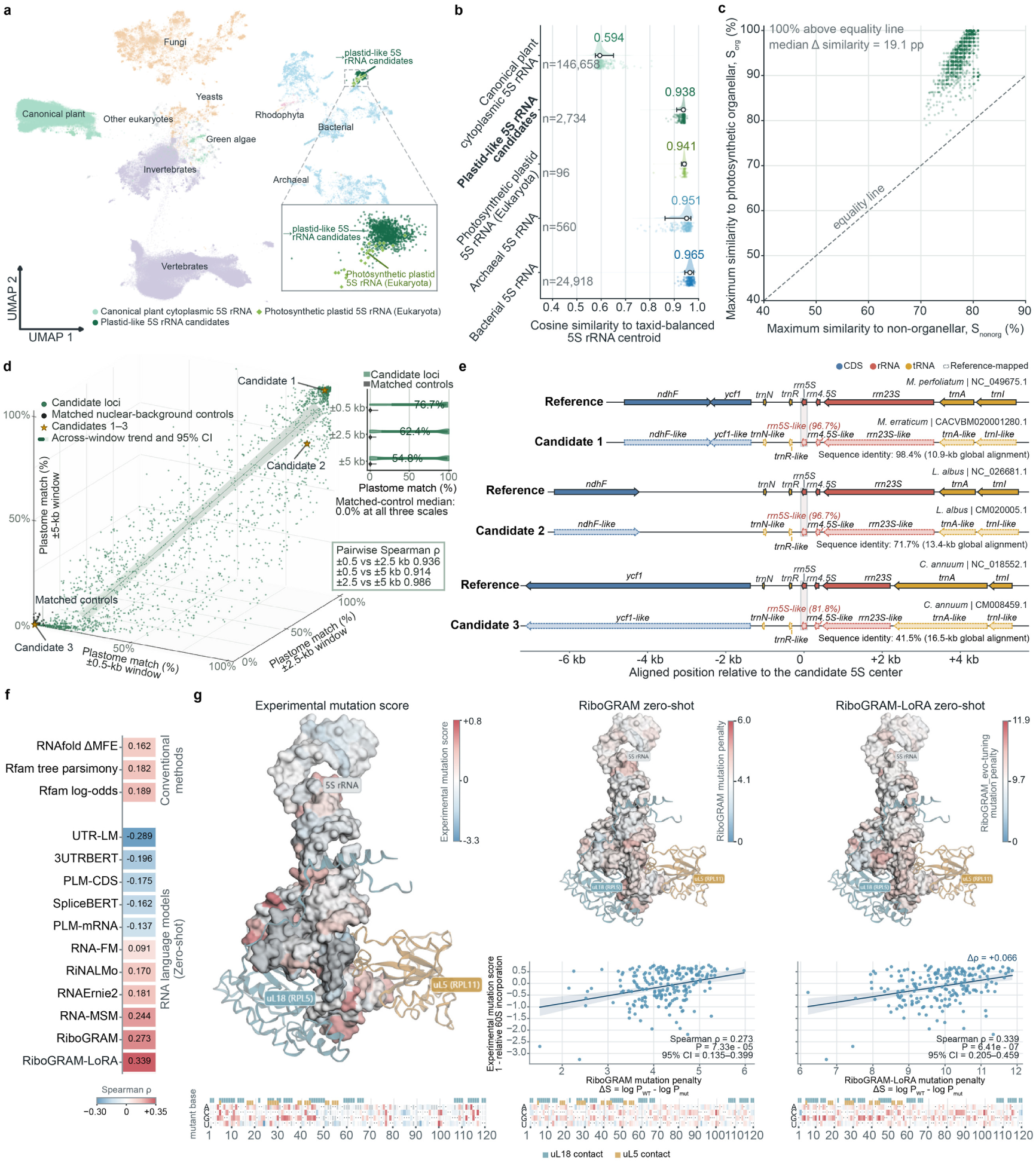
Photosynthetic organellar 5S-like candidates and human 5S variant effects. a, UMAP of mean-pooled RiboGRAM representations for the complete Rfam 5S rRNA family (RF00001), highlighting canonical plant cytoplasmic 5S rRNAs (n = 146,658), photosynthetic organellar 5S-like candidate records (n = 2,734; 1,246 distinct sequences), and explicitly annotated photosynthetic organellar 5S rRNA records (n = 96). The inset enlarges the region containing candidate and annotated photosynthetic organellar 5S rRNAs. **b,** Cosine similarity of the five indicated 5S rRNA groups to the bacterial 5S representation centroid. Half-violin plots and points show the group distributions. Open circles, thick lines, and thin lines indicate the median, interquartile range, and 5th–95th percentile range, respectively. Labelled values indicate group medians. **c,** Sequence similarity of each of the 2,734 candidate records to non-organellar and annotated photosynthetic organellar 5S rRNAs. The x and y axes show the maximum similarity to 672 non-organellar sequences from RF00001 seed alignment and to 96 explicitly annotated photosynthetic organellar 5S rRNAs, respectively. The dashed diagonal line denotes equal similarity to the two reference sets. **d,** Multiscale plastid-reference matching. For 2,721 complete candidate–control pairs from 196 TaxIDs, the percentages of valid 18-mers exactly matching the corresponding plastid (chloroplast) reference genome were quantified within ±0.5-, ±2.5-, and ±5-kb windows centered on each locus. Candidate and control distributions are shown at each scale, with open circles indicating medians and horizontal lines indicating interquartile ranges. The fitted line and shaded band denote the candidate-only three-dimensional orthogonal fit and its 95% confidence region, and the inset reports pairwise Spearman correlations across window scales. Stars identify the three representative candidate loci shown shown in panel e. **e,** Representative plastid-like flanking genomic contexts. The tracks in panel e compare the three representative candidate loci with their corresponding plastid reference intervals. Solid features denote genes annotated in the reference genomes, and pale dashed features denote their reference-mapped counterparts in the candidate regions. The examples comprise *Microthlaspi erraticum* CACVBM020001280.1 (98.4% identity across a 10.9-kb global alignment to the *M. perfoliatum* chloroplast reference NC_049675.1), *Lupinus albus* CM020005.1 (71.7% across 13.4 kb to NC_026681.1) and *Capsicum annuum* CM008459.1 (41.5% across 16.5 kb to NC_018552.1). **f,** Spearman correlations between predicted human 5S rRNA variant effects and experimentally measured effects on incorporation of mutant 5S rRNA into the 60S ribosomal subunit for 206 single-nucleotide variants at 110 positions13. Experimental effects are reported as 1-relative 60S-subunit incorporation. Conventional baselines comprise RNAfold ΔMFE and the Rfam-derived tree-parsimony and allele-log-odds scores. RNA language models were evaluated using zero-shot mutation scoring. **g,** Experimental and predicted human 5S rRNA variant effects mapped onto the human 80S ribosome structure (PDB 6QZP). Columns show the experimental mutation effect, the RiboGRAM zero-shot mutation penalty and the LoRA-adapted RiboGRAM (“evo-tuning”) mutation penalty, with ΔS = log P(WT | context) − log P(mutant | context). Structures are colored by the median score across available substitutions at each position. Scatter plots compare predicted and experimental effects across the 206 variants, with lines and shaded regions indicating linear fits and 95% confidence intervals, respectively. Heatmaps show substitution-specific effects across 5S rRNA positions 1–120. Protein-contacting nucleotides were defined by a minimum heavy-atom distance of ≤4 Å to uL5 or uL18.

**Supplementary Note 2. GC composition and RNA-class context stratify zero-shot base-pair recovery.** Zero-shot base-pair recovery varied substantially among individual RNAs, raising the question of whether this variation was related to sequence composition and RNA-class context. Across the bpRNA-1m TR0 dataset, per-sequence recovery showed a nonlinear relationship with global GC fraction. RNA structure predictors and ncRNA-rich language models generally reached their highest mean F1 at GC fractions of 0.57-0.59, with lower recovery toward more compositionally extreme sequences (**Note Fig. 3a**). To examine local compositional variation beyond global GC fraction, we quantified GC-profile ruggedness from an 11-nt-smoothed local GC/AU profile, with higher values indicating greater local heterogeneity. Across 10,814 TR0 sequences, GC-profile ruggedness showed a modest positive association with RiboGRAM base-pair recovery (Spearman *ρ* = 0.185; **Note Fig. 3b**).

**Note Fig. 3.**
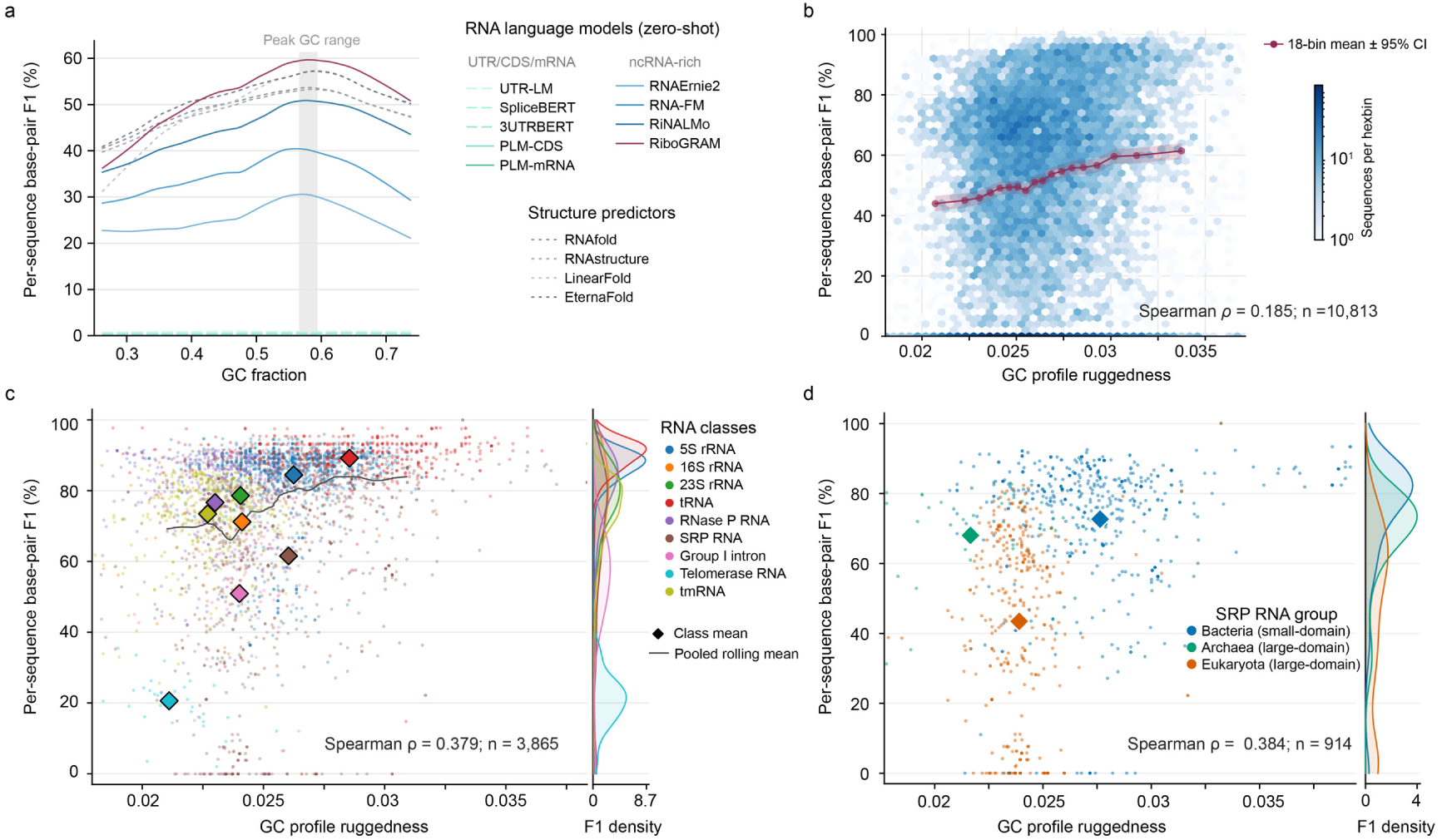
Zero-shot base-pair recovery varies with global GC composition and local GC heterogeneity. a, Per-sequence base-pair F1 across GC-fraction bins for 10,814 bpRNA-1m TR0 sequences evaluated by all methods. Lines show bin-wise means. The shaded interval marks the GC-fraction range associated with peak performance across RNA structure predictors and ncRNA-rich language models. **b,** Relationship between GC-profile ruggedness and RiboGRAM per-sequence base-pair F1 across 10,814 TR0 sequences. GC-profile ruggedness was calculated from the 11-nt-smoothed local GC/AU composition profile as the mean absolute difference between adjacent positions. Hexagons show sequence density, and the overlaid points and line show mean F1 across 18 equal-count bins with 95% bootstrap confidence intervals. **c,** GC-profile ruggedness and RiboGRAM per-sequence base-pair F1 across 3,865 ArchiveII RNAs from nine RNA classes. Points denote individual RNAs, diamonds indicate class means and the black line shows the pooled rolling mean. Marginal density curves show the class-specific F1 distributions. **d,** GC-profile ruggedness and RiboGRAM per-sequence base-pair F1 across 914 signal recognition particle (SRP) RNAs, comprising bacterial small-domain (n = 542), archaeal large-domain (n = 39) and eukaryotic large-domain (n = 333) SRP groups. Points denote individual RNAs, diamonds indicate group means and marginal density curves show the corresponding F1 distributions. Spearman correlations were calculated across individual RNAs in panels b-d. Recovery also varied across RNA classes. Among 3,865 ArchiveII RNAs from nine classes, GC-profile ruggedness and RiboGRAM F1 were positively associated across the pooled dataset (Spearman *ρ* = 0.379; **Note Fig. 3c**). The distributions differed substantially among classes. tRNA and 5S rRNA showed relatively high recovery, whereas group I introns and telomerase RNAs showed lower recovery. Thus, the pooled relationship with GC ruggedness occurred alongside broader class-specific differences in sequence and structural organization. Signal recognition particle (SRP) RNAs provide a more focused comparison within a single RNA class. Bacterial small-domain, archaeal large-domain and eukaryotic large-domain SRPs occupied distinct regions of the GC ruggedness–F1 space, with mean F1 values of 0.73, 0.68 and 0.43, respectively (**Note Fig. 3d**). Across the 914 SRP RNAs, GC-profile ruggedness remained positively associated with recovery (Spearman *ρ* = 0.384). The difference between archaeal and eukaryotic SRPs, despite both belonging to the large-domain group, indicates that the observed variation is not explained by small-versus large-domain architecture alone. Together, these analyses show that zero-shot base-pair recovery varies with both global GC composition and local GC heterogeneity, with additional stratification across RNA classes and SRP groups.

**Supplementary Note 3. RNA-class context and pseudoknot topology stratify zero-shot base-pair recovery.** Zero-shot base-pair recovery varied across nine ArchiveII RNA classes spanning broad differences in sequence length and base-pairing architecture. Across the same 3,865 RNAs, the per-sequence recovery distributions of RiboGRAM and the structure-supervised RNet-SS^6, 7^ varied across RNA classes (**Note Fig. 4a**). When class-level micro-F1 values were averaged equally across the nine classes, RiboGRAM and RNet-SS reached 0.689 and 0.681, respectively (**Supplementary** Fig. 9a). The largest class-level differences favoring RiboGRAM occurred for transfer-messenger RNA (tmRNA) (micro-F1, 0.75 versus 0.57) and RNase P RNA (0.801 versus 0.67), whereas RNet-SS showed higher recovery for group I introns and telomerase RNA. tmRNA and RNase P RNA contained the largest numbers of pseudoknot-associated reference pairs in ArchiveII, defined as base pairs participating in at least one crossing interaction (**Note Fig. 4b**). We therefore examined recovery of these pairs separately from overall base-pair recovery. Standard nested secondary structure predictors generated no crossing-pair predictions. EternaFold+TK, which permits crossing pairs through ThreshKnot decoding, reached micro-F1 values of 4.6% and 4.5% for tmRNA and RNase P RNA, respectively, whereas the structure-supervised RNet-SS reached 39.7% and 34.3%. Among language models, those pretrained on UTR/CDS/mRNA remained near zero (micro-F1 ≤ 0.4%), whereas recovery was substantially higher among the ncRNA-rich models, with RiboGRAM reaching 75.9% and 78.2% for tmRNA and RNase P RNA, respectively (**Note Fig. 4c**). These comparisons distinguish the recovery of crossing interactions from the broader class-level variation in overall base-pair recovery. Structure-level comparisons across the nine RNA classes provide a base-pair-level view of these prediction patterns (**Note Fig. 4d**). In the tRNA, 5S rRNA and SRP RNA structures, which contained no pseudoknot-associated reference pairs, both RiboGRAM and EternaFold recovered most reference pairs, with differences arising mainly from missed or additional predictions. In the RNase P RNA and tmRNA structures, RiboGRAM recovered 18 of 19 and 62 of 63 pseudoknot-associated reference pairs, respectively, with overall per-sequence F1 values of 0.93 and 0.87, compared with 0.71 and 0.56 for RNet-SS. Crossing interactions were also recovered in the 16S rRNA and group I intron structures. The telomerase RNA structure showed a different prediction profile. RiboGRAM recovered fewer reference pairs with relatively few additional predictions (precision = 0.96, recall = 0.22), whereas RNet-SS recovered more reference pairs at lower precision (precision = 0.46, recall = 0.53). Together, these structure-level comparisons indicate that class-level variation in zero-shot recovery is accompanied by differences in recovery completeness, prediction specificity and crossing-interaction recovery across structural contexts.

**Note Fig. 4.**
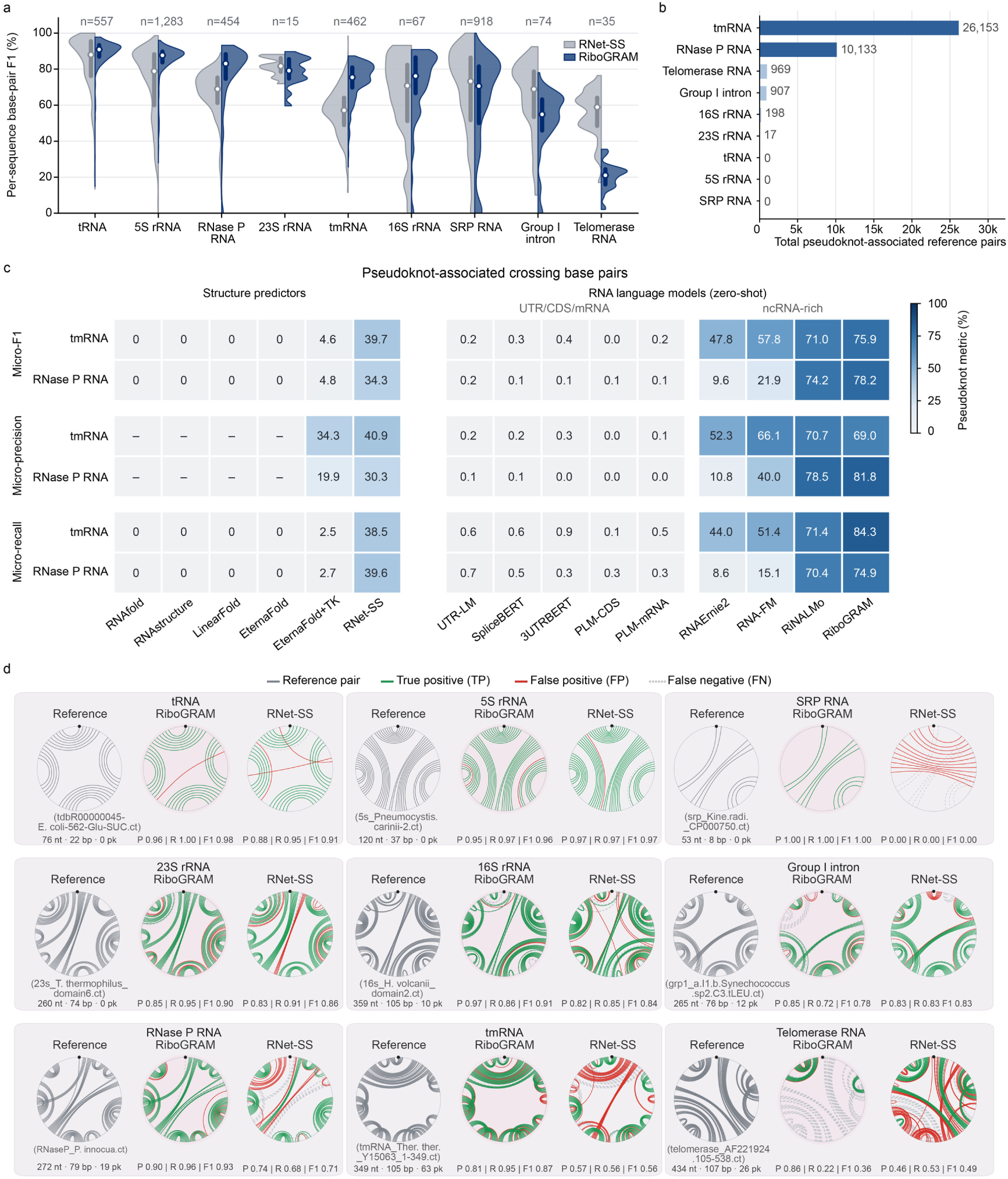
Base-pair recovery across RNA classes and recovery of pseudoknot-associated pairs. a, Per-sequence base-pair recovery across 3,865 ArchiveII RNAs from nine RNA classes. RiboGRAM and RNet-SS were evaluated on the same sequences within each class, with F1 calculated from all reference and predicted base pairs for each RNA. White circles and vertical bars indicate medians and interquartile ranges, respectively; n denotes the number of sequences in each class. **b,** Pseudoknot-associated reference base-pair counts across the nine ArchiveII RNA classes, pooled across sequences within each class. Pseudoknot-associated pairs were defined as base pairs participating in at least one crossing interaction. **c,** Pseudoknot-specific base-pair recovery in tmRNA and RNase P RNA, the two classes contributing the largest total numbers of pseudoknot-associated reference pairs. Micro-F1, micro-precision, and micro-recall were calculated by pooling pair-level true-positive, false-positive and false-negative counts across sequences within each class. Reference and predicted pseudoknot-associated pairs were identified independently using the criterion defined in panel b. En dashes indicate undefined precision when no pseudoknot-associated pairs were predicted. **d,** Reference and predicted base-pairing patterns for selected ArchiveII RNAs spanning the nine RNA classes. The circular diagrams show the reference structures alongside RiboGRAM and RNet-SS predictions. The black points mark the 5’ end. Labels below the reference structures give the original ArchiveII source identifier, sequence length, total number of reference base pairs and number of pseudoknot-associated reference pairs. Labels below the predictions report per-sequence precision, recall and F1. nt, nucleotides; bp, reference base pairs; pk, pseudoknot-associated reference base pairs; TP, true positive; FP, false positive; FN, false negative.

**Supplementary Note 4. RNA–RNA interaction classes differ in predictive performance and sequence-pair properties.** RNA–RNA interactions span diverse RNA classes and pairwise sequence contexts, providing a complementary setting for examining how sequence organization relates to interaction prediction. We therefore examined two RNA–RNA interaction (RRI) benchmarks with different interaction scopes and control designs. In the MirTarRAW miRNA–mRNA benchmark, RiboGRAM–RRI reached an AUC of 0.970 and accuracy of 0.913 on the 22,176-pair test set (**Note Fig. 5a**). Across all 1,024 miRNA 5-mers, positive and negative pairs showed marked frequency differences, with different 5-mers enriched in the two groups (**Note Fig. 5b**). The DeepMirTar^19^ component of MirTarRAW generates negative pairs by shuffling mature miRNA sequences^20^, which contributes to these label-associated compositional differences. MirTarRAW therefore represents a classification setting in which interaction-related sequence information is accompanied by benchmark-specific differences in miRNA sequence composition. The RISE-derived benchmark extended the analyses across 15 RRI classes under a locally matched control design. Experimentally supported interactions were treated as positive pairs, where matched controls were generated by replacing one interaction partner with an equal-length, same-strand flanking sequence from the same chromosome, separated from the original region by 500 nt. After 80% sequence identity filtering, the test set contained 28,551 RNA pairs. Predictive performance varied across interaction classes and models (**Note Fig. 5c**). RiboGRAM-RRI had the highest AUC in 11 of the 15 classes (mean AUC = 0.735) and the highest accuracy in 14 (mean accuracy = 0.786), with both metrics varying across classes. We next examined how sequence-pair organization differed between positive pairs and matched controls in the RISE-derived training set. Reverse-complement k-mer sharing was higher in positive pairs across all 15 interaction classes at k-mer lengths of 3–6 nt (**Note Fig. 5d**), although the magnitude of the difference varied among classes, with larger shifts in several rRNA-and snRNA-containing interactions. GC-content similarity showed the same overall direction across the 15 classes, with particularly large differences in rRNA–rRNA, snRNA– snRNA and ncRNA–ncRNA pairs (**Note Fig. 5e**). Reverse-complement k-mer sharing quantifies local complementary potential, whereas GC-content similarity measures global compositional matching between RNA partners. Low-complexity and repetitive sequence content showed a more class-dependent pattern. Positive–control differences in the low-complexity/repeat score varied in both magnitude and direction across interaction classes (**Note Fig. 5f)**. Higher scores in positive pairs were most evident for several mRNA-, lncRNA-and pseudogene-containing classes, whereas rRNA-and snRNA-associated classes generally showed lower scores than their matched controls. This pattern contrasted with the broadly positive shifts in reverse-complement k-mer sharing and GC-content similarity. Within the RISE-derived dataset, pairwise sequence organization varied among interaction classes in multiple dimensions, including local complementarity potential, global GC-content matching and low-complexity and repetitive sequence content.

**Note Fig. 5.**
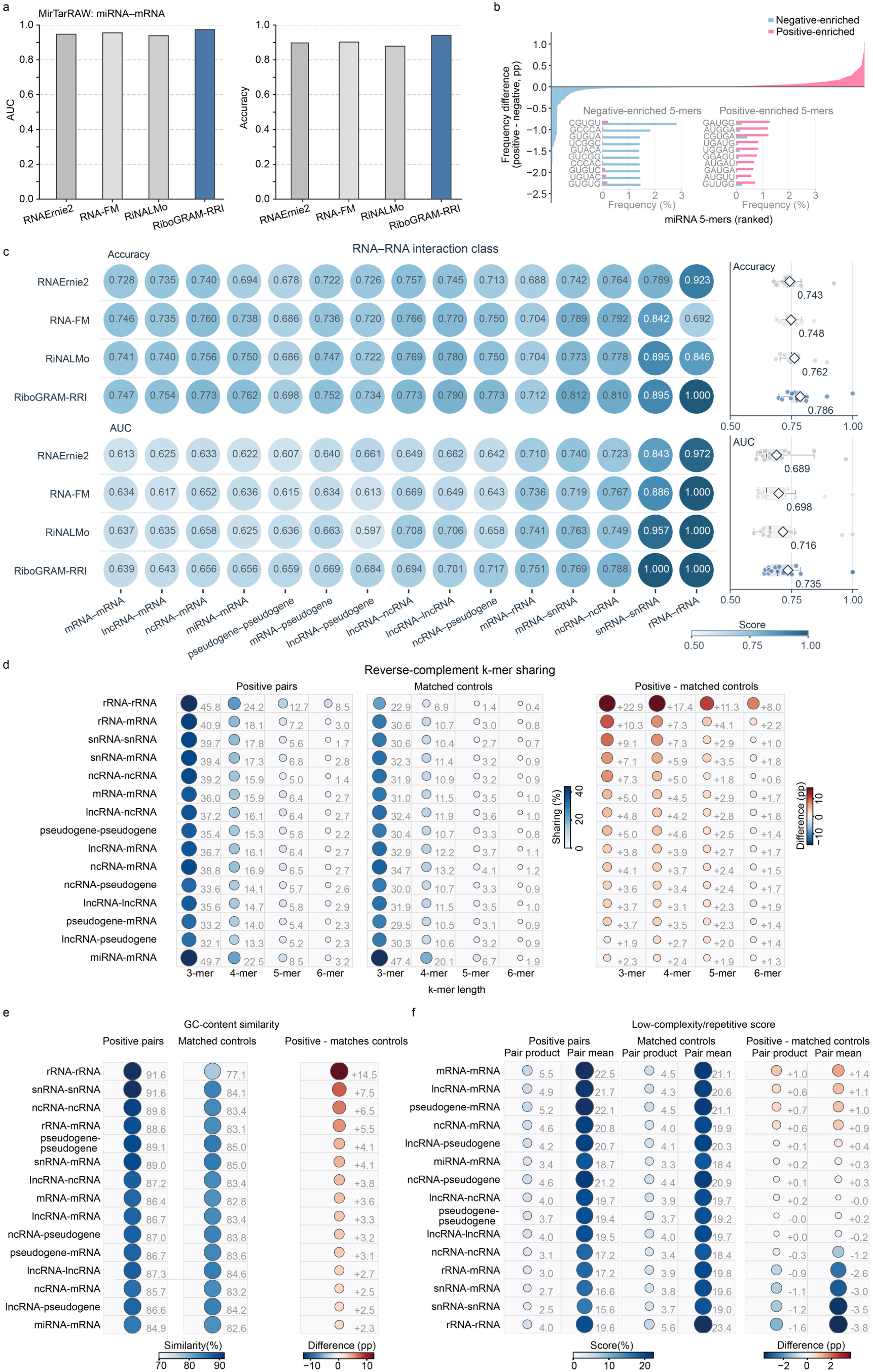
RNA–RNA interaction prediction varies across interaction classes alongside class-dependent sequence-pair properties. a, RNA–RNA interaction prediction on the MirTarRAW miRNA–mRNA benchmark. AUC and accuracy are shown for the indicated RNA language models on the held-out test set comprising 22,176 RNA pairs. **b,** Label-associated miRNA sequence composition in MirTarRAW. The ranked profile shows positive-minus-negative frequency differences across all 1,024 miRNA 5-mers, and insets show the positive-minus-negative frequencies of the ten 5-mers with the largest differences in each direction. **c,** Prediction performance across 15 RNA–RNA interaction classes in the RISE-derived benchmark. Accuracy and AUC are shown for the 28,551 interaction pairs retained in the test set after 80% sequence-identity filtering. Right, the distribution of performance across interaction classes. Points denote individual classes. Boxes indicate medians and interquartile ranges, and diamonds indicate means. **d,** Local reverse-complement sequence sharing between RNA partners across the 15 interaction classes. Sharing percentage was quantified for 3–6-mers as the fraction of k-mer occurrences shared between one RNA and the reverse complement of its partner, normalized by the smaller k-mer count of the two RNAs. **e,** Global GC-content matching between RNA partners across interaction classes. GC-content similarity was defined as (1-|GC_1-GC_2|), with higher values indicating more closely matched GC composition between the two RNAs. **f,** Low-complexity and repetitive sequence content across interaction classes. Each RNA was assigned a low-complexity/repeat score integrating nucleotide-composition complexity, homopolymer content and tandem-repeat coverage. RNA-pair values were summarized by the product and mean of the two RNA-level scores. For panel d-f, positive pairs were compared with matched controls in the RISE-derived training set (n = 261,997 pairs), and differences denote positive-pair minus matched-control values.

**Supplementary Note 5. RNA–chromatin interaction prediction across experimental contexts and transcript-associated representations.** RNA–chromatin interaction maps comprise heterogeneous assay-defined associations involving diverse RNA species and genomic contexts and do not distinguish direct RNA–DNA contacts from associations mediated by proteins or other chromatin components. Within this assay-defined setting, we evaluated RNA–chromatin interaction (RCI) prediction across experimental contexts and examined how task-specific post-training reorganized RNA– chromatin pair representations. On the HEK293T test set^21^, the resulting RiboGRAM–RCI model showed an AUC of 0.951 and an accuracy of 0.890, compared with 0.911 and 0.831 for DeepRCI^22^, respectively (**Note Fig. 6a**). The same RiboGRAM-RCI model, without cell-line-specific retraining, was then evaluated on five external datasets from HUVEC, fibroblasts, K562, MDA-MB-231 and MM.1S cells, derived from iMARGI^23, 24^, Red-C^25^ and GRID-seq^26^ studies. Across these external contexts, RiboGRAM-RCI and DeepRCI^22^ exhibited mean AUCs of 0.692 and 0.655 and mean accuracies of 0.620 and 0.604, respectively, with performance varying among datasets. We next examined how RCI-specific post-training altered the organization of RNA–chromatin pair representations. UMAP projections of the same 200,000 HEK293T test pairs showed extensive overlap between detected contacts and matched controls before post-training, whereas the RiboGRAM–RCI representation showed clearer separation between the two groups (**Note Fig. 6b**). Within the RiboGRAM-RCI representation, RNA–DNA complementary-tract length was distributed non-uniformly among the 100,000 detected contact pairs (**Note Fig. 6c**). Pairs containing longer contiguous tracts with potential complementarity were concentrated within a localized region of the projection. This spatial concentration links sequence complementarity potential to one dimension of organization within the post-trained representation. In this setting, complementary-tract length is interpreted as a sequence-level property of assay-defined RNA– chromatin contacts rather than direct evidence of RNA–DNA base pairing. RNA annotation provided a complementary view of the same representation. Annotation classes occupied non-uniform regions of the projection, including a region enriched for lncRNA-associated contacts (**Note Fig. 6d**). At the transcript level, this region contained 529 detected contact pairs from 17 genes, with XIST accounting for 374 (70.7%; **Note Fig. 6e**)27. The lncRNA-enriched region was therefore concentrated in a limited set of transcripts rather than being broadly distributed across lncRNAs. Overall, RNA–chromatin interaction prediction varied across experimental contexts, and the post-trained representation was associated with complementarity potential, RNA annotation class and transcript identity.

**Note Fig. 6.**
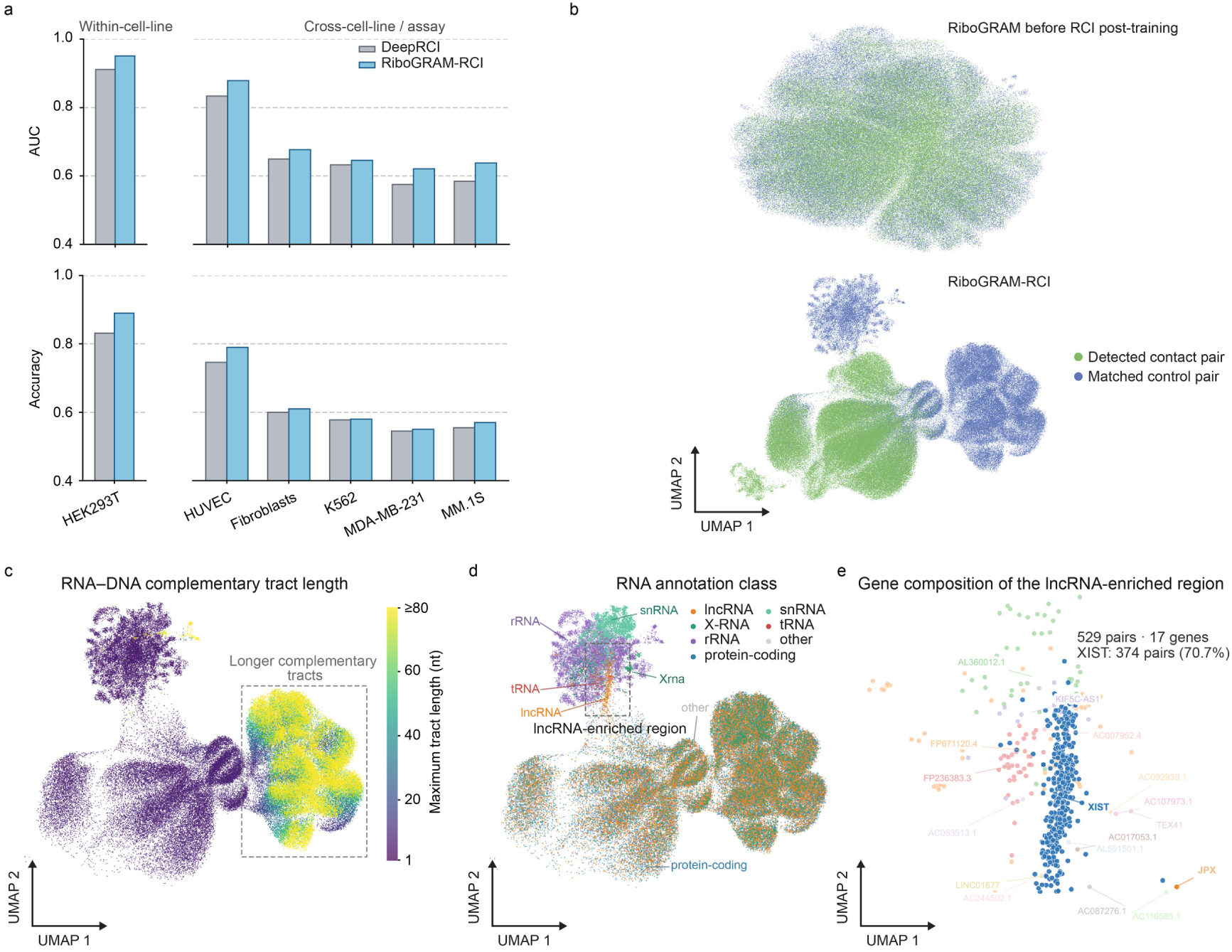
RNA–chromatin interaction prediction varies across experimental contexts with sequence-and transcript-associated representation patterns. a, RNA–chromatin interaction prediction within HEK293T and across five external cell-line datasets. AUC and accuracy compare RiboGRAM–RCI with DeepRCI on the HEK293T test set and on HUVEC, fibroblasts, K562, MDA-MB-231 and MM.1S datasets, reported in iMARGI23, 24, Red-C25 and GRID-seq26 studies. RiboGRAM-RCI was applied to the external datasets without cell-line-specific retraining. **b,** Organization of HEK293T RNA–chromatin pair representations before and after RCI-specific post-training. The same 200,000 test pairs are shown in both projections, comprising 100,000 detected contact pairs and 100,000 matched control pairs. **c,** RNA–DNA complementary-tract length across the RiboGRAM–RCI representation of the 100,000 detected contact pairs in panel b. Each pair is colored by the longest contiguous tract with potential complementarity between the RNA and either DNA strand. The boxed region contains pairs with longer complementary tracts. **d,** RNA annotation classes across the same RiboGRAM–RCI representation. The dashed box marks a region enriched for lncRNA-associated contacts and expanded in panel e. X-RNA denotes RNA-Chrom-defined transcript clusters assembled from previously unannotated RNA segments. **e,** Gene composition of the lncRNA-enriched region boxed in panel d. The region contains 529 detected contact pairs from 17 genes, of which XIST accounts for 374 (70.7%)^27^.

## Notes

https://zenodo.org/records/23201637

https://www.ultrarnalab.com/ribogram.php

https://codeocean.com/capsule/2602506/tree/v1

https://www.ultrarnalab.com/ribogram_usage.php

### Competing Interest Statement

The authors have declared no competing interest.

